# Genome-wide association study of open field behavior in outbred heterogeneous stock rats identifies multiple loci implicated in psychiatric disorders

**DOI:** 10.1101/2021.10.18.464872

**Authors:** Mustafa Hakan Gunturkun, Tengfei Wang, Apurva S. Chitre, Angel Garcia Martinez, Katie Holl, Celine St. Pierre, Hannah Bimschleger, Jianjun Gao, Riyan Cheng, Oksana Polesskaya, Leah C. Solberg-Woods, Abraham A. Palmer, Hao Chen

## Abstract

Many personality traits are influenced by genetic factors. Rodents models provide an efficient system for analyzing genetic contribution to these traits. Using 1,246 adolescent heterogeneous stock (HS) male and female rats, we conducted a genome-wide association study (GWAS) of behaviors measured in an open field, including locomotion, novel object interaction, and social interaction. We identified 30 genome-wide significant quantitative trait loci (QTL). Using multiple criteria, including the presence of high impact genomic variants and co-localization of cis-eQTL, we identified 13 candidate genes (*Adarb2, Ankrd26, Cacna1c, Clock, Crhr1, Ctu2, Cyp26b1, Eva1a, Fam114a1, Kcnj9, Mlf2, Rab27b, Sec11a*) for these traits. Most of these genes have been implicated by human GWAS of various psychiatric traits. For example, *Cacna1c*, a gene known to be critical for social behavior in rodents and implicated in human schizophrenia and bipolar disorder, is a candidate gene for distance to the social zone. In addition, the QTL region for total distance to the novel object zone, on *Chr1* at 144 Mb, is syntenic to a hotspot on human Chr15 (82.5-90.8 Mb) that contains 14 genes associated with psychiatric or substance abuse traits. Although some of the genes identified by this study appear to replicate findings from prior human GWAS, others likely represent novel findings that can be the catalyst for future molecular and genetic insights into human psychiatric diseases. Together, these findings provide strong support for the use of the HS population to study psychiatric disorders.

## 1 Introduction

Many personality traits are predictors of vulnerability to addiction [1]. For example, individuals with symptoms of anxiety are more likely to be smokers [2, 3], and novelty seeking is positively correlated with both smoking onset [4] and cocaine abuse [5]. In addition, the social environment plays a critical role in the development and treatment of addiction [6]. Many of these phenomena can be modeled using rodents to unveil their neural, genetic, and molecular mechanisms [7, 8, 9, 10].

The open-field test (OFT) is a widely used behavioral test for measuring anxiety-like and exploratory behavior in rodents [11, 12, 13, 14]. A rodent is typically placed in an open chamber surrounded by tall walls. Video recording of the rodent’s locomotor movements is then analyzed. In general, rats spend most of the testing session walking along the wall (i.e. thigmotaxis). Increased time spent in the center of the area or decreased latency to enter the center are interpreted as indications of lower anxiety. The OFT is widely used to model anxiety and is sensitive to the anxiolytic-like effects of classical benzodiazepines, and 5-HT1A receptor agonists [11]. The novel object interaction test (NOIT) is usually conducted in an open arena where a novel object is placed in the center. The time spent and distance traveled around the object zone are used as indicators of preference for novelty. Novel object interaction has been considered as an important predictor in addiction-like traits [15, 16] and high novelty preference increases the propensity for addictive drug-seeking behavior [17, 18, 9]. There are multiple different methods for conducting social interaction test (SIT) in rats [19, 20, 21]. In general, an unfamiliar stimulus rat and the rats to be tested are placed in the same arena. While manual scoring of social interaction often allows both rats to be freely moving, experiments using automated video analysis often limit the movement of the stimulus rat and calculate the time spend and distance traveled by the test rat around the stimulus rat.

The heterogeneous stock (HS) rats were originally derived from interbreeding eight inbred strains [22] and have been maintained as outbred for more than 90 generations. HS rats have been successfully used in several high-resolution genome-wide association studies (GWAS) [23, 24, 25, 26]. Here we report the results on associations of genomic loci with measures obtained from OFT, NOIT and SIT. These analyses were based on an expanded data set that contained about twice the sample size of that reported previously [27]. These data were collected as part of a larger GWAS on socially acquired nicotine intravenous self-administration, which will be the subject of a separate publication.

## 2 Materials and Methods

### 2.1 Animals

The N/NIH heterogeneous stock (HS) rat (RRID:RGD2314009), was created at the NIH in 1984 by interbreeding the following eight inbred founder strains: ACI/N, BN/SsN, BUF/N, F344/N, M520/N, MR/N, WKY/N and WN/N [22]. The HS rats used in this study were sent from The Medical College of Wisconsin to the University of Tennessee Health Science Center (UTHSC) at 3–6 weeks of age. A total of 16 batches of HS rats were transferred between October 27, 2014 and September 20, 2018. Each batch consisted of 25 males and 25 females that were used as breeders. After a two-week quarantine period, rats were transferred to a reversed 12h light-dark cycle (lights off at 9:00 AM) housing room. Breeding pairs were assigned according to an algorithm that maximized the genetic diversity of the offspring. Litters were culled to a maximum of 8 pups to ensure a consistent nutritional environment. Rats were weaned on PND 21. An RFID was inserted subcutaneously when rats were weaned. Two male and two female rats per litter were used for behavioral studies. Teklad Irradiated LM-485 Mouse/Rat Diet and water were provided *ad libitum*. All rats were group-housed with 2-4 same-sex peers throughout the experiments to avoid social isolation. All procedures were conducted in accordance with the NIH Guidelines concerning the Care and Use of Laboratory Animals, as approved by the Institutional Animal Care and Use Committee of the University of Tennessee Health Science Center.

### 2.2 Study Design

All HS rats (626 males and 620 females in total from 16 batches) were adolescents when tests began. Their age was 31.8 *±* 2.6 (mean *±* STD) on the day of the OFT. Each HS rat was tested in all three behavioral tests, one test per day, in the following sequence: OFT, NOIT, and SIT. All tests were conducted in the dark phase of the light cycle (9 AM – 4 PM) and were conducted in the same open field and recorded using the same video capture system.

### 2.3 Behavioral testing procedure

#### 2.3.1 Open field test

Two OFT arenas were constructed using black acrylic glass, measuring 100*cm*(*L*)*×* 100*cm*(*W*)*×*50*cm*(*H*), which were placed side by side. The floors were covered by wood boards painted with either black or white acrylic paint (ART-Alternatives, ASTM D-4236, Emeryville, CA, USA) to contrast the coat of the animals (i.e. a black board was used for rats with white fur). The test chambers were illuminated by a long-range, 850-nm infrared light (LIR850-70, LDP LLC, Carlstadt, NJ) located 160 cm above the center of the two test chambers. No source of visible light was present during behavioral testing, with the exception of a flat panel monitor (Dell 1908FP). A digital camera (Panasonic WV-BP334) fitted with an 830 nm infrared filter (X-Nite830-M37, LTP LLC, Carlstadt, NJ) and located next to the infrared light source was used to record the behavior of the rats. All rats were released at the same corner of the test chamber, and data were collected for 1 h.

#### 2.3.2 Novel object interaction test

This test was conducted the day after the OFT in the same arena. A cylindrical rat cage constructed using 24 aluminum rods (30 cm in length) spaced 1.7 cm apart was used as the novel object. The bottom and top of the cage (15 cm in diameter) were manufactured using a 3D printer from polylactic acid. The design can be downloaded from https://github.com/chen42/RatSocialInteractionTest. The novel object was placed in the center of the arena before testing. The test duration was 20 min and was recorded using the same camera as that used in the OFT.

#### 2.3.3 Social interaction test

This test was conducted the day after the NOIT. This test compares the preference of a subject rat for a stimulus rat restricted in a cylindrical cage (i.e. the novel object used in the NOIT) against an empty cylindrical cage. The test arena was reduced to 100*cm*(*L*) *×* 60*cm*(*W*) *×* 50*cm*(*H*) by using a black board placed vertically in the arena. Two cylindrical cages described above were placed ∼30 cm away from the walls on opposite sides (i.e., similar to the arrangement commonly used in the three-chamber test). A randomly selected stimulus Sprague-Dawley rat of the same sex and similar weight as the HS test rat was placed into one of the cylindrical cages (kept the same throughout the experiment) 5 min before the HS subject rat was placed into the arena. The stimulus and subject rats were never housed together and thus were unfamiliar to each other. No social isolation was conducted on either rat. Each stimulus rat was used no more than once per day. The test duration was 20 min and was recorded using the same camera as that used in the OFT.

#### 2.3.4 Analysis of video data

Ethovision XT video tracking system (Version 4.0, Noldus Information Technology, The Netherlands) was used to analyze the videos recorded in all behavioral tests. After identifying the arena and calibrating the size of the arena, specific zones in the arena were outlined. For OFT and NOIT, one center zone, which was a circular region with a diameter of 20 cm, was used. For the SIT, one object zone and one social zone, both were circular regions with diameters of 20 cm, corresponding to the two cylindrical cages respectively, were specified. The extracted data included the total distance traveled in the arena, the duration and the frequency the test rat was present in specific zones, the distance of the subject to the zones, and the latency of the test rat entering the zones. The center of the subject rat was used for all calculations. Phenotypic correlations were determined using the Pearson test.

### 2.4 Pre-processing of phenotype data

For genetic analysis, each trait was quantile-normalized separately for males and females; this approach is similar to using sex as a covariate. Other relevant covariates (including age, batch number, and dissector) were identified for each trait, and covariate effects were regressed out if they were significant and if they explained more than 2% of the variance. Residuals were then quantile-normalized again, after which the data for each sex were pooled prior to further analysis. This approach removed mean differences due to sex; further, it did not attempt to model gene-by-sex interactions.

### 2.5 Genotyping and estimates of heritability

Genotypes were determined using genotyping-by-sequencing (GBS), as described previously [28]. This produced 3,513,494 SNPs with an estimated error rate *<*1%. Variants for X- and Y-chromosomes were not called. We used this set of SNPs for GWAS, genetic correlations, and heritability estimates. We used GCTA-GREML [29] analysis to estimate proportion of variance attributable to SNPs.

### 2.6 Genetic Mapping

GWAS analysis employed a linear mixed model, as implemented in the software GCTA [30], using a genetic relatedness matrix (GRM) to account for the complex family relationships within the HS population and the Leave One Chromosome Out (LOCO) method to avoid proximal contamination [31, 32]. Significance thresholds were calculated using permutation. Because all traits were quantile normalized, we used the same threshold for all traits [33]. To identify QTLs, we scanned each chromosome to determine if there was at least one SNP that exceeded the permutation-derived threshold of *−log*_10_(*p*) *>* 5.6, which was supported by a second SNP within 0.5 Mb that had a p-value that was within 2 *− log*_10_(*p*) units of the index SNP.

Other QTLs on the same chromosome were tested to ensure that they were independent of the first. To establish independence, we used the top SNP from the first QTL as a covariate and performed a second GWAS of the chromsome in question. If the resulting GWAS had an additional SNP with a p-value that exceeded our permutation-derived threshold, it was considered to be a second, independent locus. This process was repeated (including all previously significant SNPs as covariates), until no more QTLs were detected on a given chromosome. Linkage disequilibrium (LD) intervals for the identified QTL were determined by identifying those markers that had a high correlation coefficient with the peak marker (*r*^2^ = 0.6).

## 3 Results

### 3.1 Sex differences

We found that many of the traits measured in OFT, NOIT, and SIT are different between males and females (Table S1). In OFT, with the exception of latency of entering the center zone, all traits have statistically significant sex differences. In addition, four out of six traits in NOIT and seven out of eleven traits in SIT are different between males and females. The range of effect size (Cohen’ d) for statistically significant differences is (0.14, 0.31). Our genetic analysis quantile-normalized each trait separately for males and females. This approach removed mean differences due to sex and allowed us to combine males and females in the same analysis to increase the power of GWAS,

**Table 1:** Heritability of open field (OFT), novel object (NOIT) and social interaction (SIT) tests

| Test | Trait | Heritability ± SE |
| --- | --- | --- |
| OFT | Duration in center zone | $0.284 \pm 0.045$ |
| | Frequency of entering center zone | $0.323 \pm 0.044$ |
| | Latency of entering center zone | $0.083 \pm 0.034$ |
| | Mean distance to center zone | $0.295 \pm 0.043$ |
| | Total distance to center zone | $0.300 \pm 0.043$ |
| | Total travel distance | $0.379 \pm 0.044$ |
| NOIT | Duration in center zone | $0.247 \pm 0.043$ |
| | Frequency of entering center zone | $0.209 \pm 0.041$ |
| | Latency of entering center zone | $0.100 \pm 0.034$ |
| | Mean distance to center zone | $0.249 \pm 0.042$ |
| | Total distance to center zone | $0.221 \pm 0.041$ |
| | Total travel distance | $0.287 \pm 0.044$ |
| SIT | Duration in object zone | $0.161 \pm 0.037$ |
| | Duration in social zone | $0.275 \pm 0.040$ |
| | Frequency of entering object zone | $0.177 \pm 0.036$ |
| | Frequency of entering social zone | $0.215 \pm 0.036$ |
| | Latency of entering object zone | $0.082 \pm 0.032$ |
| | Latency of entering social zone | $0.142 \pm 0.034$ |
| | Mean distance to object zone | $0.165 \pm 0.038$ |
| | Mean distance to social zone | $0.265 \pm 0.041$ |
| | Total distance to object zone | $0.153 \pm 0.037$ |
| | Total distance to social zone | $0.265 \pm 0.041$ |
| | Total travel distance | $0.281 \pm 0.040$ |

### 3.2 Phenotypic correlations

We calculated Pearson correlation between the 23 traits (Figure 1). We found 197 correlations with un-adjusted p values less than 0.05. Most of these correlations have relatively low Person coefficient (mean is 0.23, median is 0.18). However, due to the large sample size, most of these correlations are highly significant (median *−log*_10_(*p*) is 7.8). In general, correlations of traits obtained from the same behavioral test are among the strongest. For example, frequency of visiting the center and duration of staying in the center are positively correlated in OFT (r=0.76), and duration in the social zone and distance to the social zone in the SIT are negatively correlated (r=-0.76). Most of these correlations are expected from the definitions of these variables.

**Figure 1:**
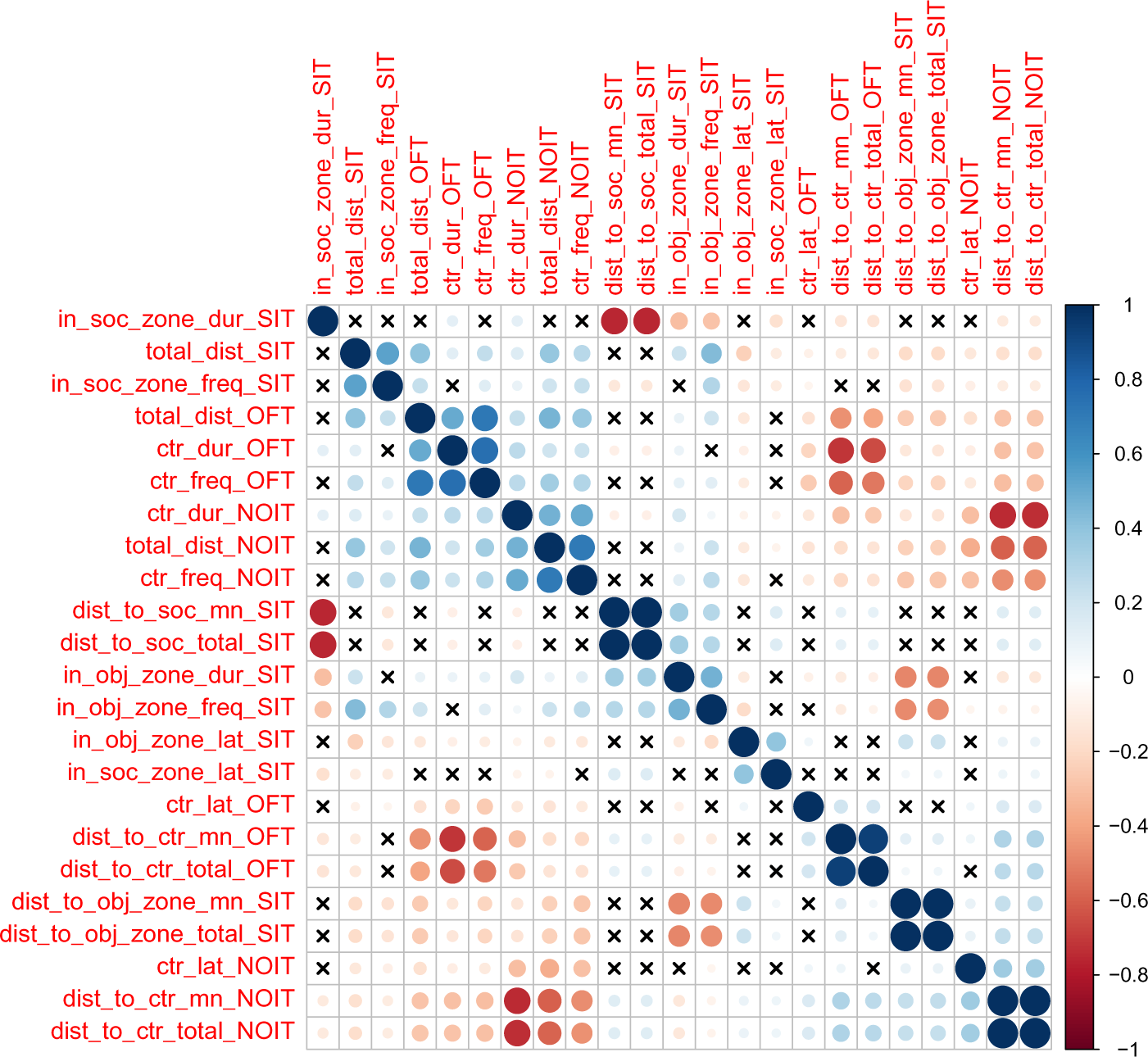
Heatmap showing the correlations between behavioral traits. The color scheme represents the direction of the correlation, whereas the intensity of the colors and the size of the circles are proportional to coefficients of the correlation. The cross signs indicates that the correlation of the two traits is not statistically significant (*p >* 0.05)

Among the correlations of variables derived from two different behavioral tests, correlations for measures of distance traveled are among the highest (range of Pearson r: 0.39 – 0.47, e.g. Figure 6 A, B). Distance traveled in the OFT is also correlated with duration of center time in the NOIT (e.g. Figure 6 C). Interestingly, the frequencies of visiting the center of the area in the NOIT is correlated with the frequency of visiting the social zone in the SIT (Figure 6 D). In contrast, OFT center frequency is negatively correlated with NOIT mean distance to center in NOIT (Figure 6 E), and distance to object zone in SIT is negatively correlated with center frequency in NOIT (Figure 6 F).

### 3.3 Heritability

SNP heritability estimates (*h*^2^) for traits are provided in Table 1. In all the three behavioral tests, total travel distance has the highest heritability. In OFT, all heritability estimates are between 0.28 – 0.38, with the exception of that for latency of entering the center zone (*h*^2^ = 0.08). Heritability estimates for variables from the NOIT are slightly lower than that of the OFT; most of them are in the range of 0.21 – 0.29, with the exception of that for the latency of entering the center zone (*h*^2^ = 0.10). Heritability estimates for various measures of the SIT are in the range of 0.10 – 0.28. Interestingly, heritability estimates for measures on the social zone are consistently greater than those for the object zone.

### 3.4 Identification of multiple GWAS hits

In Table 2, we present single nucleotide polymorphisms (SNPs) that are significantly associated with the phenotypes. The genome-wide statistical significance of the association is determined by *−log*_10_*P* values which ranges from 5.609 to 8.268. The p-values correspond to these are 2.46 *×* 10*^−^*^6^ and 0.5 *×* 10*^−^*^8^, respectively. For OFT, there are 9 significant loci for 5 traits. We did not find a significant QTL for *Duration in center zone* (*h*^2^ = 0.284 *±* 0.045). We identified two loci for *Frequency of entering center zone* and *Total travel distance*, 3 loci for *Total distance to center zone*. We found 4 NOIT traits have significant loci. Among them, *Total distance to center zone* has 3 loci and *Mean distance to center zone* has 2 loci. We did not find any significant loci for *Frequency of entering center zone* (*h*^2^ = 0.209 *±* 0.041) and *Latency of entering center zone* (*h*^2^ = 0.100 *±* 0.034). For SIT, we identified significant loci for all traits except *Latency of entering object zone* which has heritability of *h*^2^ = 0.082 *±* 0.032. We found 2 loci for the traits *Latency of entering social zone*, *Mean distance to social zone Total distance to social zone* and *Total travel distance*. All genome-wide significant loci are shown in Figure 2. Genetic mapping of individual traits are shown as Manhattan plots as Supplementary Figures S1–S23. Regional association plots for representative traits are shown in Fig 3– these traits are shown in Supplementary Figures 3–5 for OIT, NOIT, and SIT, respectively.

**Figure 2:**
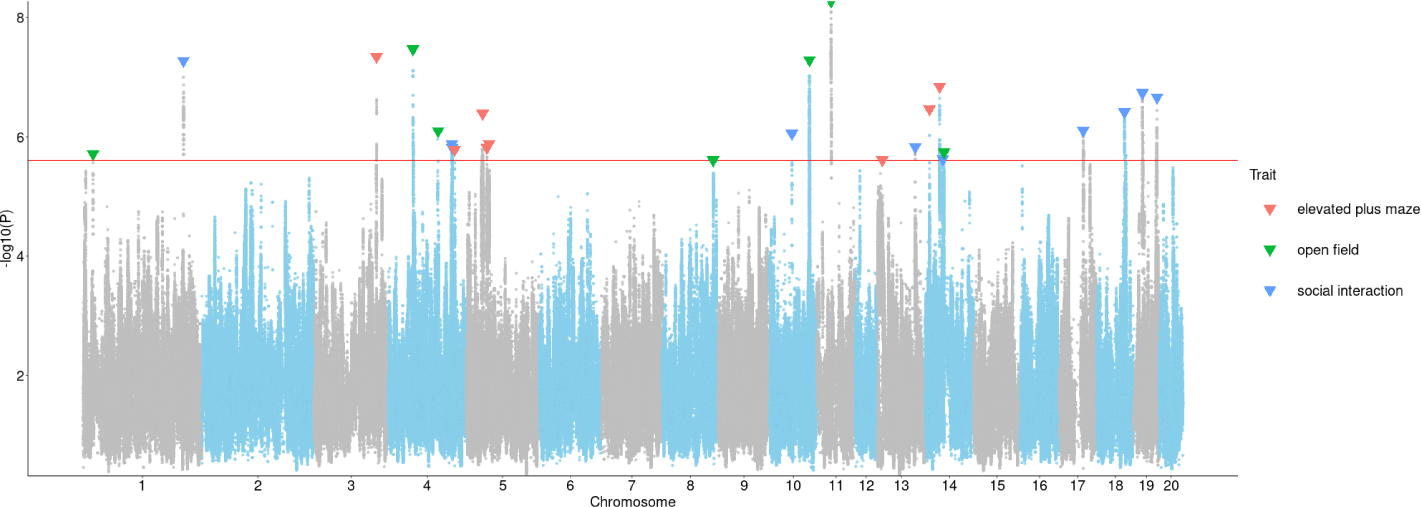
Association of approximately 3 million SNPs with behavioral traits measured in OFT, NOIT, or SIT. The red horizontal line denotes the p value for reaching genome-wide significance. The downward arrows denote the SNPs with the largest -log10(P) for each genome-wide significant association.

**Figure 3:**
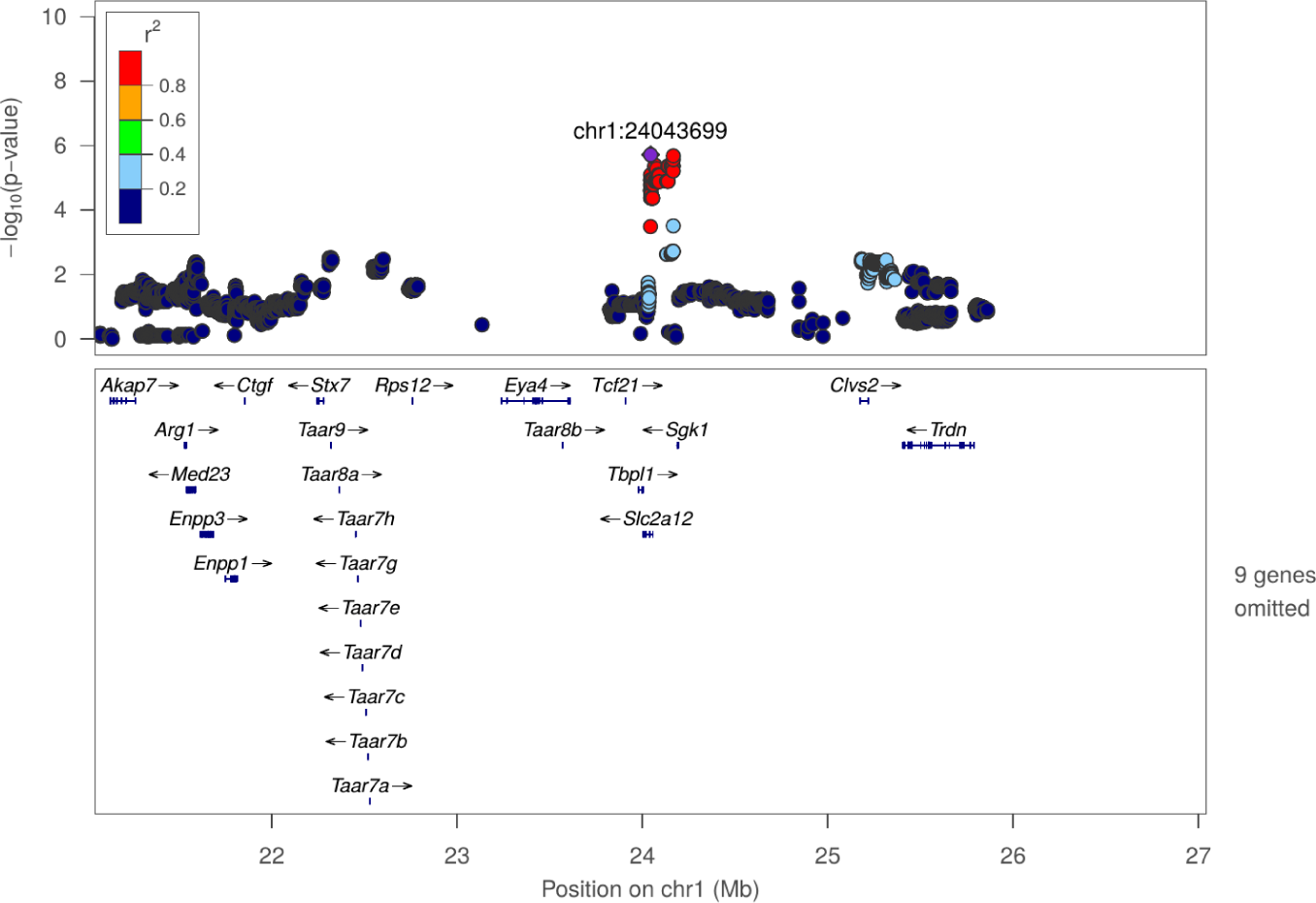
Regional association plot for frequency of entering center zone in OFT at chr1:24043699

**Figure 4:**
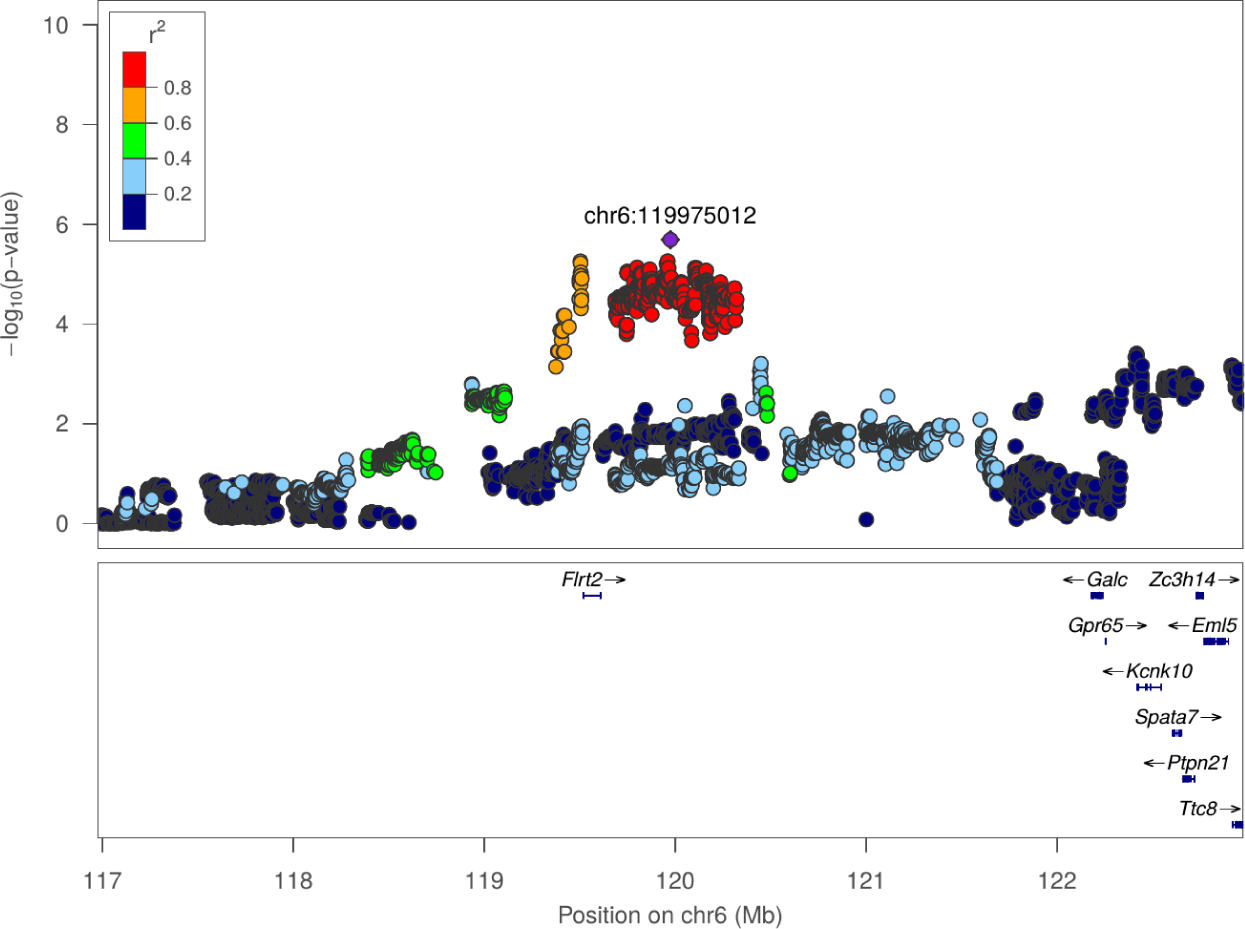
Regional association plot for mean distance to center zone in NOIT at chr6:119975012

**Figure 5:**
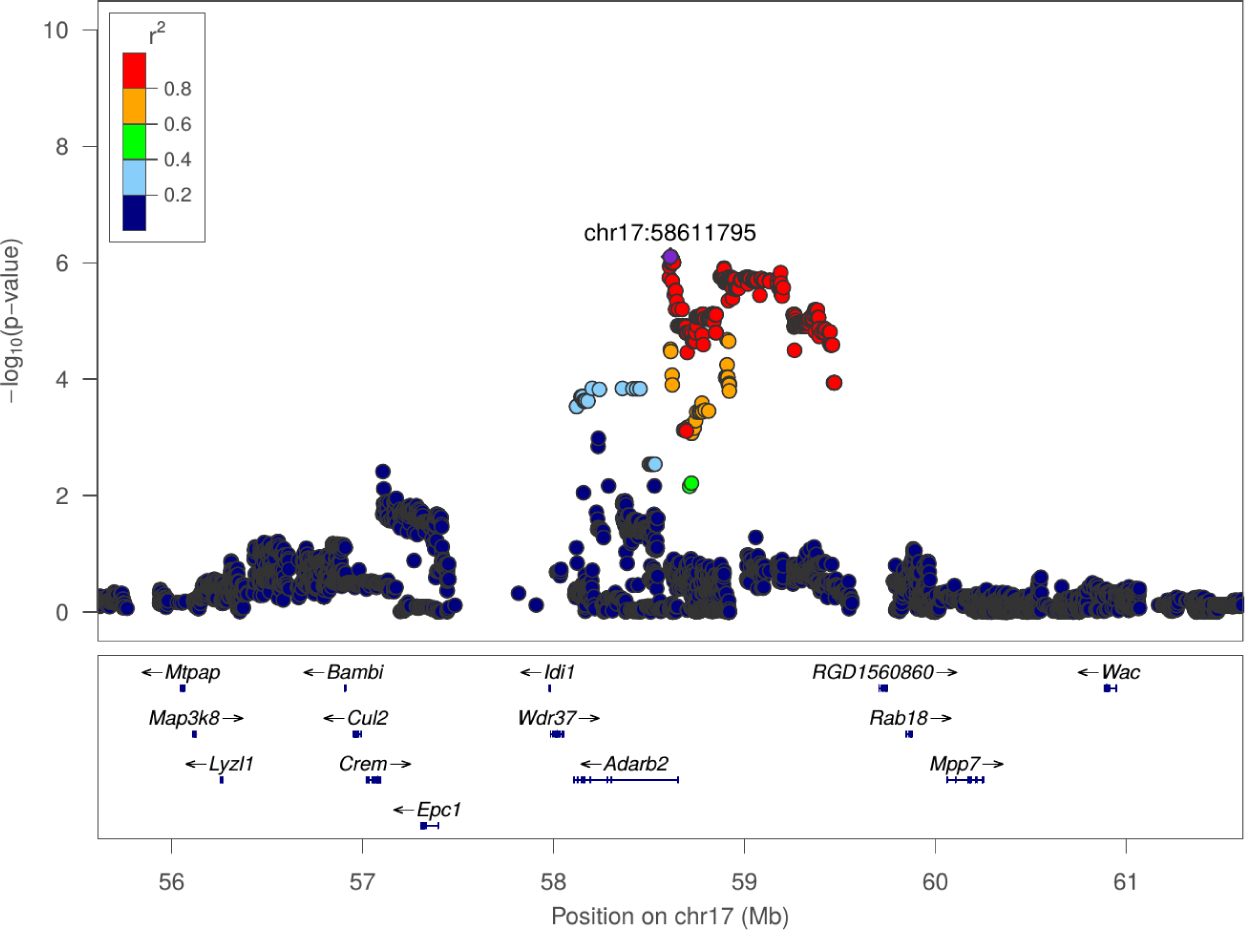
Regional association plot for latency of entering social zone in SIT at chr17:58611795

**Figure 6:**
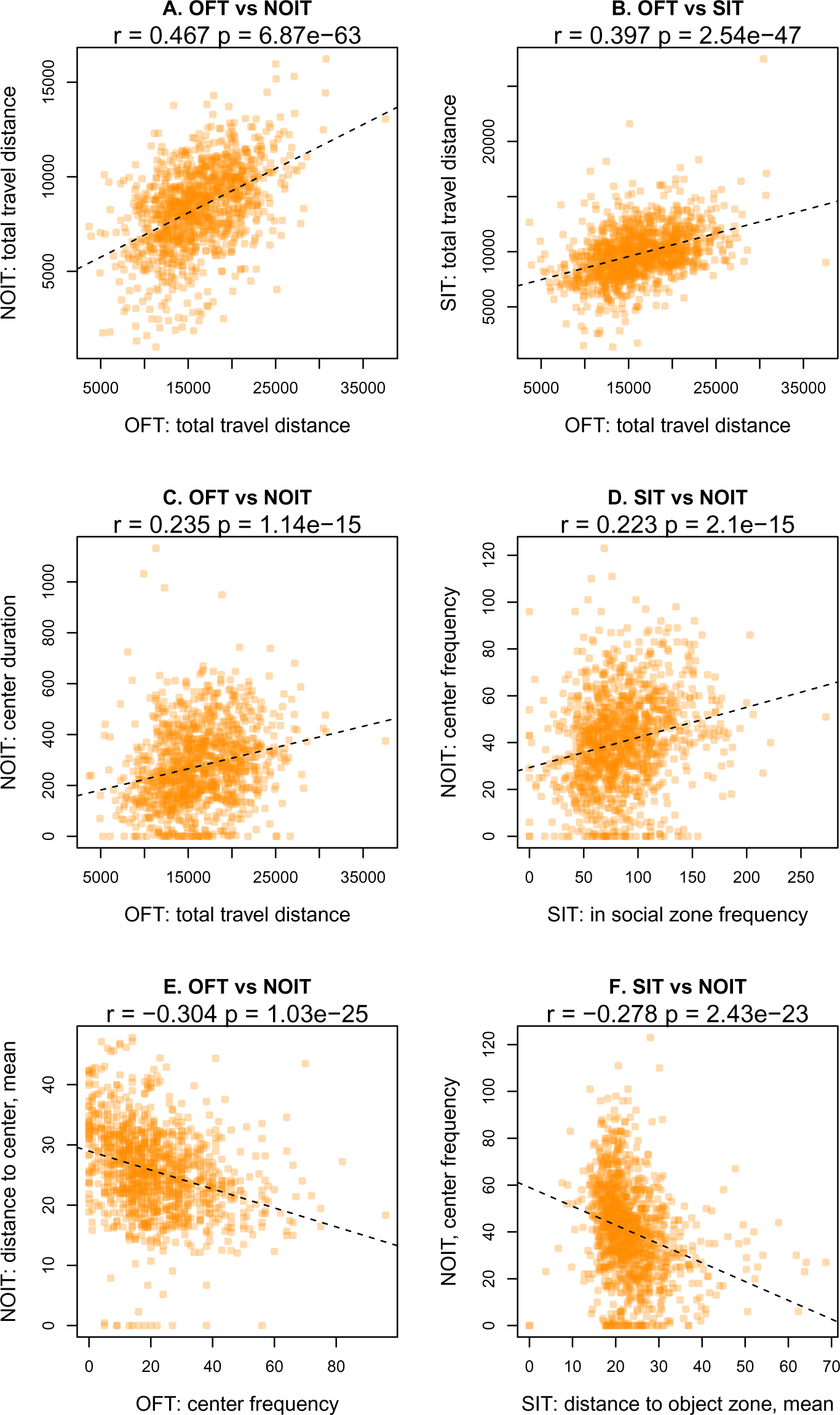
Selected scatter plots for correlation between behavioral tests shown in Figure 1.

**Table 2:**
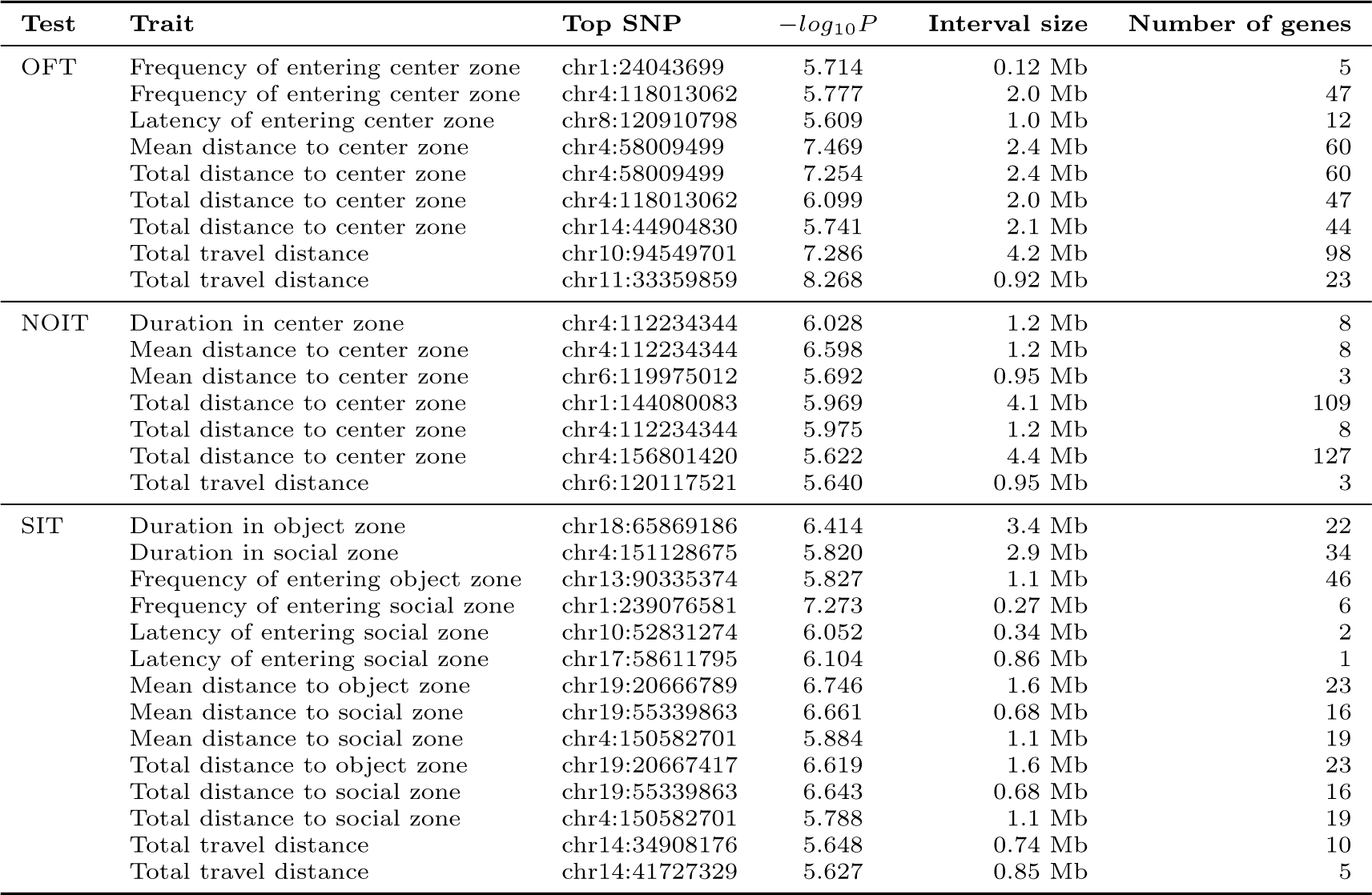
QTL for open field (OFT), novel object interaction (NOIT), and social interaction (SIT) tests

### 3.5 Pleiotropic loci

To determine if traits that mapped to the same location are pleiotropic, we considered the minor allele frequency (MAF), and the SDP of the index SNP among the 8 founder strains that were used to create the HS. Using these criteria, we did not observe any pleiotropic loci between the traits analyzed in different tests. However, we did identify pleiotropic loci between the traits of the same behavior test. Most of these traits are highly correlated, as shown in Figure 1. With the exception of three sets of QTL (Table S3), all others share the same top SNP (Table S2).

**Table 3.**
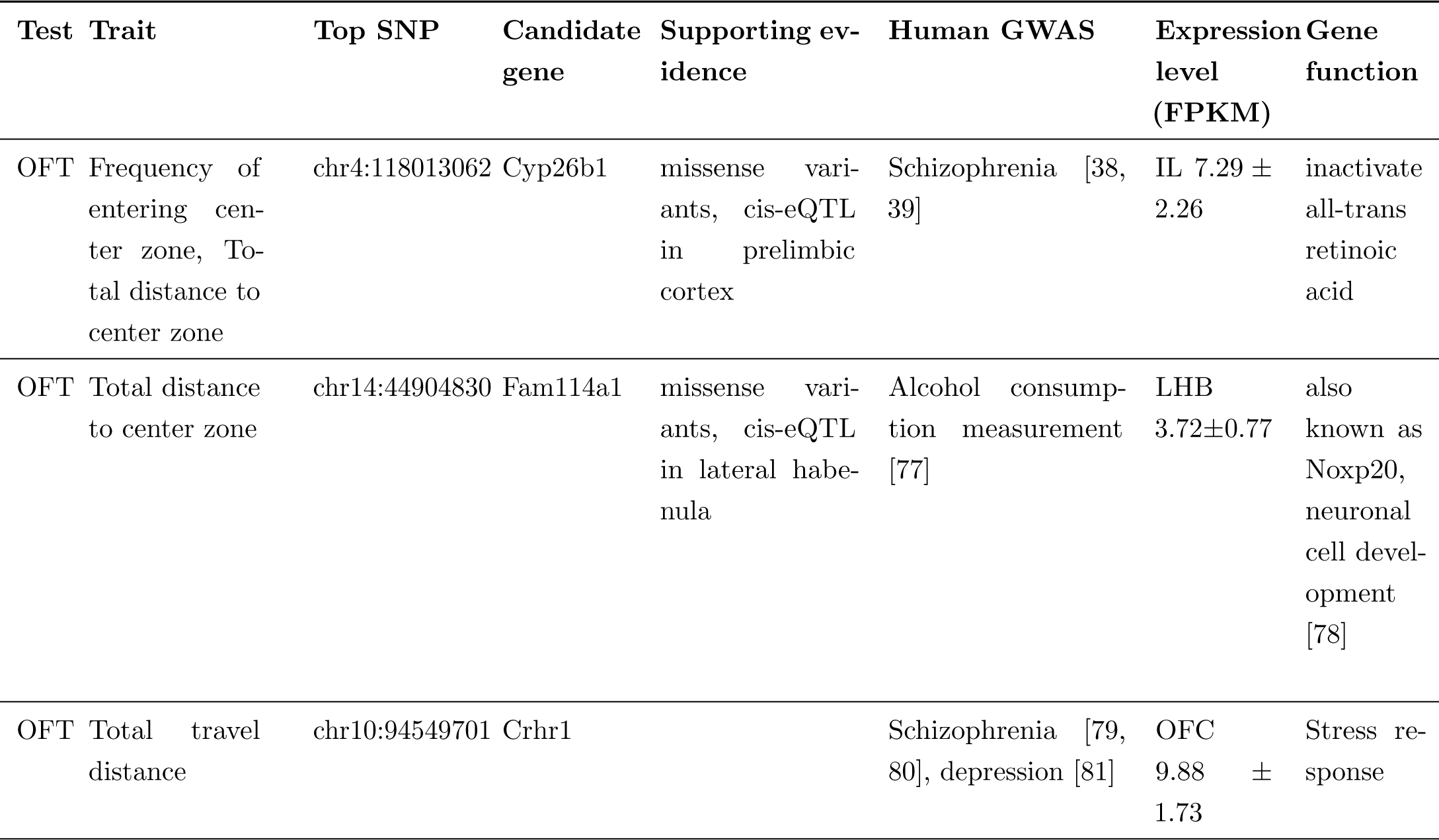
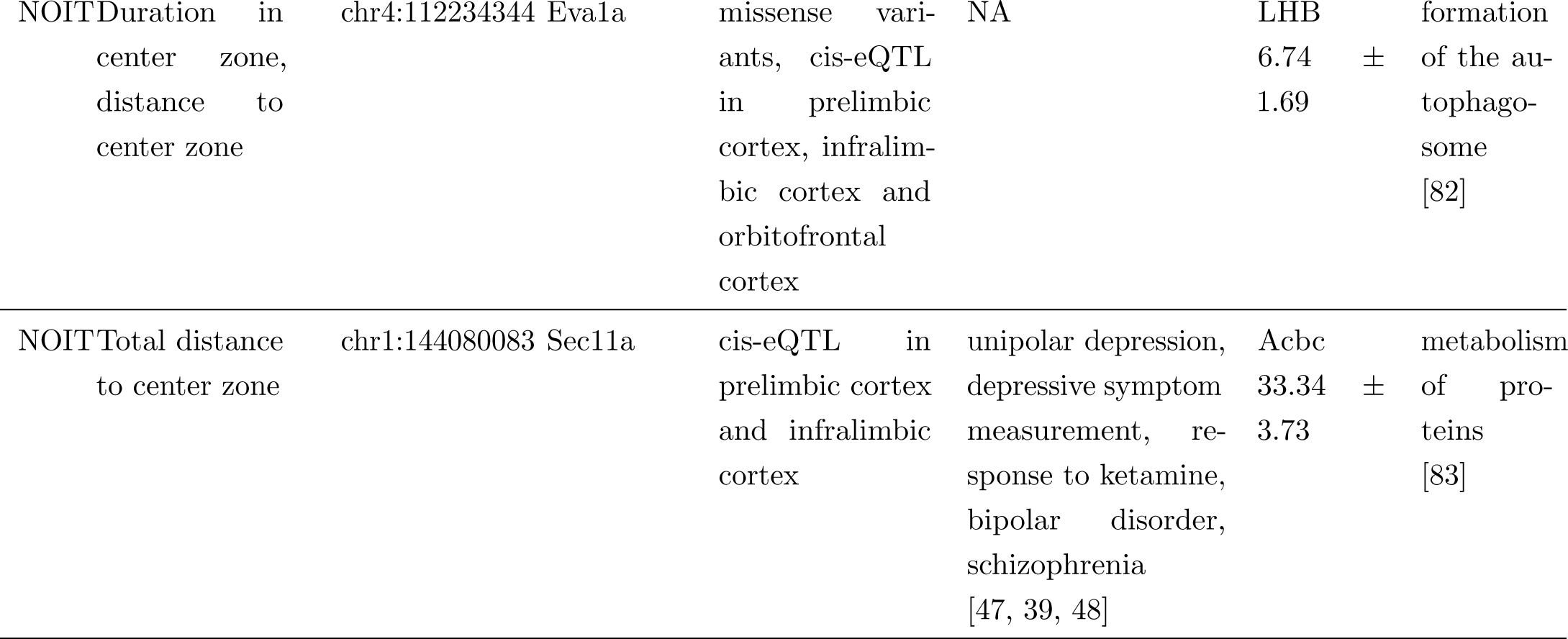
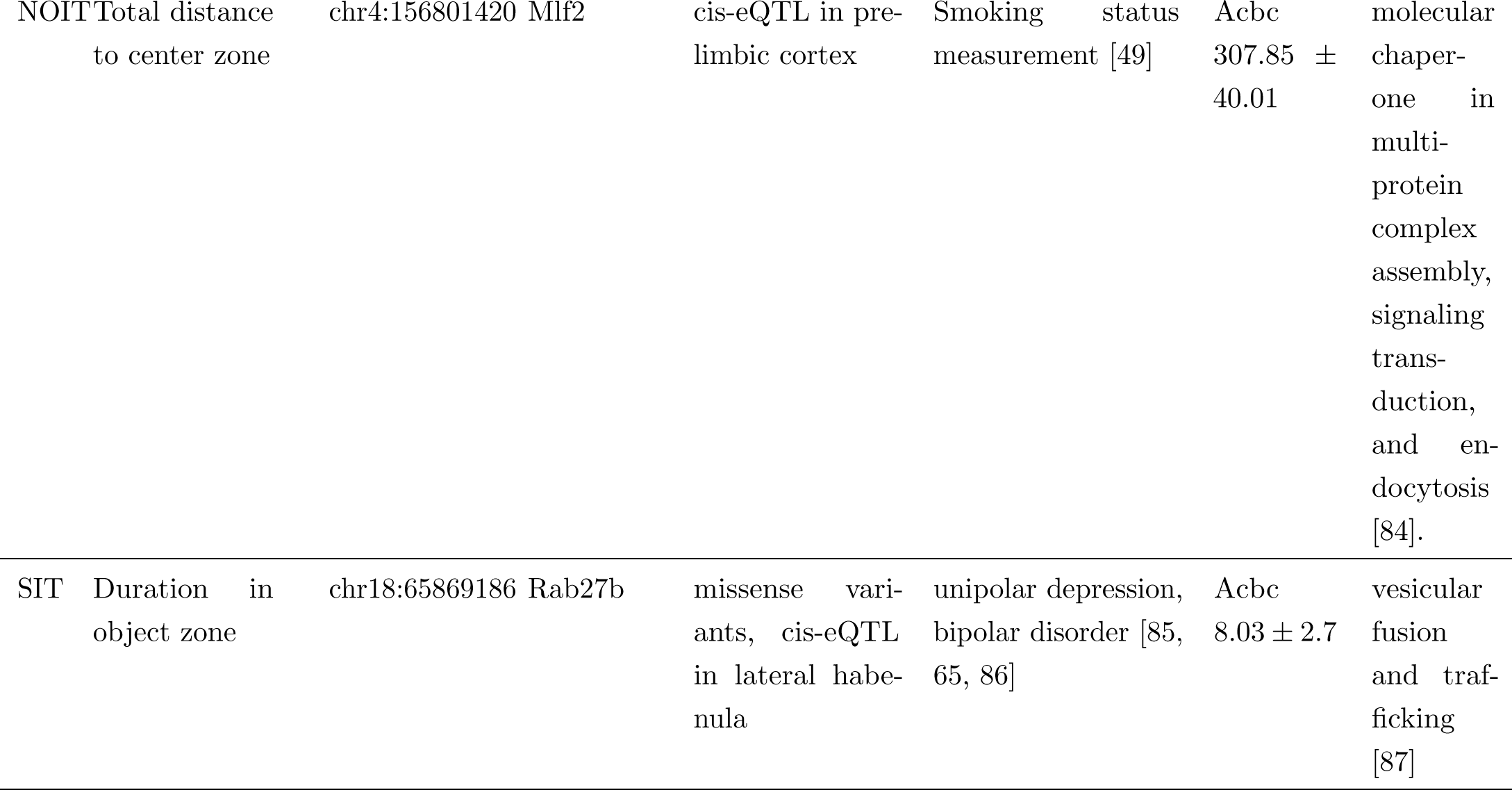
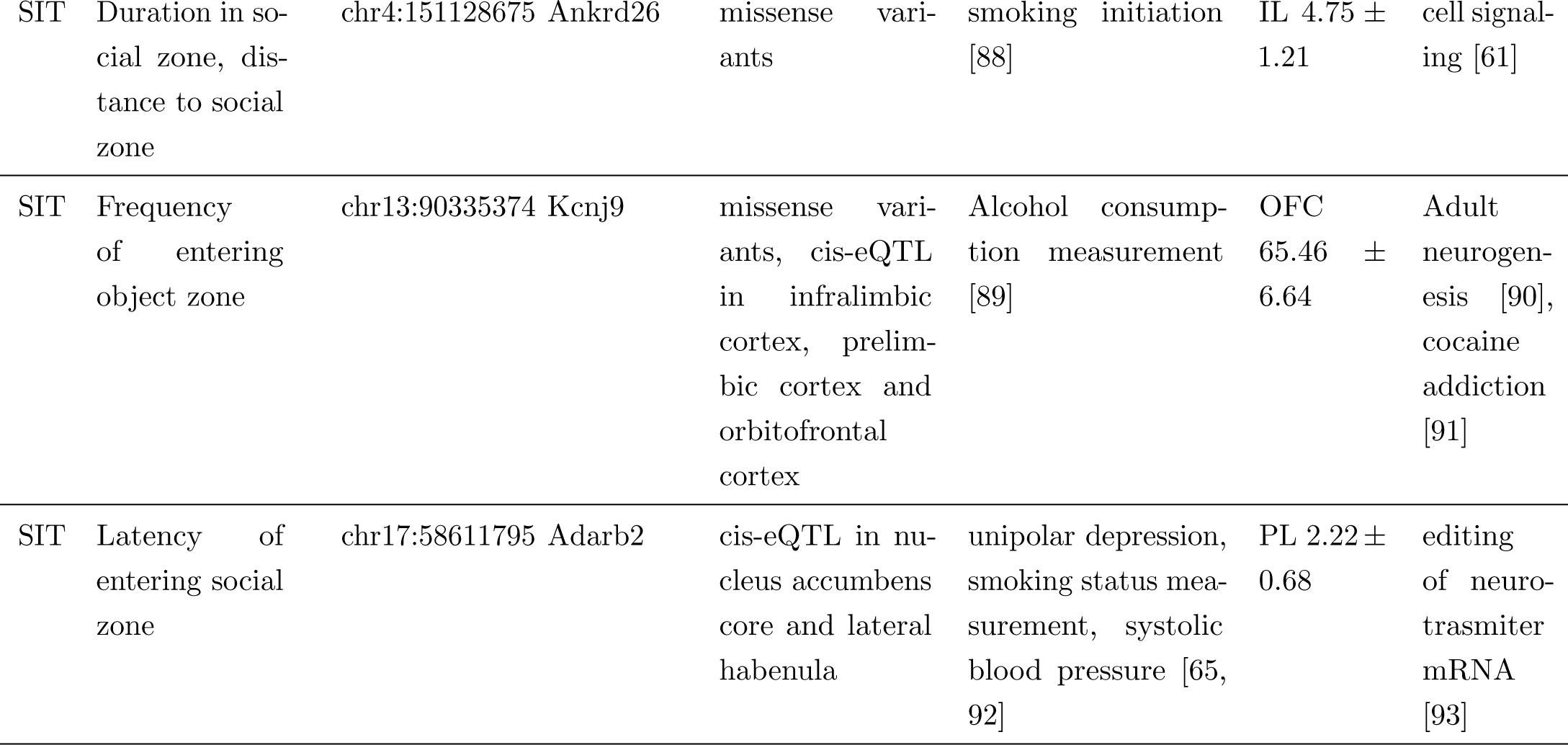
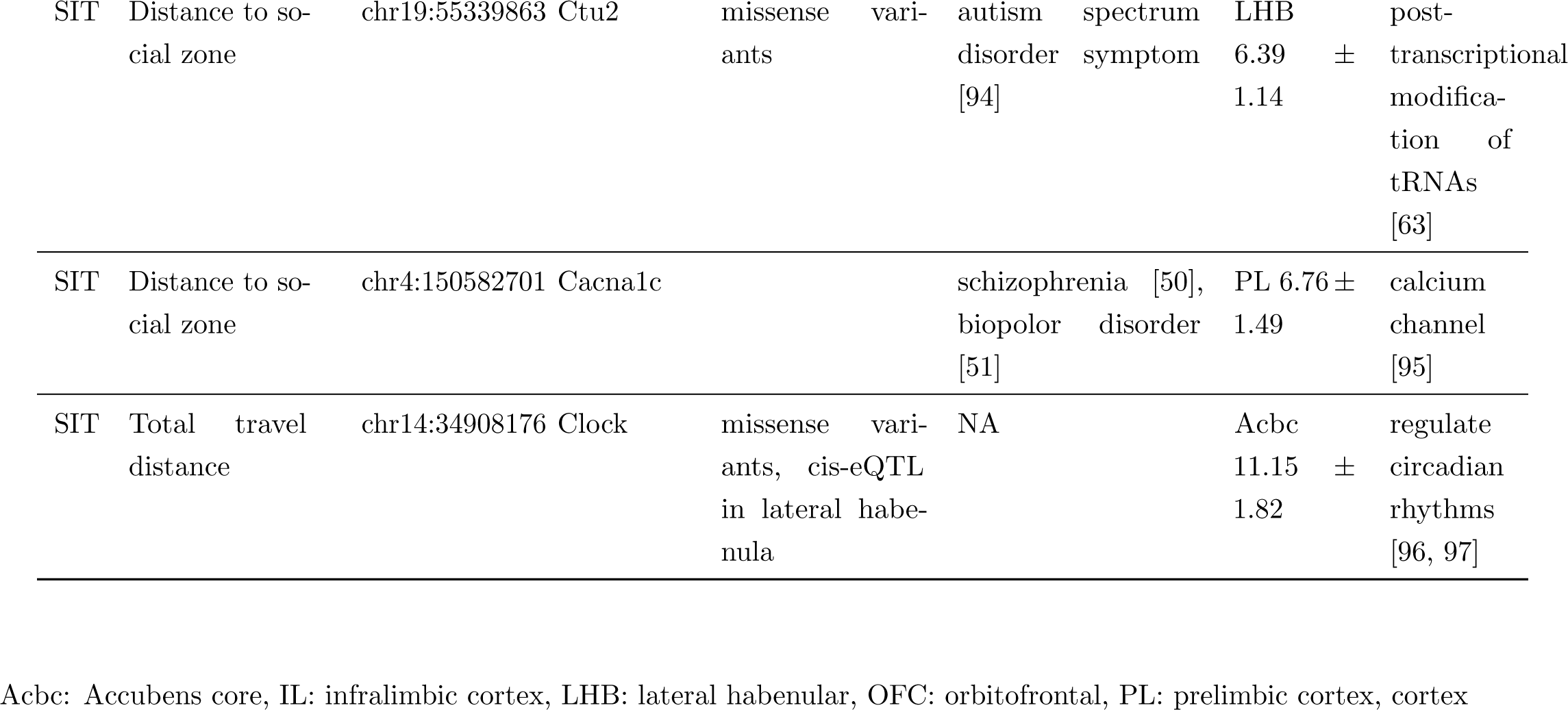
Candidate genes

### 3.6 Candidate gene identification

The number of genes within the identified QTL ranges from 1 to 127 (mean: 30.1, median: 19. Table 2). There is only one region that contains a single gene: *Adarb2* within chr17:58Mb for latency of entering social zone in SIT. However, it is also possible that the causal allele is a regulatory variant that is located in this interval but regulates a gene outside of the identified interval.

All other loci contained more than one gene. To identify candidate genes, we combined several criteria: 1) the presence of moderate or high impact variants located within the gene, as predicted by SnpEff [34]. We also require these variants are in high LD with the top SNP. We identified 149 coding variants within 30 QTL, 8 of which were predicted to have a high impact (Supplementary Table S4). 2) the presence of a significant cis-eQTL in one or more of the five brain regions in a dataset containing 88 navie adult HS rats[35], 3) has a human ortholog that has been reported to be associated with psychiatric diseases (including drug abuse). When multiple candidates are present using the above criteria, we remove the gene with very low expression levels across all five regions in the RNA-seq data set (e.g., FPKM *<* 0.5) and select the candidate with the strongest support for the literature. Combining these criteria with a literature search conducted using GeneCup [36], we identified plausible candidate genes within 13 loci (Table 3).

In addition, for total distance to the novel object zone, the QTL region on chr1 (144 Mb, size: 4.1 Mb, Figure S34) contains 69 gene with human orthologs. We found 14 of these genes have been reported in human GWAS to be associated with psychiatric conditions or addiction with genome-wide significance (*ACAN, ADAMTSL3, ALPK3, CPEB1, FES, FURIN, LINC00933, MIR9-3HG, MRPL46, NMB, POLG-DT, SEC11A, ZNF592, ZSCAN2*, Table S5). Three additional genes with sub-threshold significance in human GWAS are also included. These genes are all located in a syntenic region on human chromosome 15 (82.5-90.8 Mb). Although based on the criteria described above, *Sec11a* is the best candidate gene (Table 3), it is possible that this region contains multiple genes that are associated with the trait.

## 4 Discussion

As part of a GWAS on intravenous nicotine self-administration in adolescent HS rats that we are conducting [27, 37], we collected several behavioral phenotypes related to anxiety, novelty exploration, and social interaction. We have previously reported that these behavioral traits contribute to the variation in nicotine intake [27]. We report here GWAS results of three behavioral traits: OFT, NOIT, and SIT, which were all conducted in the same open field. We identified 24 QTL for 30 traits. Using a set criteria outlined above, we identified 13 candidate genes.

OFT, NOIT, and SIT are widely used behavioral assays in rodents. With over 1,200 rats, ours represent some of the largest data collected using these assays. Similar to our interim report on this data set [27], we found a large number of correlations with relatively low coefficients (e.g., *r <* 0.4) but with high statistical significance. It is likely that these correlated traits are controlled by the same behavioral processes and thus are influenced by the same genetic factors. In fact, our genetic analysis did find several pleiotropic sites (Table S3). Almost all pleiotropic loci are reported for traits measured in the same behavior assay. It is likely that further increasing sample size will provide greater statistical power to detect pleiotropic effect across different behavioral assays.

Many of the candidate genes in this study have been associated with psychiatric or drug abuse traits in humans. For example, we identified *Cyp26b1*, a retinoic acid degrading enzyme, as a candidate gene for the frequency of entering the center zone and total distance to the center zone in OFT; both of which are measures of anxiety-like behaviors. *Cyp26b1* has been associated with Schizophrenia in several human GWAS [38, 39]. Anxiety symptoms are common in schizophrenia patients [40, 41]. *Cyp26b1* is is expressed in parvalbumin-positive interneurons [42]. Most interestingly, knockdown *Cyp26b1* in the nucleus accumbens shell decreased anxiety-like behavior [43].

The *Crhr1* gene, which encodes corticotrophin release hormone receptor 1, is a candidate gene for total travel distance in the OFT. *Crhr1* is involved in anxiety-like behavior in OFT in rats [44, 45]. In mice, conditional knockout approach showed that *Crhr1* in the forebrain underlies the effect of early life stress on total travel distance in the OFT [46], which provides a direct confirmation for the association we report here.

Among the candidate genes for NOIT, *Eva1a*, a candidate gene for the duration in the center zone, is supported by strong cis-eQTL and a missense variant. *Eva1a* has no literature support and thus could lead to the discovery of new mechanisms for novelty seeking-like behavior. *Sec11a*, a candidate gene for total distance in the center zone, is associated with depression and schizophrenia [47, 39, 48]. *Mlf2*, a candidate gene for total distance to center zone in NOIT, is associated with smoking in humans [49] and has very high expression levels in the accumbens (Table 3).

For the SIT, we identified *Cacna1c*, encoding the Ca_v_1.2 subunit of the L-type Ca^2+^ channel, as a candidate gene for distance to the social zone. *Cacna1c* has been associated with schizophrenia [50] and bipolar disorder [51] in human GWAS. Both schizophrenia and bipolar disorders are associated with impairments in a range of social deficits [52, 53]. In animal studies, Sprague-Dawley rats with heterozygotic deletion of the *Cacna1c* gene (homozygotic mutation is lethal) showed many deficits in social behavior. These included reduced levels of ultrasonic vocalizations during rough-and-tumble play, as well as social approach behavior elicited by playback of ultrasonic vocalizations [54, 55]. In mice, a knockdown of *Cacna1c* in the nucleus accumbens significantly increased susceptibility to social stress [56]. Knocking down of *Cacna1c* in the prefrontal cortex of adult mice also recapitulated many of the social deficits [57]. Importantly, some of the behavioral effects of *Cacna1c* appear to interact with genetic background [58].

Among the other candidate genes for the SIT traits, *Rab27b* is involved in the presynaptic mechanism of long-term potentiation [59] as well as myelin biogenesis in oligodendrocytes [60]. *Ankrd26* is expressed in the arcuate and ventromedial nuclei and in the ependyma[61]. *Kcnj9* is involved in neurite outgrowth [62]. *Ctu2* is involved in post-translational modification of tRNAs [63]. *Adarb2* has been associated with home cage activity [64] and unipolar depression [65]. The *Clock* gene is involved in the maintenance of locomotor rhythms [66]. Mutations of the *CLOCK* gene have been implicated in many psychiatric disorders [67]. Although these candidates are well supported by multiple lines of evidence, additional work is needed to confirm their causal relationship to the corresponding behavioral traits.

The total distance to the novel object zone is associated with chr1:144080083 (allele frequency: 0.91, *−log*_10_(*p*) = 5.969, size of interval: 4.1 Mb, Figure S34). This SNP is also associated with the duration rats stayed in the novel object zone, although the p value did not reach genome-wide significance (-logP=4.63). This region contains 69 known genes. Its syntenic region on human Chr15 (82.5-90.8 Mb) is a hotspot for human pyschiatric diseases, containing 30 SNPs and 14 genes (*ACAN, ADAMTSL3, ALPK3, CPEB1, FES, FURIN, LINC00933, MIR9-3HG, MRPL46, NMB, POLG-DT, SEC11A, ZNF592, ZSCAN2*) associated with generalized anxiety disorder, schizophrenia, bipolar disorder, obsessive compulsive disorder, attentions deficit hyperactivity disorder, autism spectrum disorder, and unipolar depression, smoking behavior, etc. These results are reported in 21 publications (Table S5). Using the criteria described above, we identified *Sec11a* as the best candidate gene (Table 3). However, given the large number of genetic variants reported in human GWAS that are associated with psychiatric conditions within this syntenic region, it is very likely that this region contains multiple genes that are associated with novelty seeking-like behavior.

We include overlapping with human psychiatric GWAS results as part of the criteria in prioritizing candidate genes. It is possible that this approach could introduce bias and prevent us from making novel discoveries. For example, two (*Cyp26b1* and *Crhr1*) of the three candidate genes for OFT have been associated with schizophrenia, rather than anxiety. However, many genetic variants are pleiotropic for multiple psychiatric diseases [68]. For example, polygenic risk scores for schizophrenia have been associated with many other psychiatric diseases, such as anxiety disorder [69] or major depressive disorder [70], or cognitive performance [71]. Together with other evidence, we believe considering human psychiatric GWAS findings when identifying candidate genes in our study, even when the behavior trait in rats does not map directly to the psychiatric disease, is still valid and will likely increase the transnational value of our findings.

The presence of cis-eQTL in the brain is one of the strongest pieces of evidence that we use to prioritizes candidate genes. Nine of the 13 candidate genes we identified have cis-eQTL. Two of the strongest candidate genes in our results, *Cacna1c* for social behavior and *Crhr1* for anxiety-like behavior, are both supported by prior studies on similar traits using knockout mice. However, we did not find significant cis-eQTL of these two genes in our dataset. This could imply either our cis-eQTL dataset lack sufficient power or that genetic regulation of the traits does not directly involve gene expression in the brain regions that we have eQTL data. In addition, several QTL regions contain multiple cis-eQTL. It is possible this is due to strong LD within the region.

The HS rat population has already been successfully used in genetic mapping studies of physiological or behavioral traits [72, 73, 26]. Prior study mapped several anxiety-like traits using zero maze [23]. Several GWAS using HS to study behavioral regulation [74], response to cocaine cues [75], cocaine self-administration [76], nicotine self-administration [37, 27], or oxycodone self-administration are underway. Our study add to the literature 30 QTLs and 13 candidate genes for psychiatric related behavioral traits. While some of the candidate genes are well supported by knockout studies in mice and human GWAS, others likely represent novel findings that can be the catalyst for future molecular and genetic insights on psychiatric diseases.

## Author Contributions

H.C. and A.A.P. designed the study. T.W. and A.G.M collected the data. A.S.C., O.P. and M.H.G. analyzed the data. The manuscript was written by M.H.G., A.A.P. and H.C. All authors contributed to the article and approved the submitted version.

## Acknowledgments

The authors thank Wenyan Han, Yanyan Lin, and Pawandeep Kaur for their contributions in collecting some of the behavioral data. We thank the GeneNetwork team for hosting the data.

## Data Availability Statement

The datasets generated for this study can be found in GeneNetwork (http://www.genenetwork.org).

---

**Table S1:** Sex differences of OFT, NOIT and SIT (Mean *±* SD)

| Test | Trait | Female | Male | p-value | Cohen's d |
| --- | --- | --- | --- | --- | --- |
| OFT | Duration in center zone | 65 ± 51.4 | 55.7 ± 43.5 | 0.0006 | 0.19 |
| | Frequency of entering center zone | 22.7 ± 14 | 19.9 ± 13.7 | 0.0003 | 0.21 |
| | Latency of entering center zone | 250.8 ± 301.3 | 271.6 ± 305.1 | 0.23 | 0.07 |
| | Mean distance to center zone | 36.3 ± 3.9 | 37.1 ± 3.9 | 0.0002 | 0.21 |
| | Total distance to center zone | 1930273 ± 224949 | 1976133 ± 229112 | 0.0004 | 0.20 |
| | Total travel distance | 16578 ± 4372 | 15762 ± 4272 | 0.0009 | 0.19 |
| NOIT | Duration in center zone | 283.9 ± 151.7 | 268.1 ± 160.7 | 0.08 | 0.10 |
| | Frequency of entering center zone | 40.4 ± 19.3 | 40.2 ± 20.6 | 0.92 | 0.01 |
| | Latency of entering center zone | 88.4 ± 123.8 | 108.3 ± 148 | 0.01 | 0.15 |
| | Mean distance to center zone | 24.6 ± 7.2 | 26.6 ± 7.3 | <0.0001 | 0.27 |
| | Total distance to center zone | 437507 ± 127709 | 473355 ± 130656 | <0.0001 | 0.28 |
| | Total travel distance | 8611 ± 2113 | 8217 ± 2226 | 0.001 | 0.18 |
| SIT | Duration in object zone | 141.2 ± 77.4 | 163.1 ± 86.9 | <0.0001 | 0.27 |
| | Duration in social zone | 454.3 ± 208.8 | 392.4 ± 187.5 | <0.0001 | 0.31 |
| | Frequency of entering object zone | 44.9 ± 20.8 | 50 ± 25.1 | <0.0001 | 0.22 |
| | Frequency of entering social zone | 85.2 ± 37.2 | 87 ± 32.7 | 0.35 | 0.05 |
| | Latency of entering object zone | 35.9 ± 51.2 | 37 ± 68.5 | 0.73 | 0.02 |
| | Latency of entering social zone | 5.2 ± 22.8 | 7.4 ± 45 | 0.28 | 0.06 |
| | Mean distance to object zone | 22.6 ± 6.7 | 23.6 ± 6.8 | 0.01 | 0.14 |
| | Mean distance to social zone | 12.3 ± 4.9 | 13.7 ± 4.5 | <0.0001 | 0.30 |
| | Total distance to object zone | 400955 ± 117623 | 420714 ± 121683 | 0.003 | 0.17 |
| | Total distance to social zone | 218013 ± 87249 | 244171 ± 80511 | <0.0001 | 0.31 |
| | Total travel distance | 9819 ± 2239 | 9841 ± 2312 | 0.86 | 0.01 |

**Table S2:** Strain distribution pattern of nearby QTL

| Top SNP | AF | SE | Beta | Allele1 | Allele2 | ACI | BN | BUF | F344 | M520 | MR | WN | WKY |
| --- | --- | --- | --- | --- | --- | --- | --- | --- | --- | --- | --- | --- | --- |
| chr4:150582701 | 0.362 | 0.042 | -0.203 | C | A | AA | CC | AA | AA | AA | AA | AA | AA |
| chr4:151128675 | 0.512 | 0.040 | -0.192 | C | A | CC | CC | CC | CC | CC | CC | CC | AA |
| chr6:119975012 | 0.696 | 0.048 | -0.229 | C | T | CC | CC | CC | CC | TT | CC | CC | CC |
| chr6:120117521 | 0.312 | 0.046 | -0.216 | T | G | GG | TT | 00 | GG | TT | GG | GG | GG |
| chr19:20666789 | 0.946 | 0.088 | -0.458 | G | C | GG | GG | GG | GG | CC | GG | GG | CC |
| chr19:20667417 | 0.946 | 0.088 | -0.452 | A | G | AA | AA | AA | AA | AA | AA | AA | GG |

**Table S3:** Pleiotropic effects.

| Top SNP | Trait | $-\log_{10}P$ |
| --- | --- | --- |
| chr1:144080083 | NOIT distance to center, total | 5.969 |
|  | NOIT center duration | 4.633 |
|  | NOIT distance to center, mean | 5.143 |
| chr4:58009499 | OFT distance to center, total | 7.254 |
|  | OFT distance to center, mean | 7.469 |
|  | OFT total travel distance | 4.191 |
| chr4:112234344 | NOIT center duration | 6.028 |
|  | NOIT distance to center, mean | 6.598 |
|  | NOIT distance to center, total | 5.975 |
| chr4:118013062 | OFT center frequency | 5.777 |
|  | OFT distance to center, total | 6.099 |
|  | OFT center duration | 5.373 |
|  | OFT distance to center, mean | 5.001 |
| chr4:150582701 | SIT distance to social zone, mean | 5.884 |
|  | SIT distance to social zone, total | 5.788 |
|  | SIT in social zone duration | 4.324 |
| chr4:151128675 | SIT in social zone duration | 5.820 |
|  | SIT distance to social zone mean | 4.351 |
|  | SIT distance to social zone, total | 4.366 |
| chr4:156801420 | NOIT distance to center, total | 5.622 |
|  | NOIT distance to center, mean | 4.543 |
| chr6:119975012 | NOIT distance to center, mean | 5.692 |
|  | NOIT center frequency | 4.275 |
|  | NOIT distance to center, total | 5.542 |
|  | NOIT total travel distance | 5.270 |
| chr6:120117521 | NOIT total travel distance | 5.640 |
|  | NOIT center frequency | 4.330 |
|  | NOIT distance to center, mean | 5.021 |
|  | NOIT distance to center, total | 4.961 |
| chr11:33359859 | OFT total travel distance | 8.268 |
|  | OFT center duration | 4.125 |
|  | OFT center frequency | 4.895 |
| chr14:44904830 | OFT distance to center, total | 5.741 |
|  | OFT center frequency | 4.189 |
|  | OFT distance to center, mean | 5.096 |
| chr14:41727329 | SIT total travel distance | 5.627 |
|  | NOIT total travel distance | 4.291 |
| chr19:20666789 | SIT distance to object zone, mean | 6.746 |
|  | SIT distance to object zone, total | 6.613 |
| chr19:20667417 | SIT distance to object zone, total | 6.619 |
|  | SIT distance to object zone, mean | 6.732 |
| chr19:55339863 | SIT distance to social zone, mean | 6.661 |
|  | SIT distance to social zone, total | 6.643 |

**Table S4.**
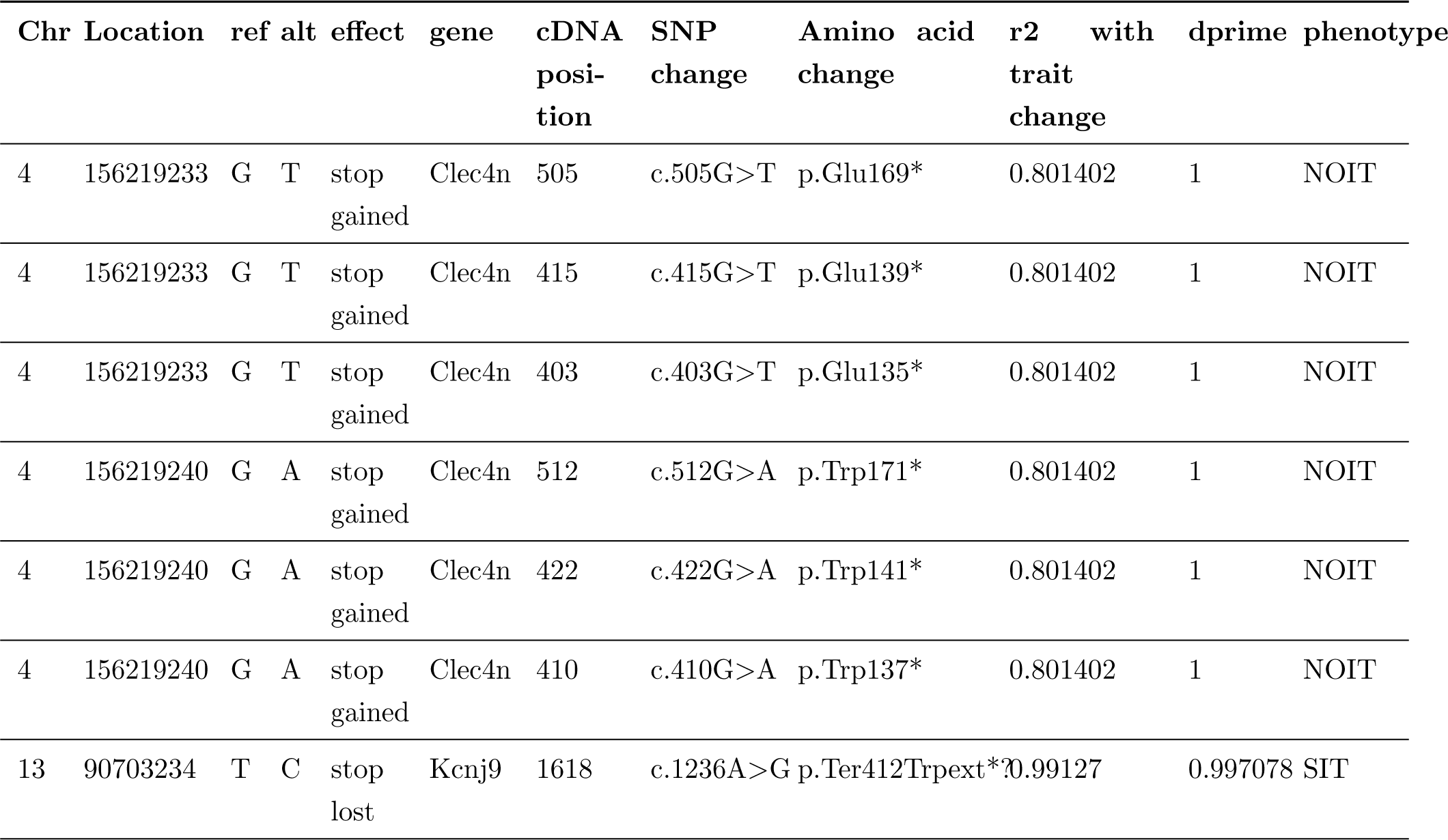
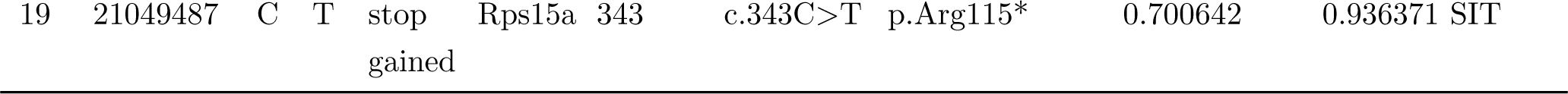
Putatively causal coding variants, HIGH impact

**Table S5.**
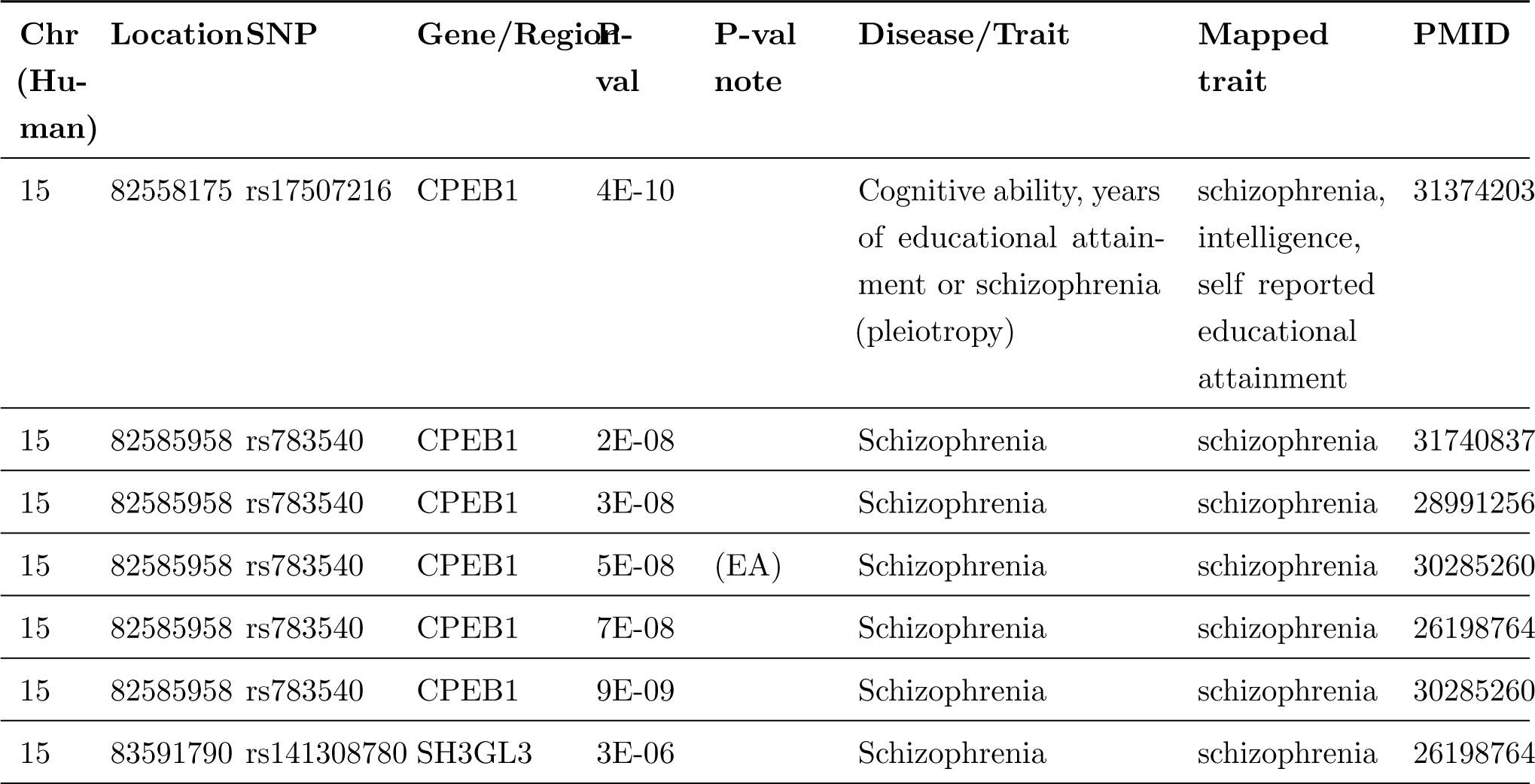
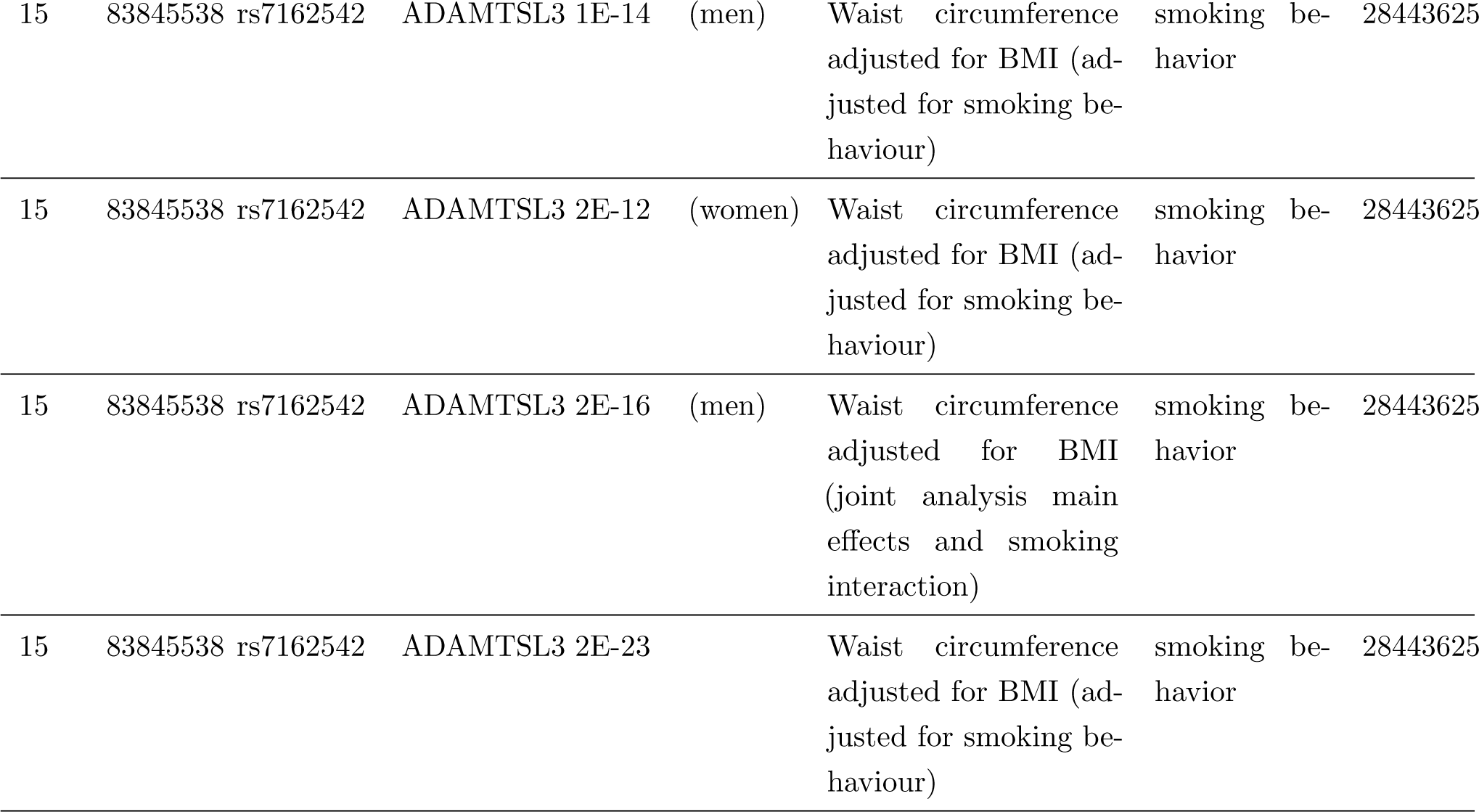
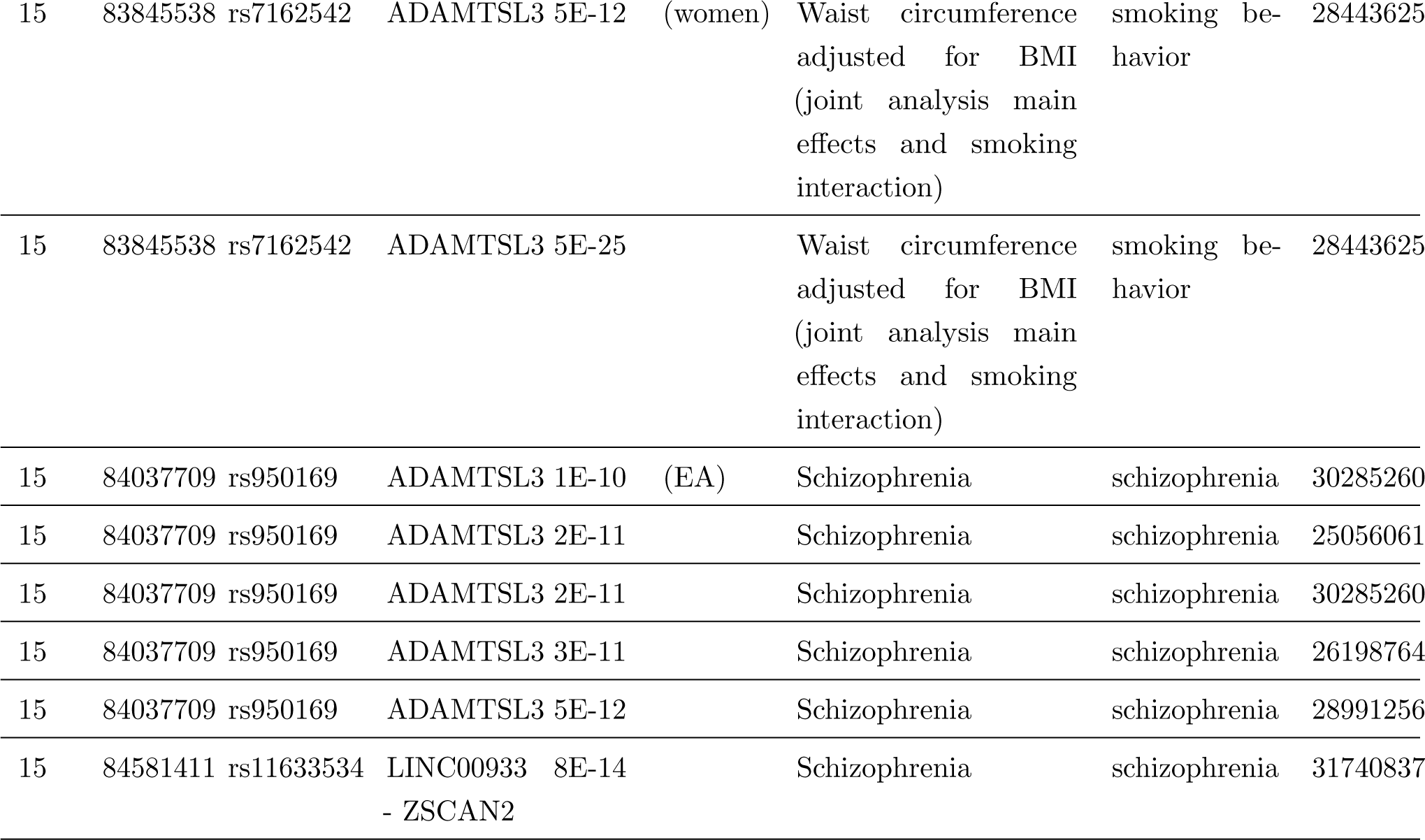
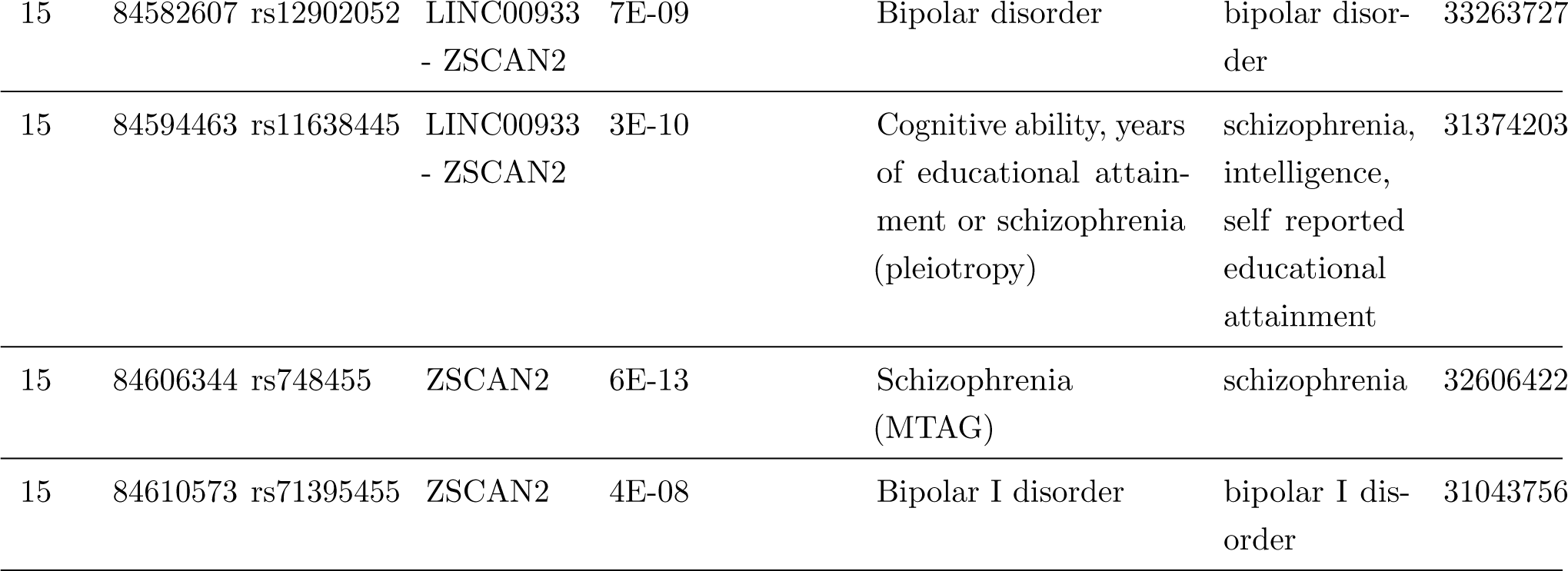
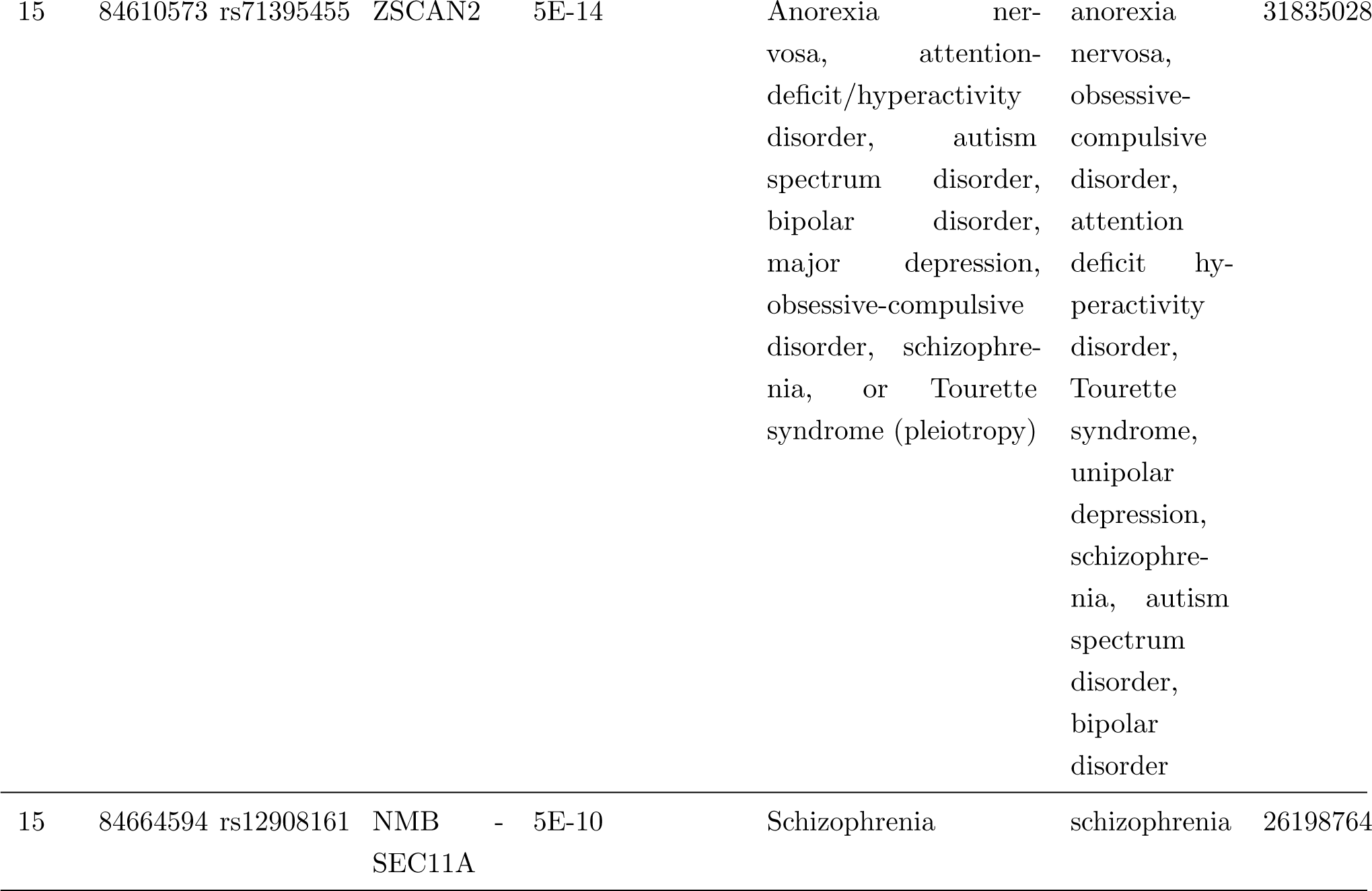
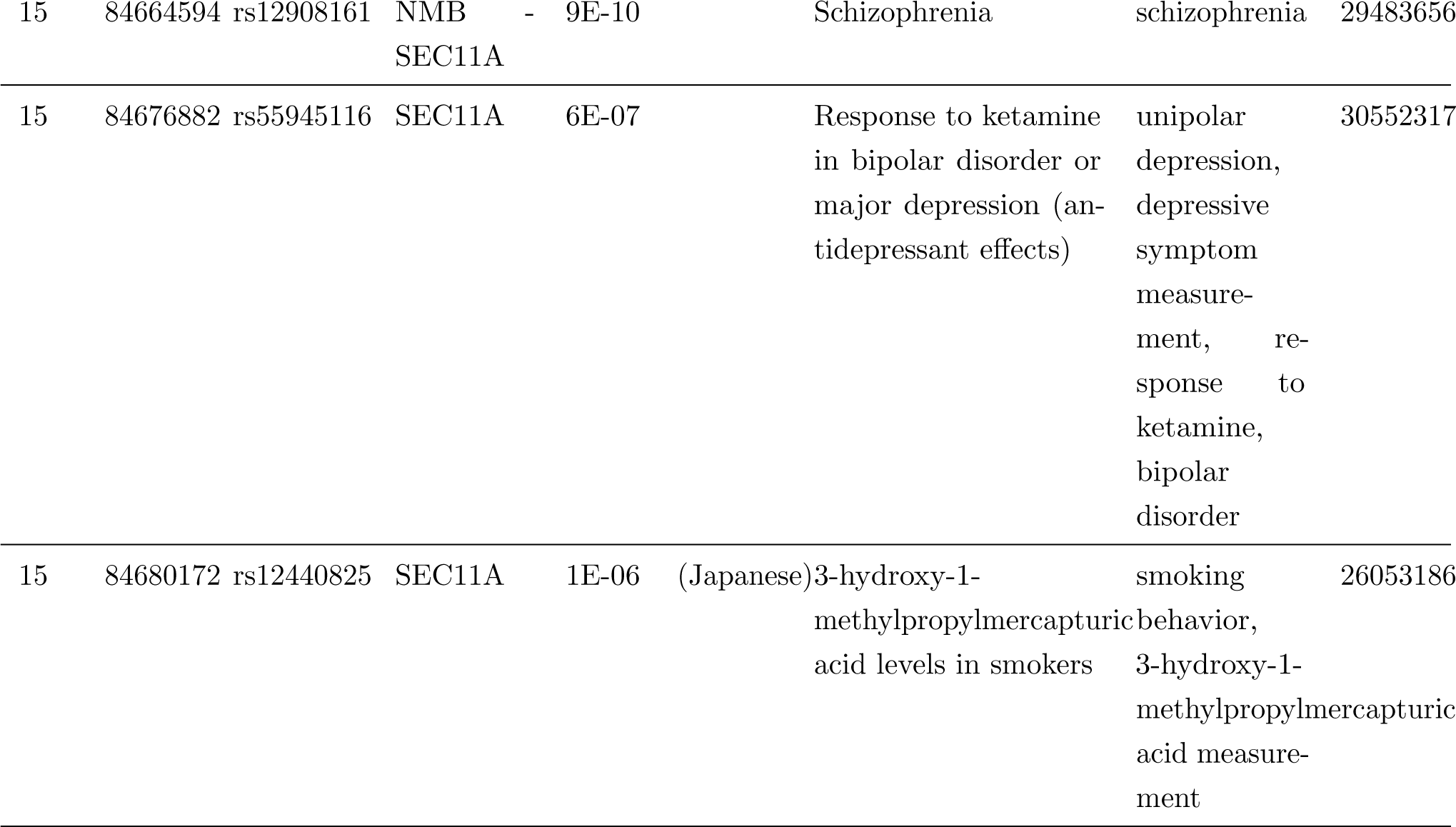
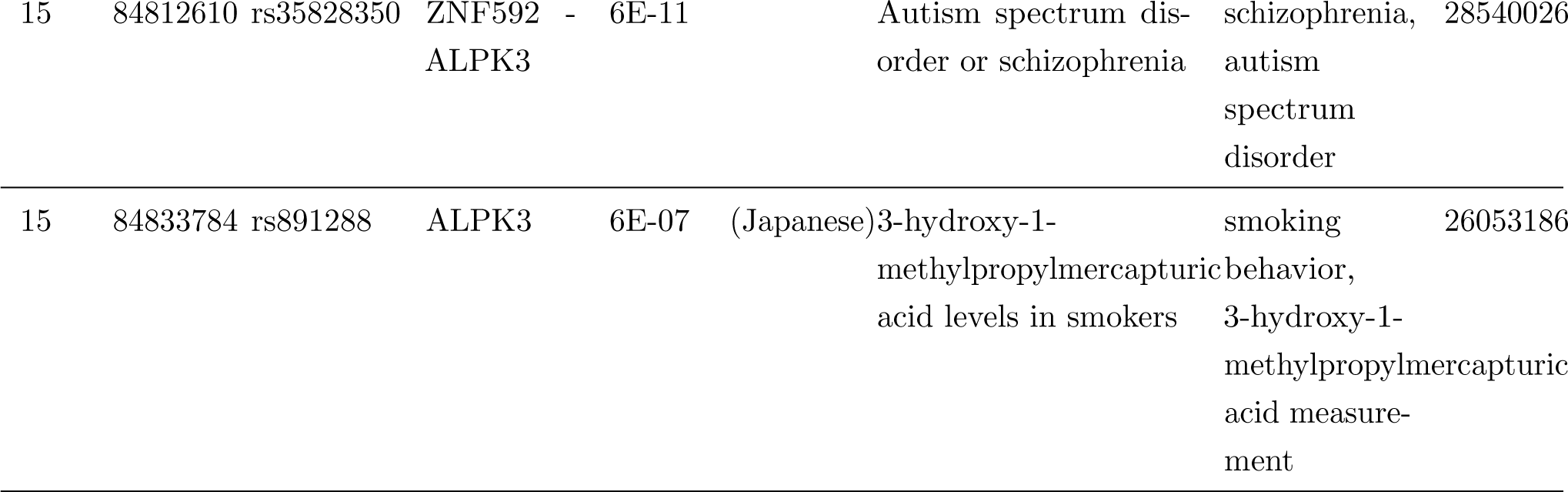
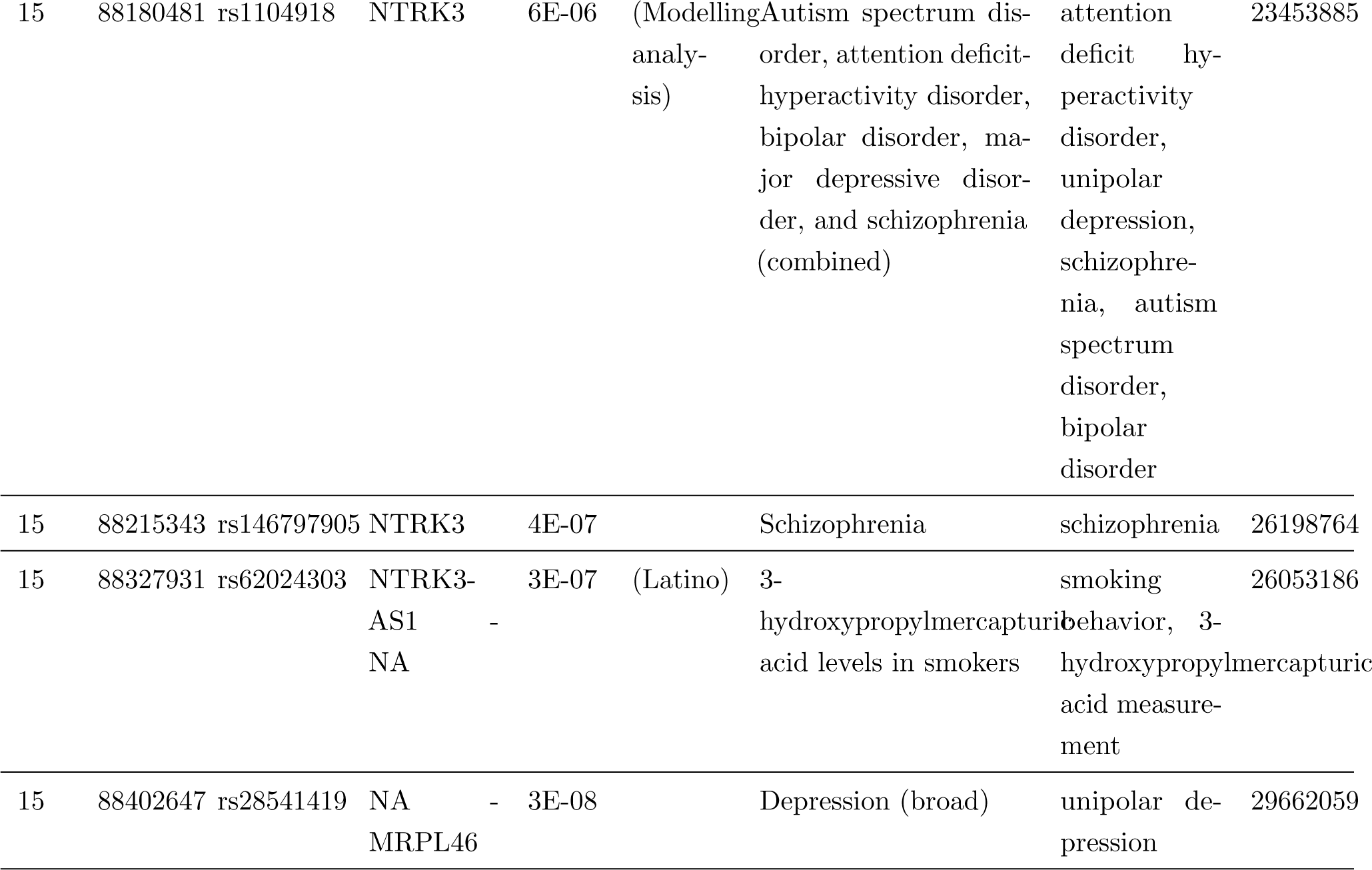
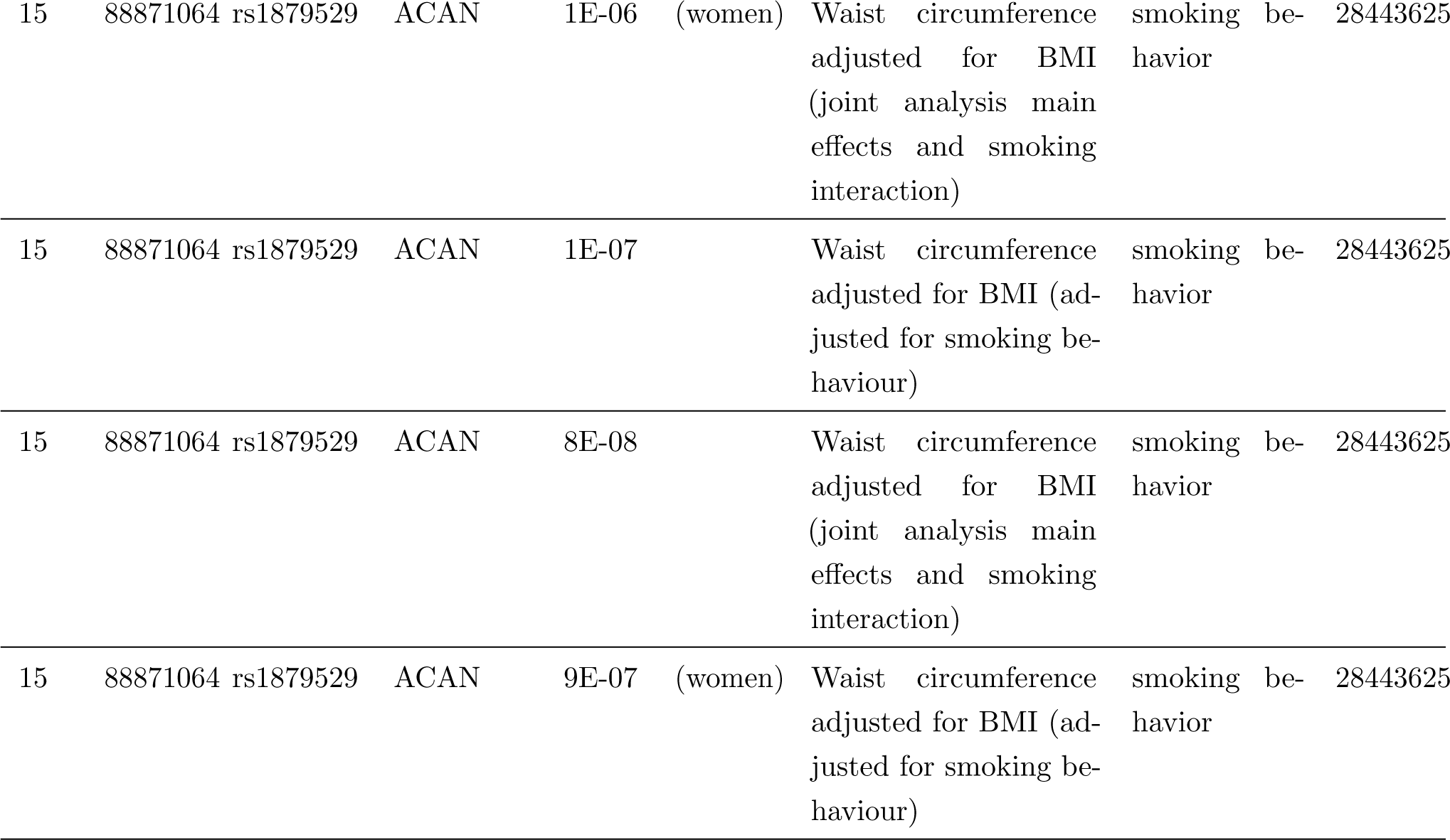
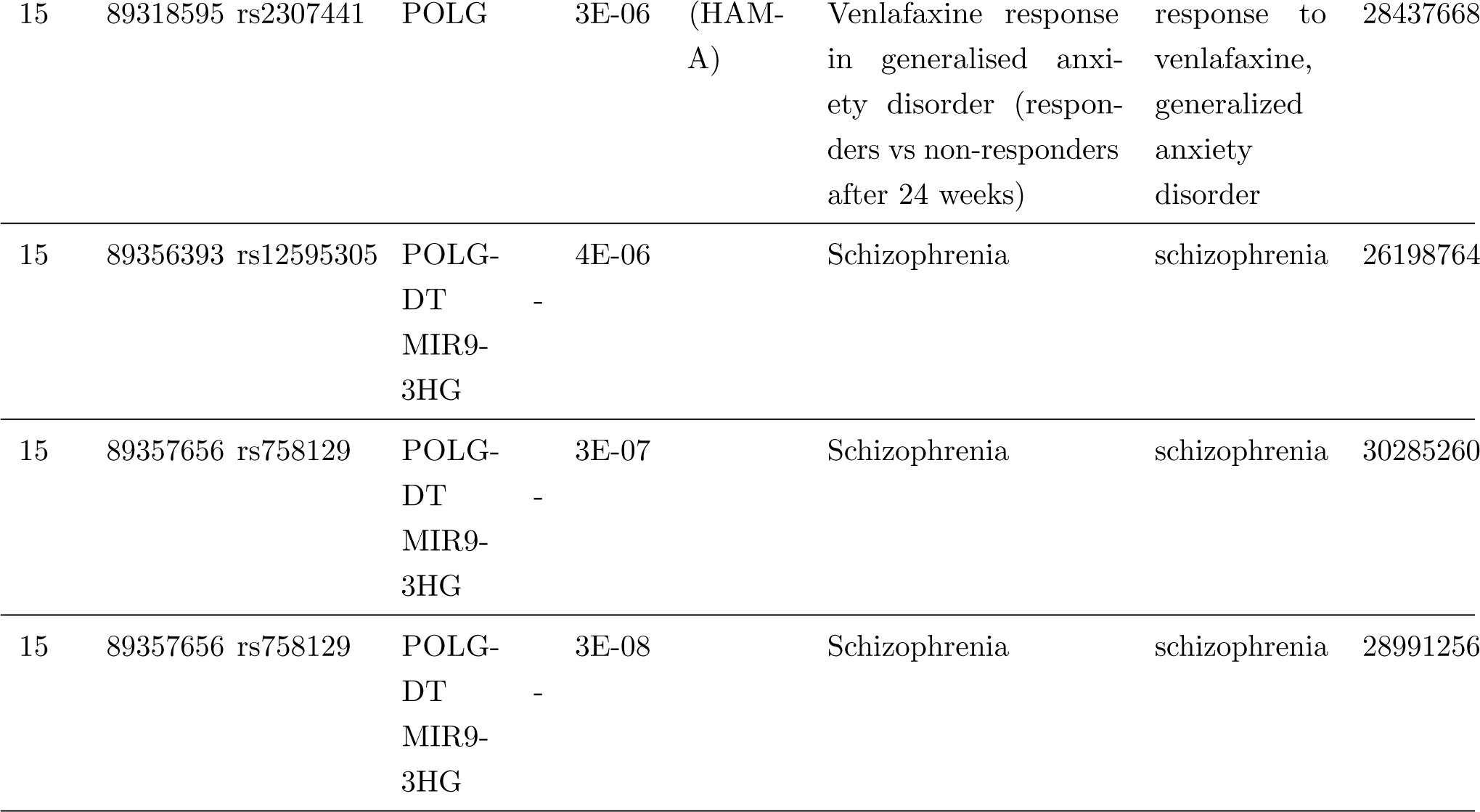
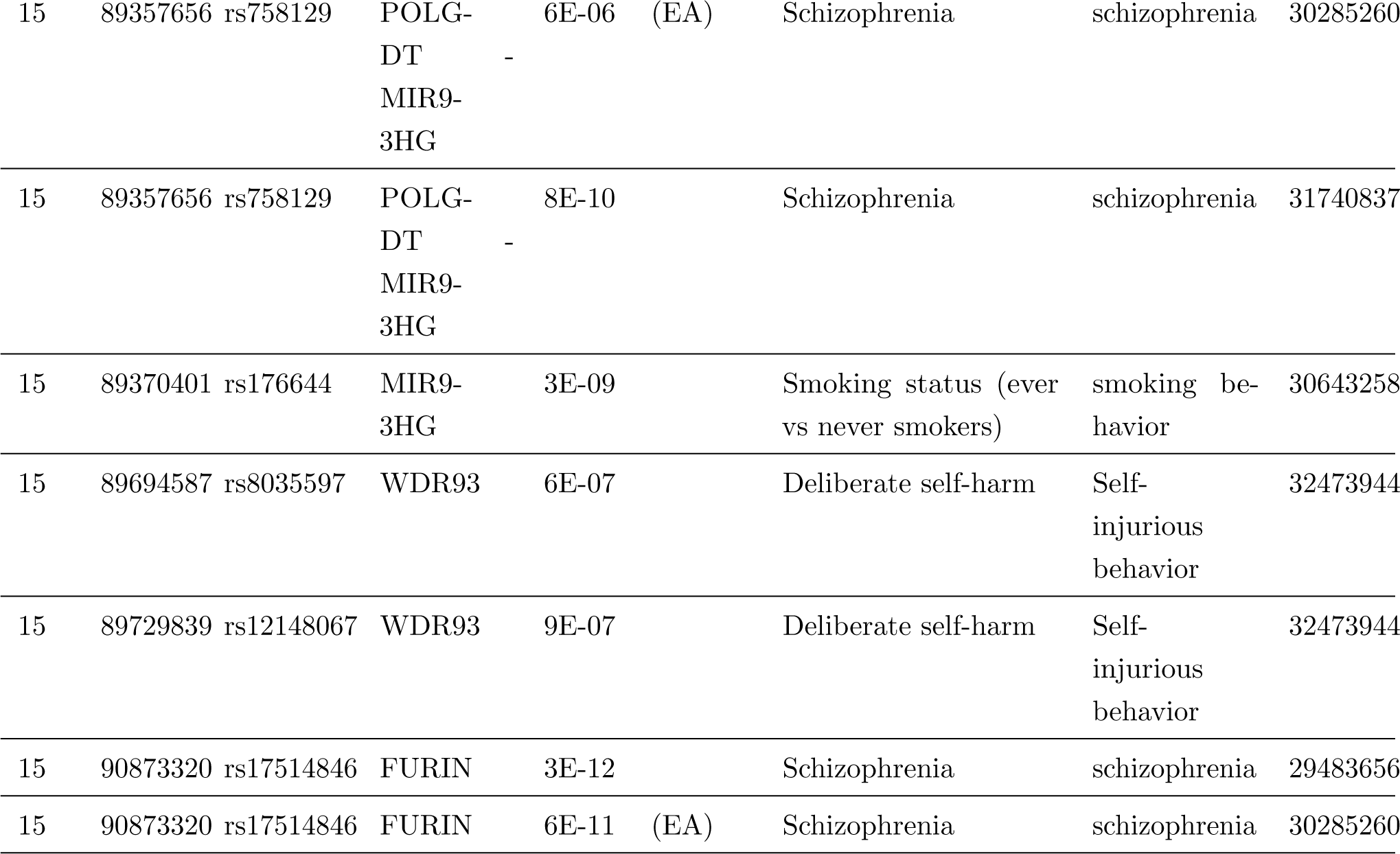
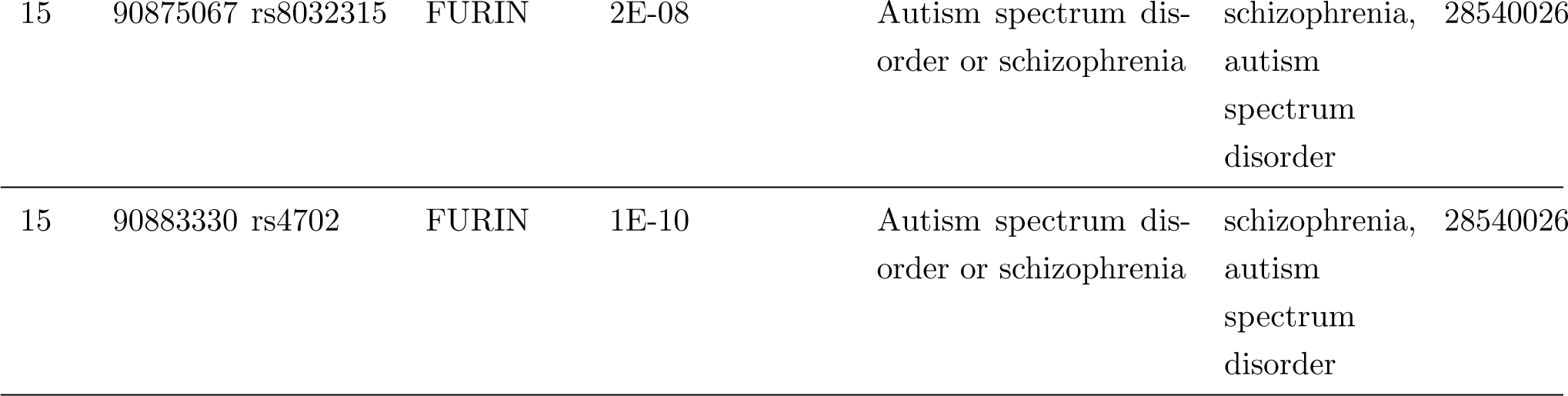
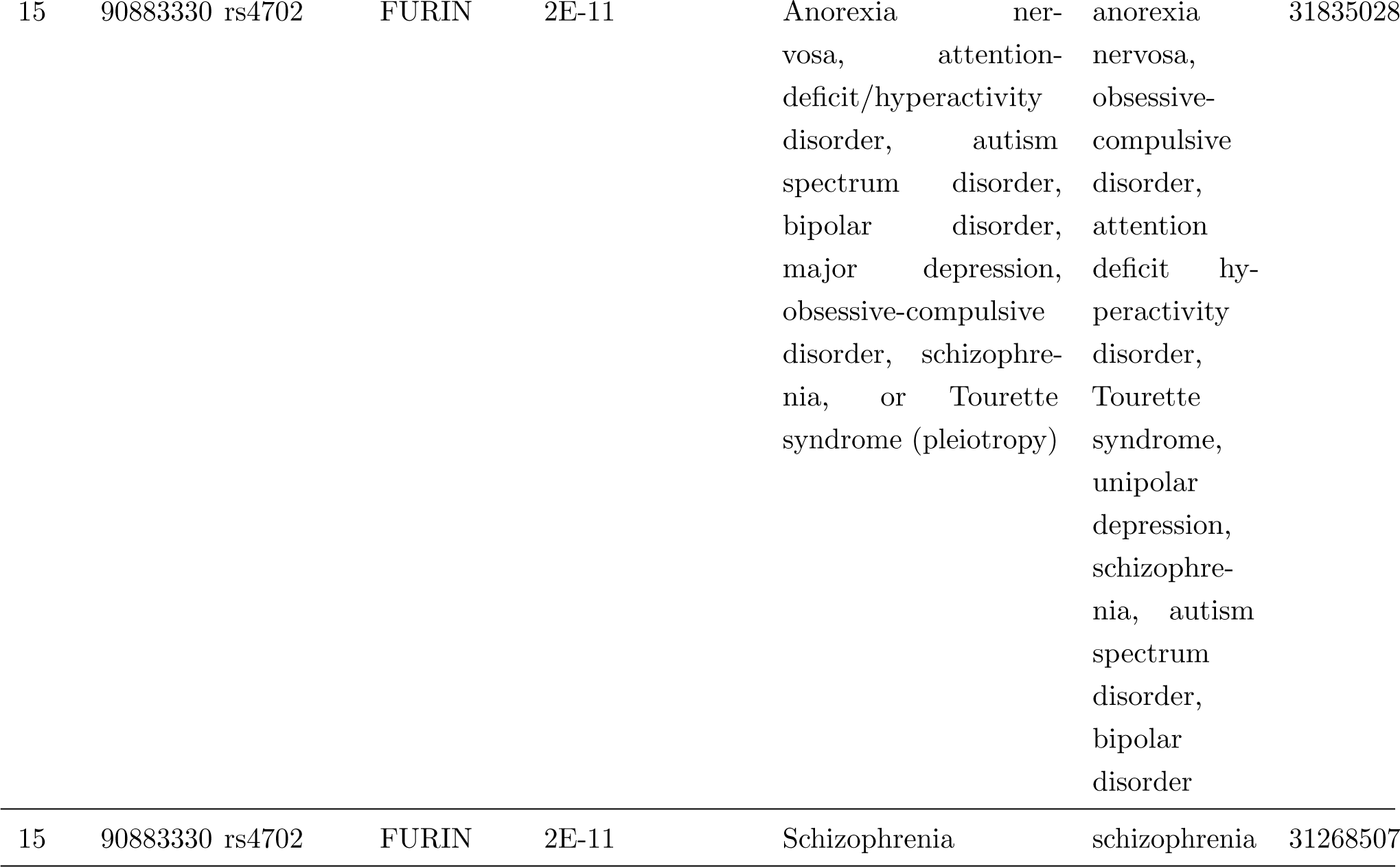
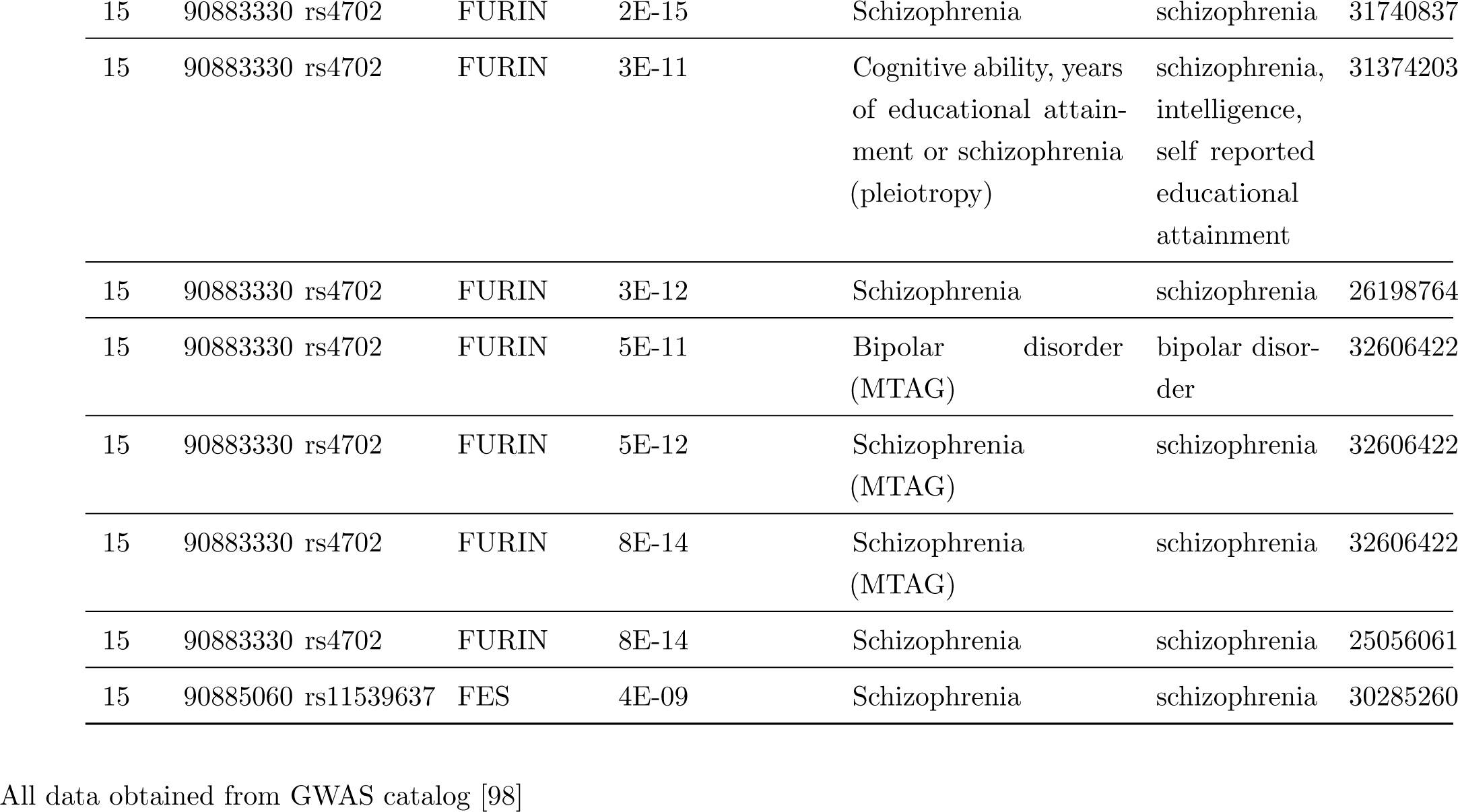
Hot spot on rat Chr1 144Mb loci has 17 genes associated psychiatric conditions in human

**Figure S1:**
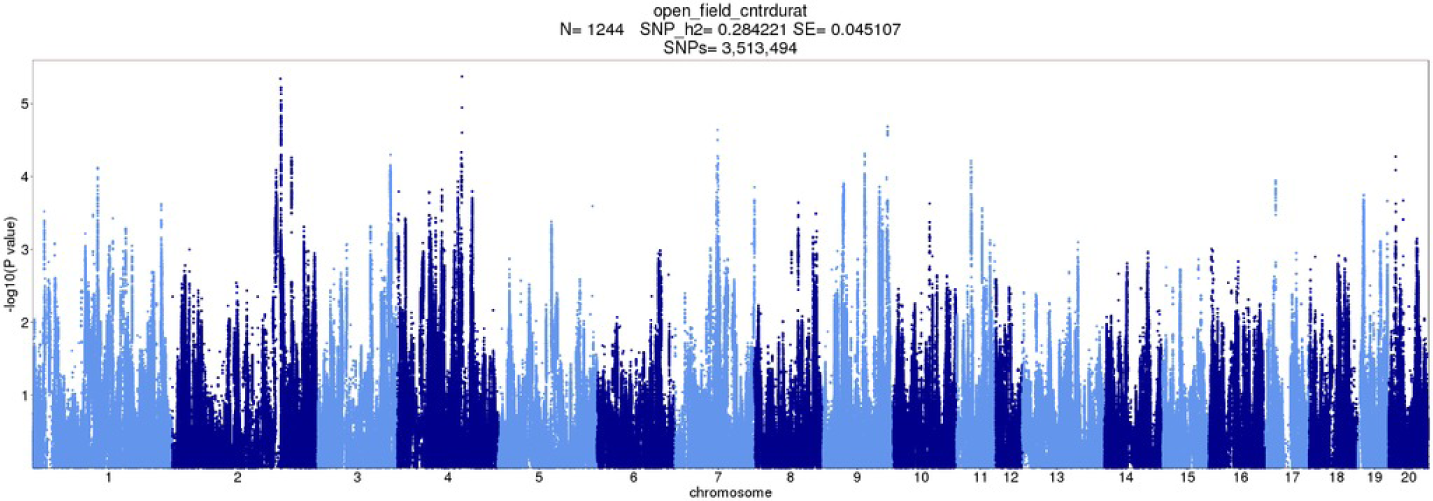
OFT GWAS: Duration in center zone, mean

**Figure S2:**
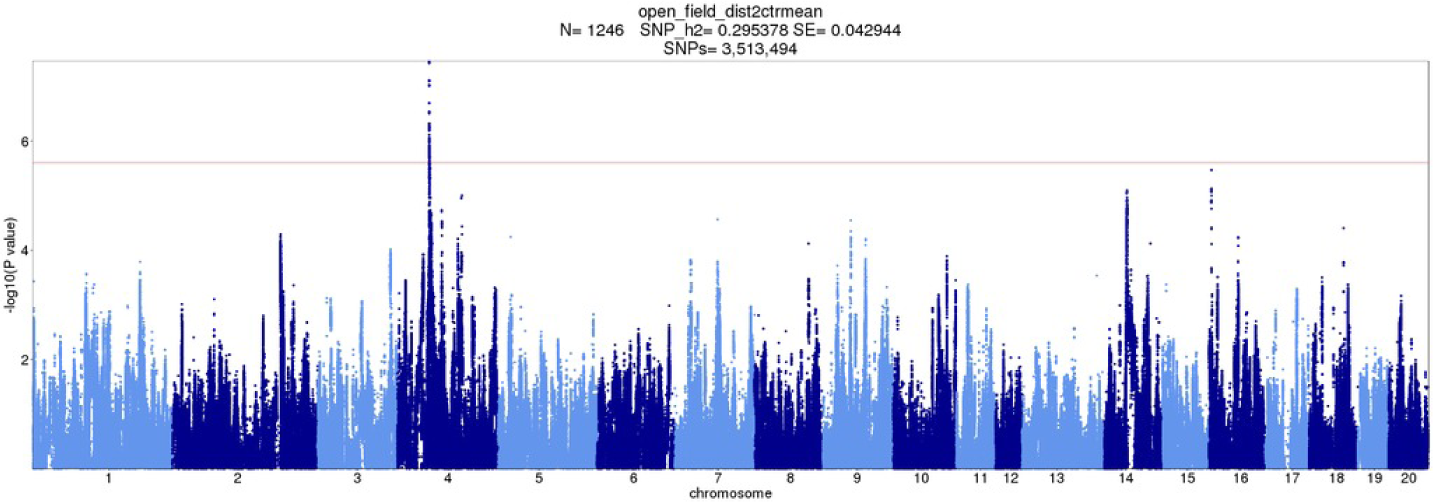
OFT GWAS: Mean distance to center zone

**Figure S3:**
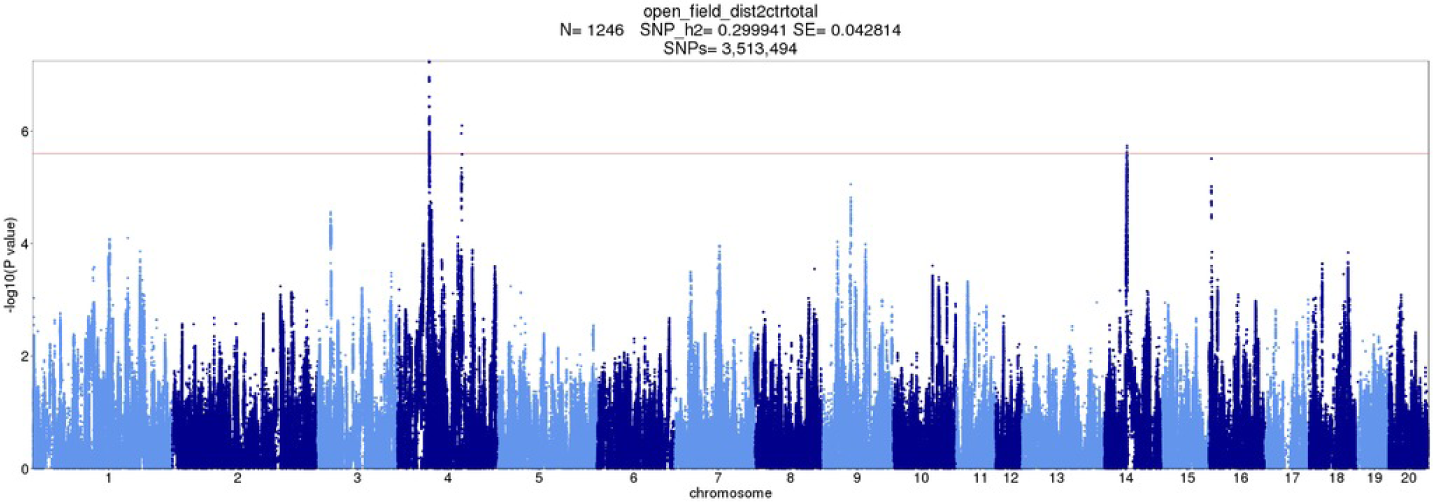
OFT GWAS: Total distance to center zone

**Figure S4:**
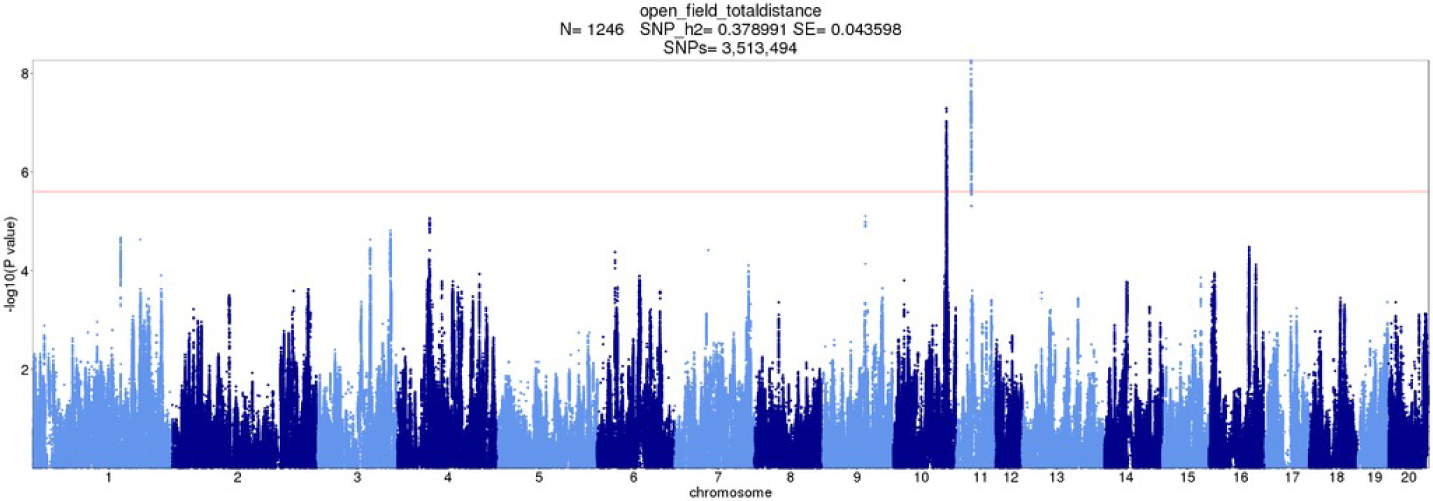
OFT GWAS: Total travel distance

**Figure S5:**
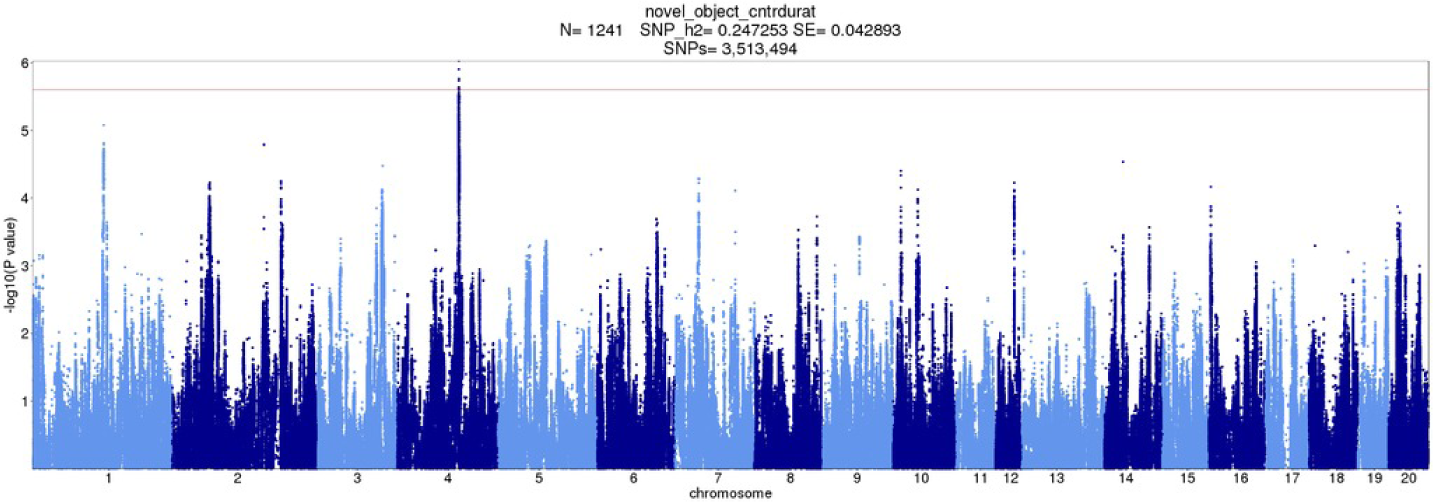
NOIT GWAS: Duration in center zone

**Figure S6:**
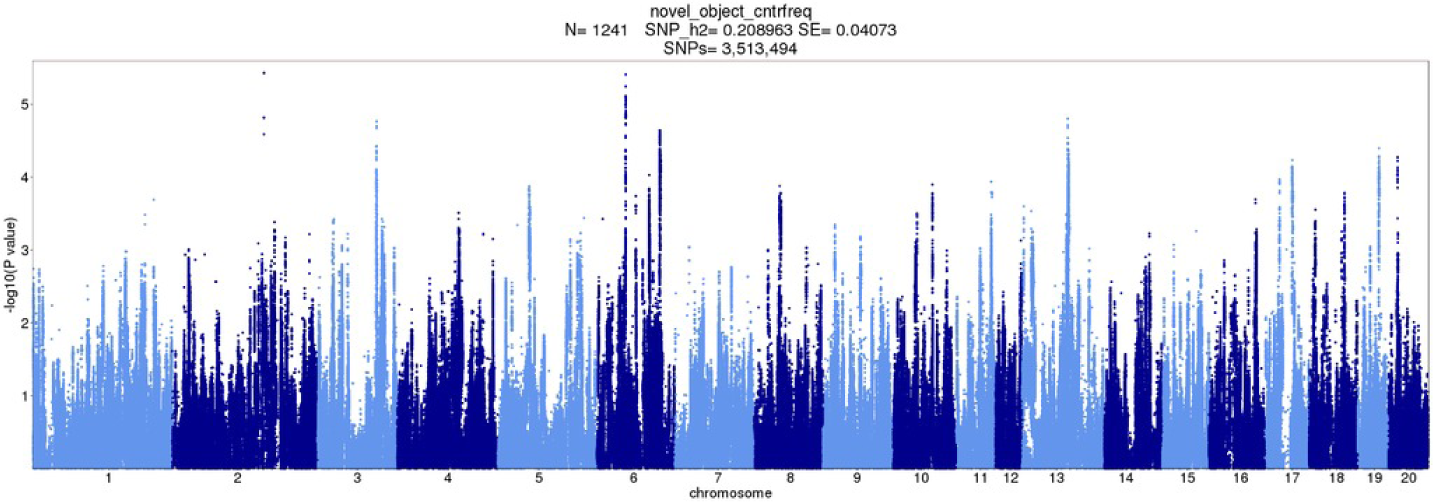
NOIT GWAS: Frequency of entering center zone

**Figure S7:**
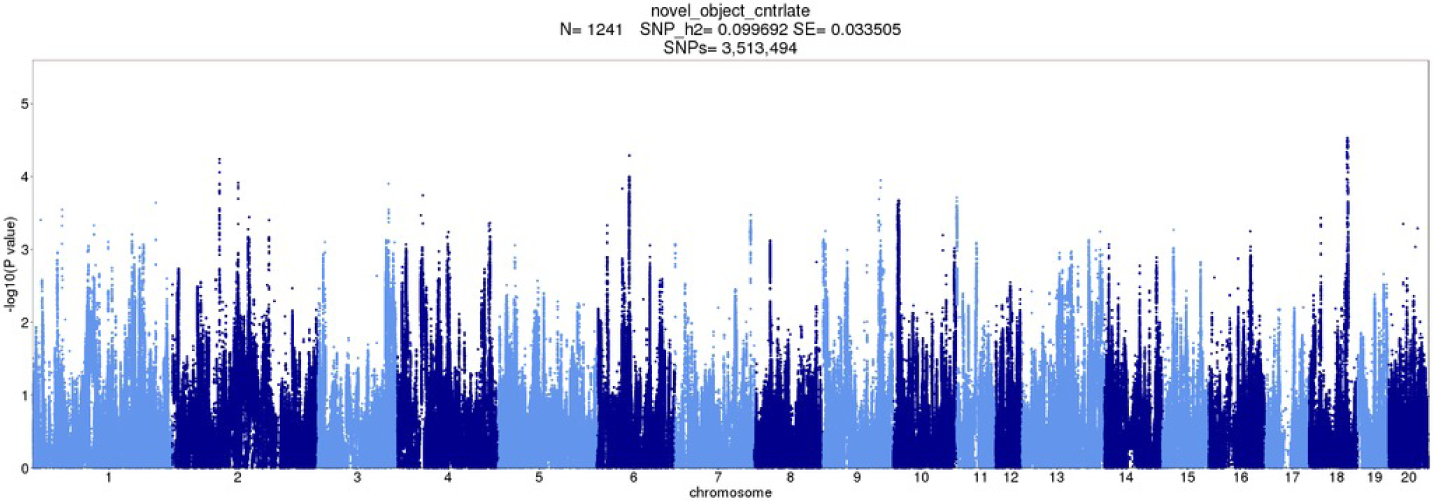
NOIT GWAS: Latency of entering center zone

**Figure S8:**
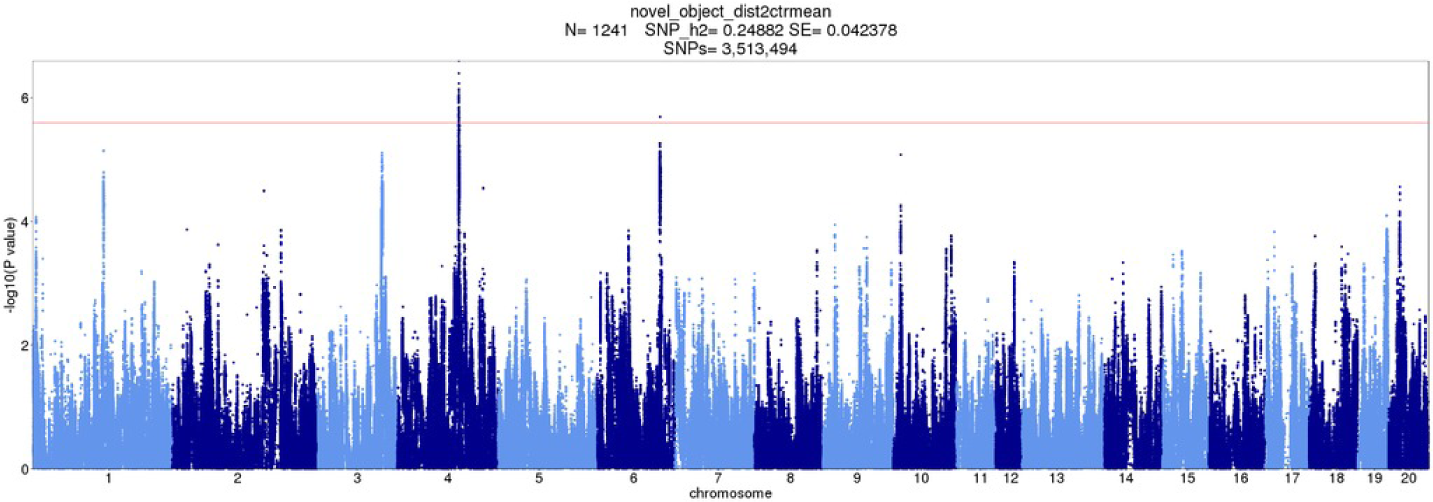
NOIT GWAS: Mean distance to center zone

**Figure S9:**
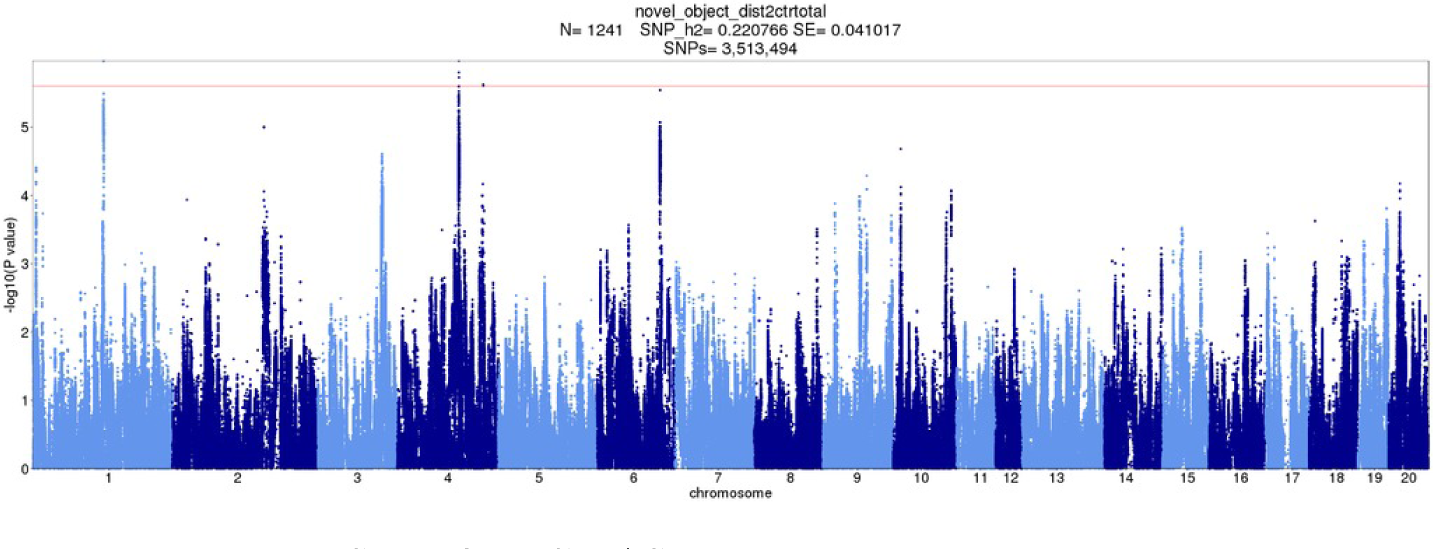
NOIT GWAS: Total distance to center zone

**Figure S10:**
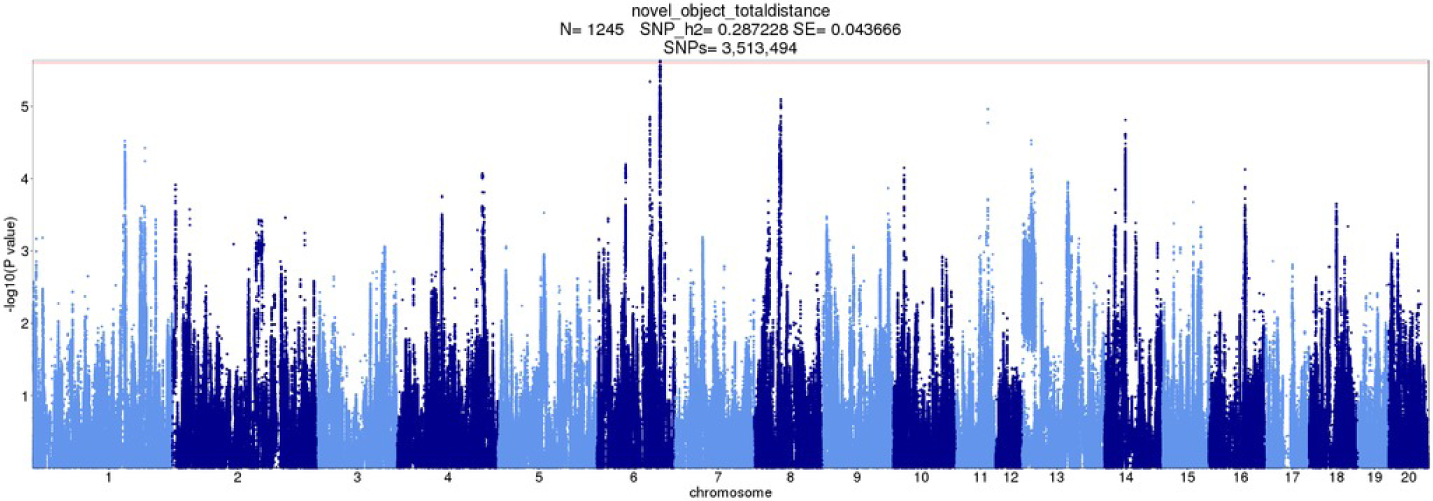
NOIT GWAS: Total travel distance

**Figure S11:**
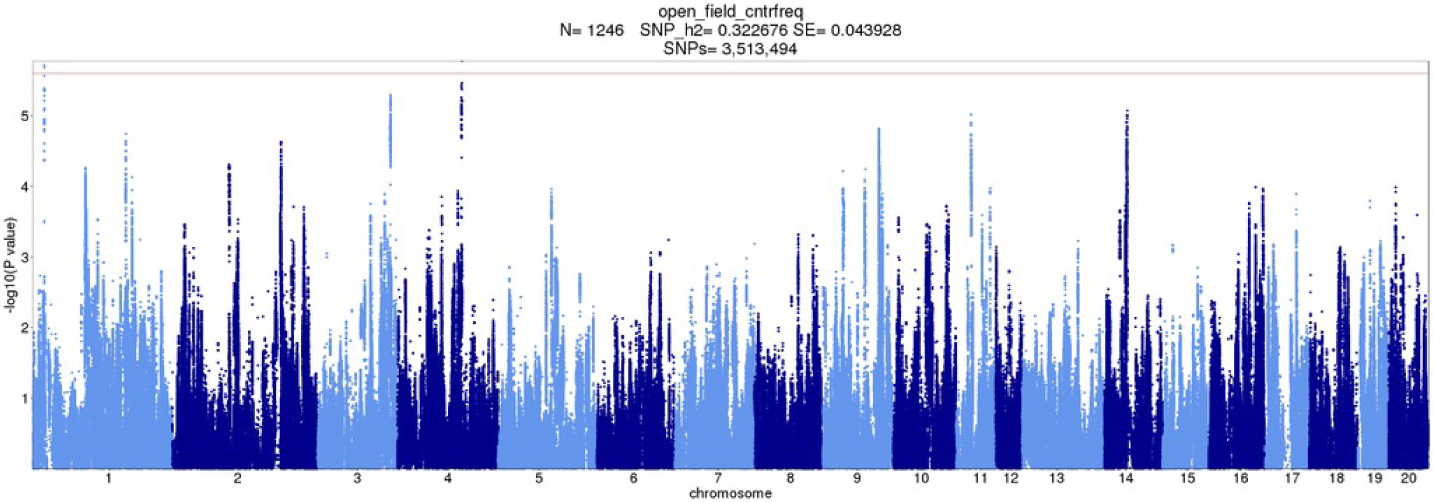
NOIT GWAS: Frequency of entering center zone, mean

**Figure S12:**
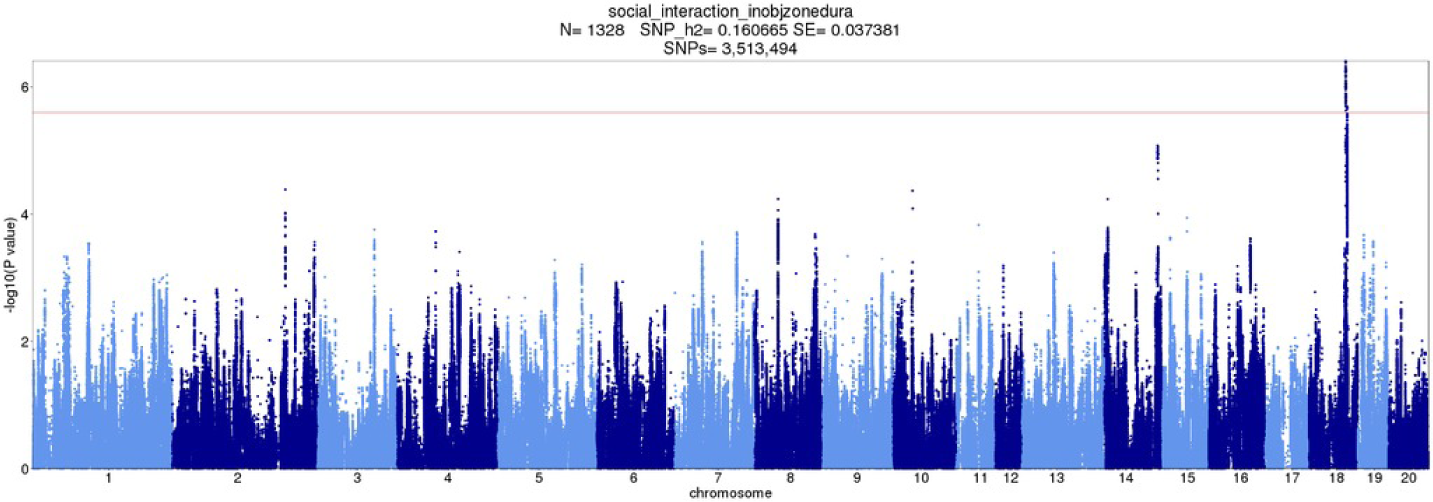
SIT GWAS: Duration in object zone

**Figure S13:**
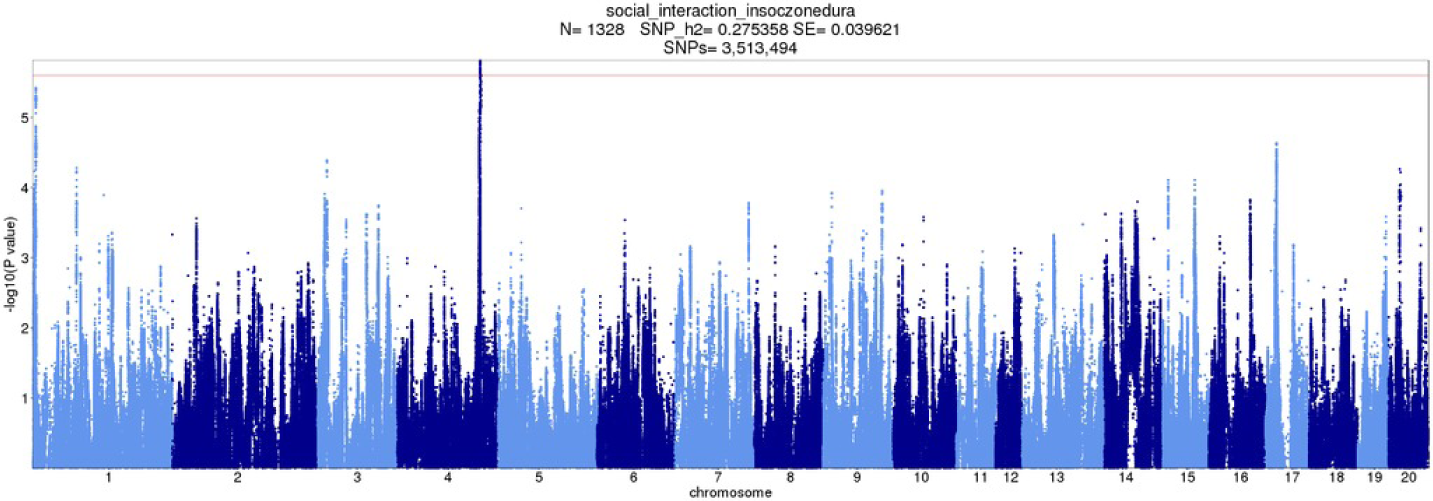
SIT GWAS: Duration in social zone

**Figure S14:**
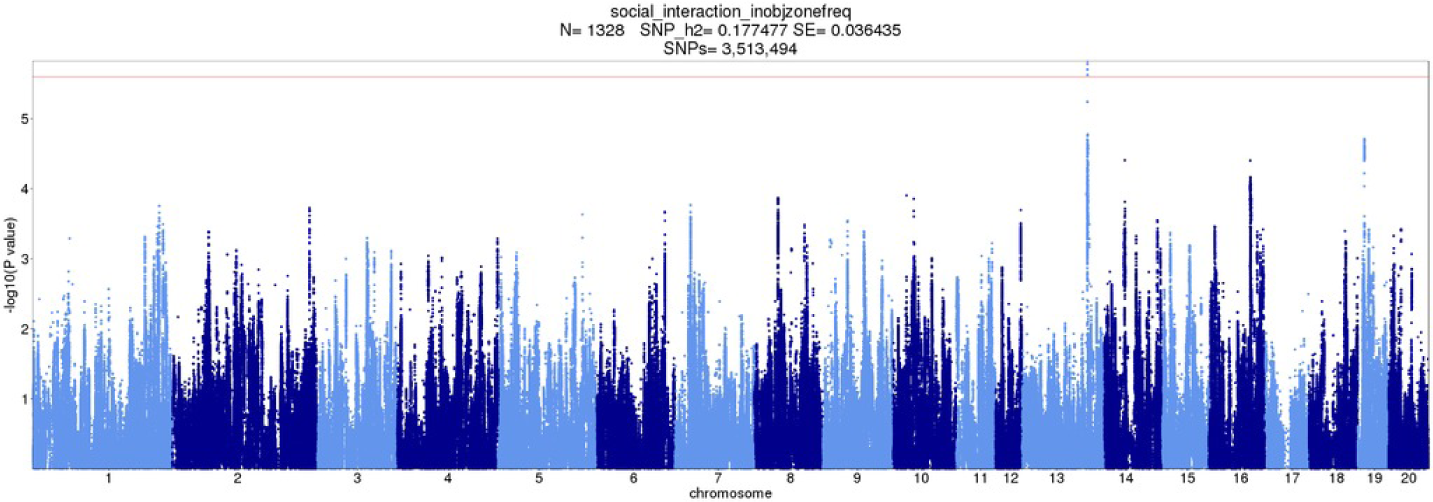
SIT GWAS: Frequency of entering object zone

**Figure S15:**
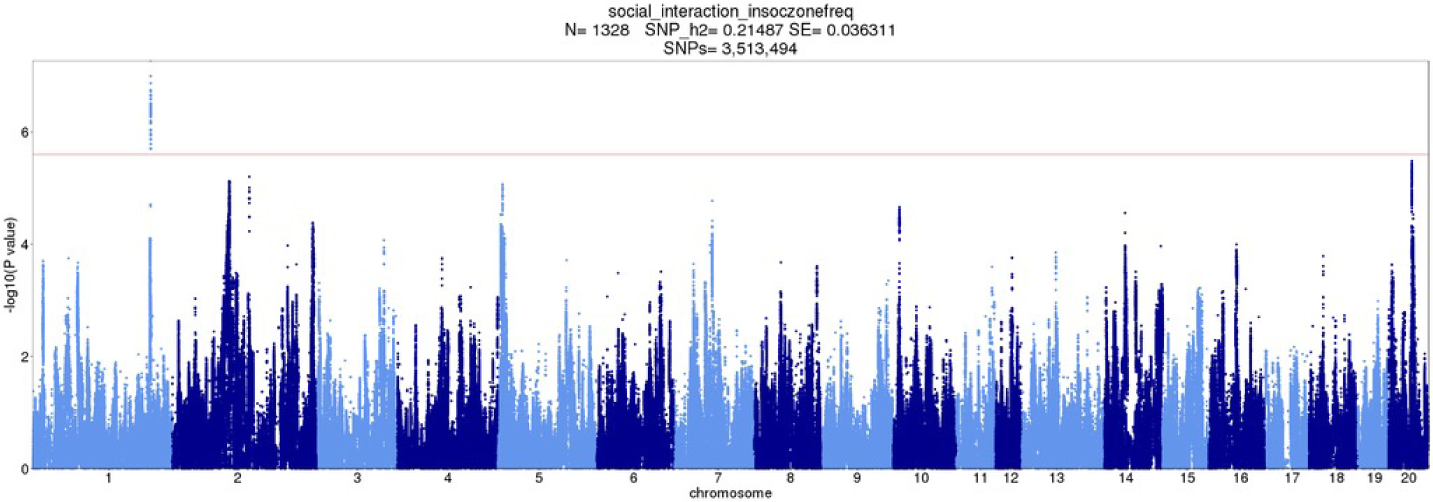
SIT GWAS: Frequency of entering social zone

**Figure S16:**
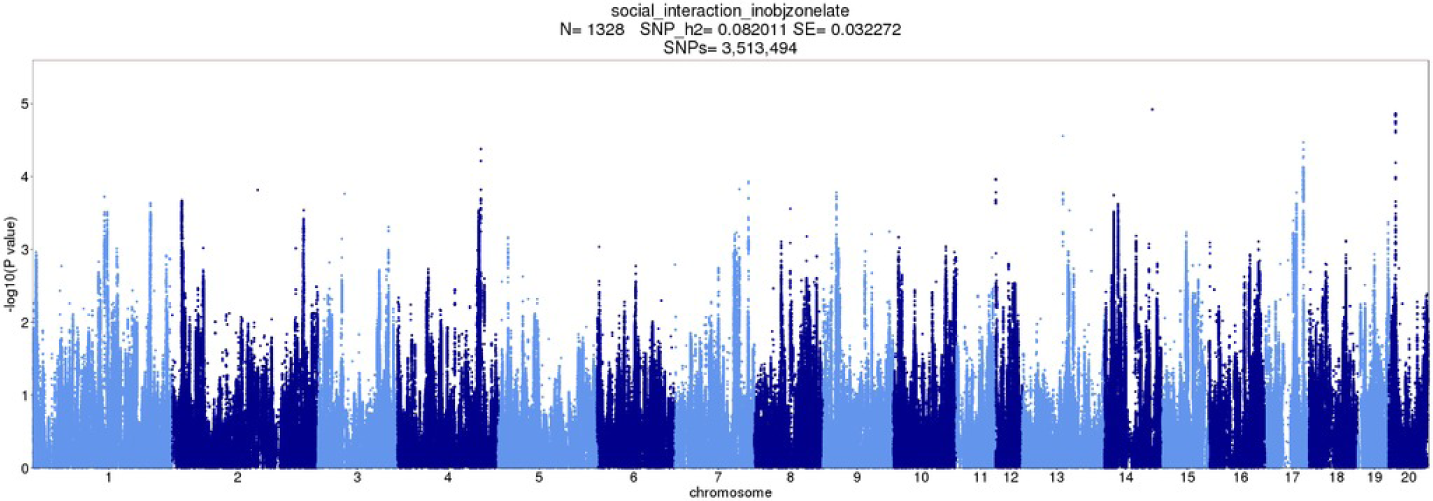
SIT GWAS: Latency of entering object zone

**Figure S17:**
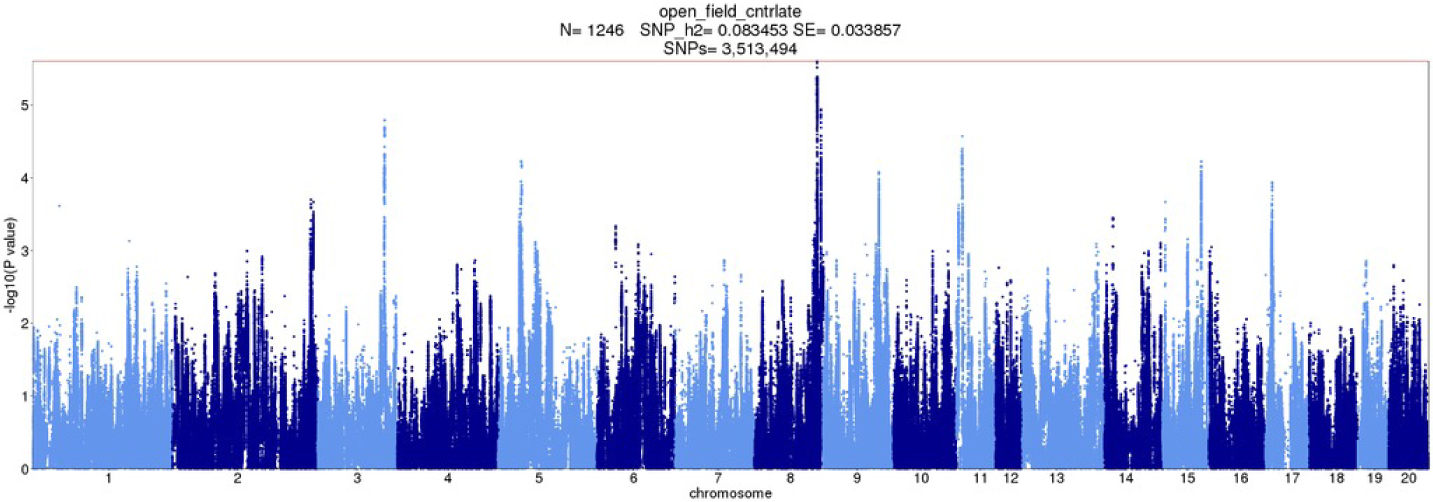
SIT GWAS: Latency of entering center zone, mean

**Figure S18:**
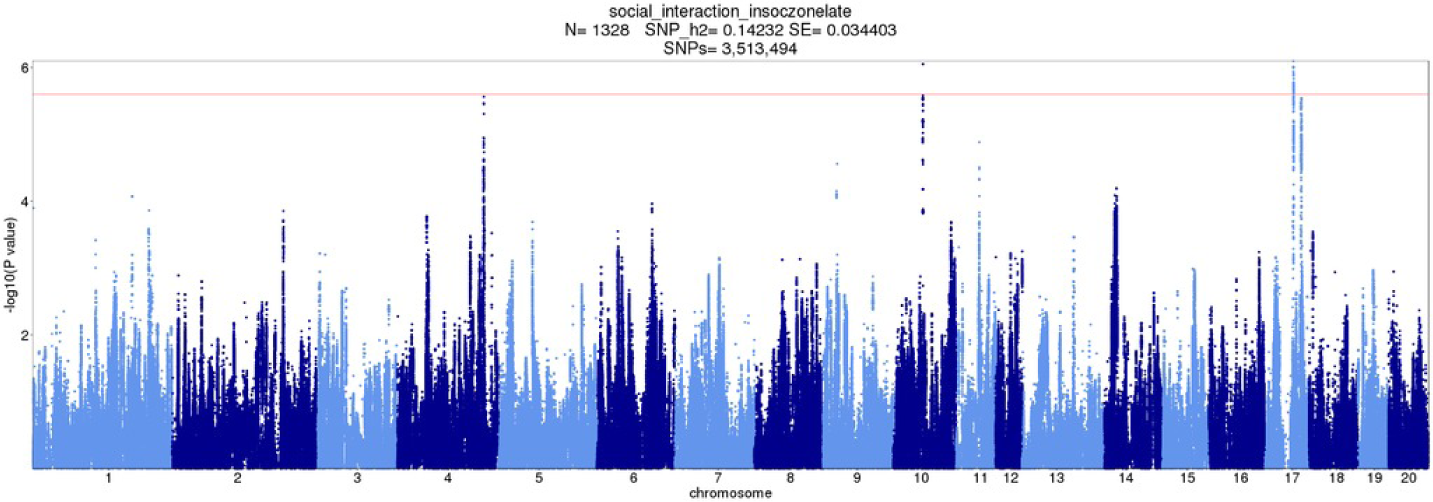
SIT GWAS: Latency of entering social zone

**Figure S19:**
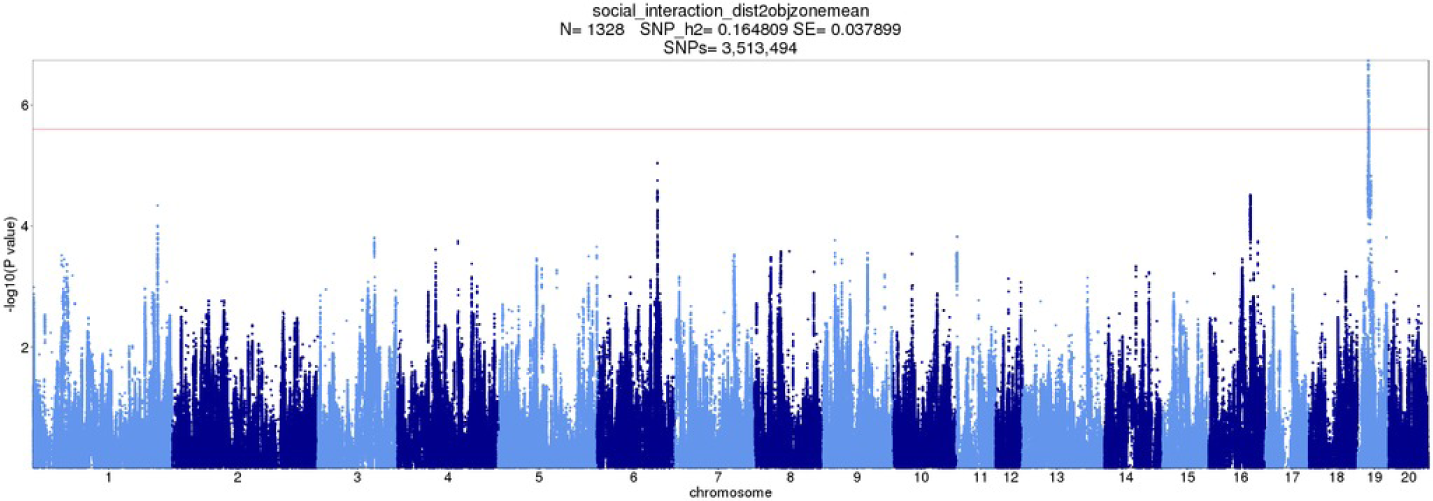
SIT GWAS: Mean distance to object zone

**Figure S20:**
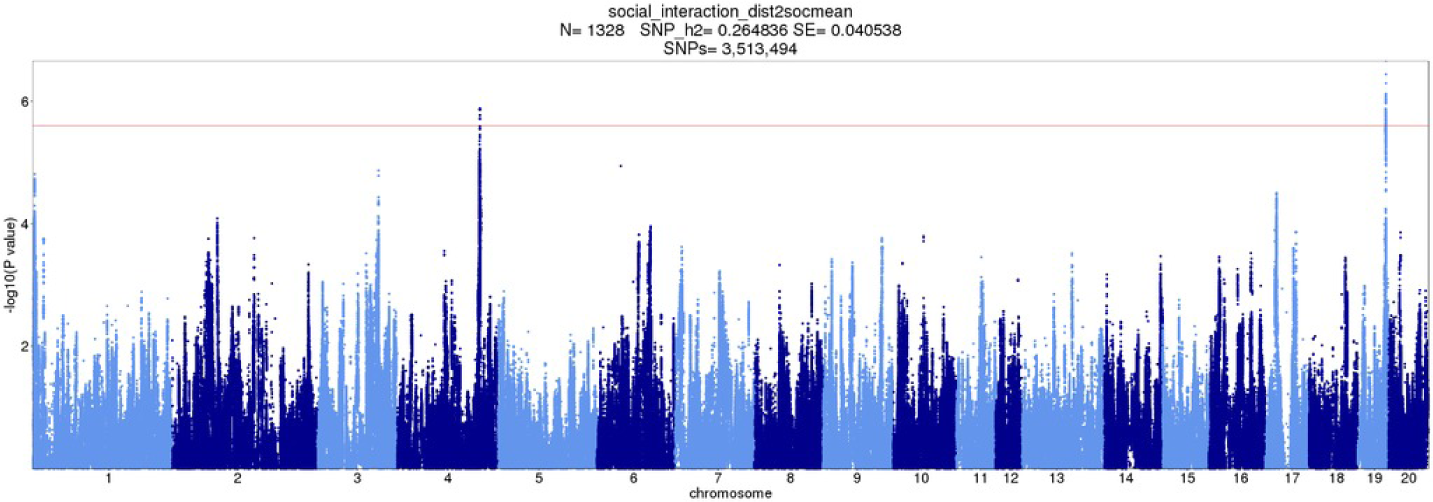
SIT GWAS: Mean distance to social zone

**Figure S21:**
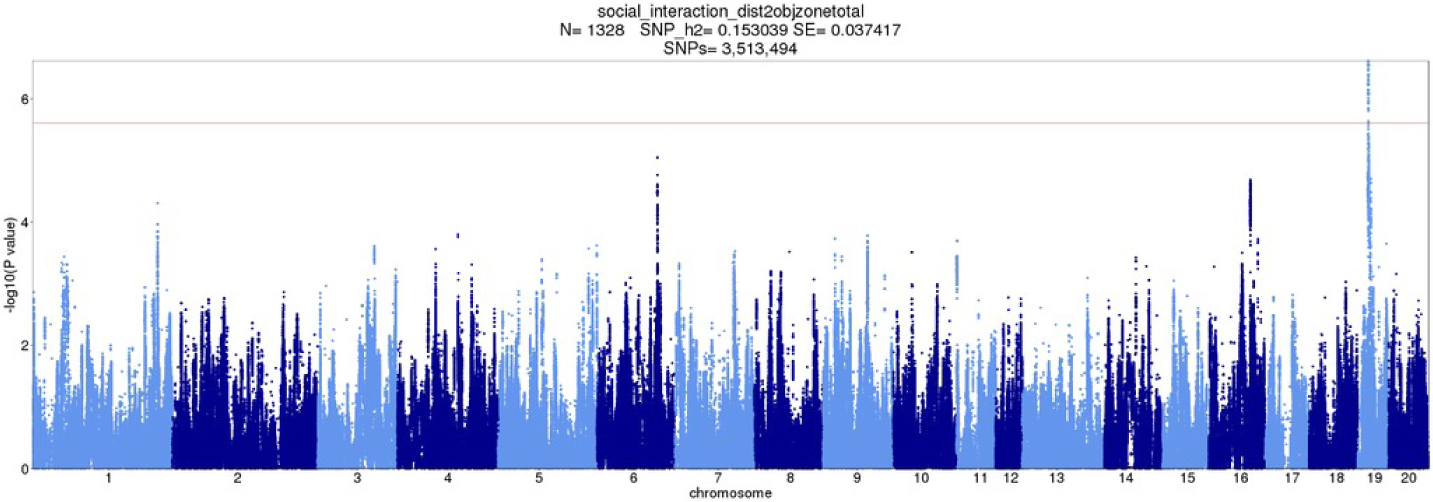
SIT GWAS: Total distance to object zone

**Figure S22:**
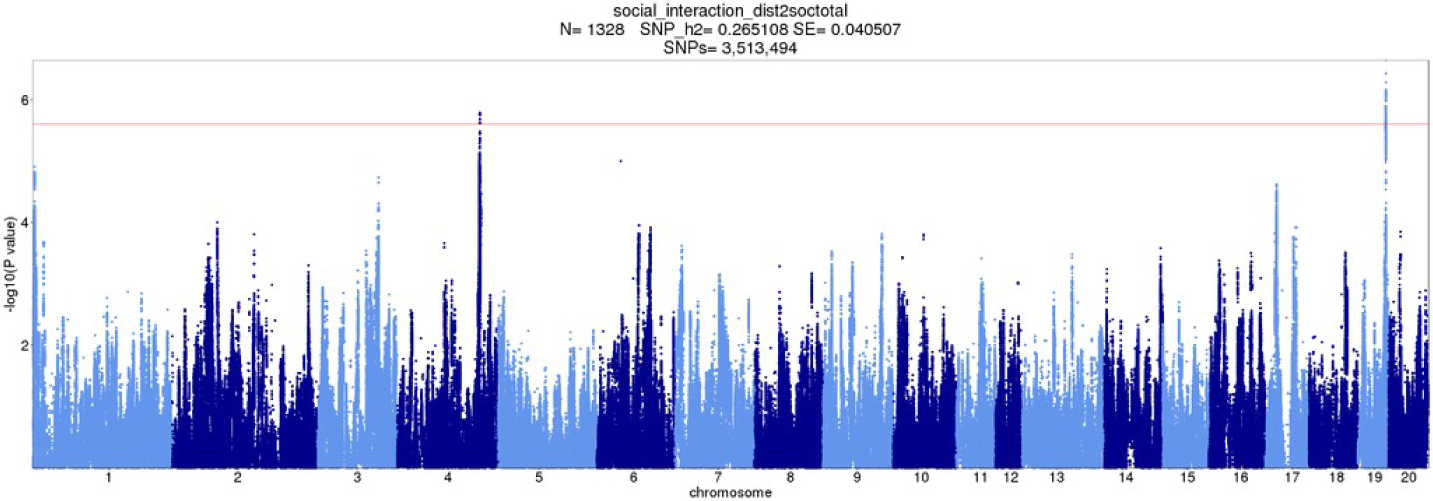
SIT GWAS: Total distance to social zone

**Figure S23:**
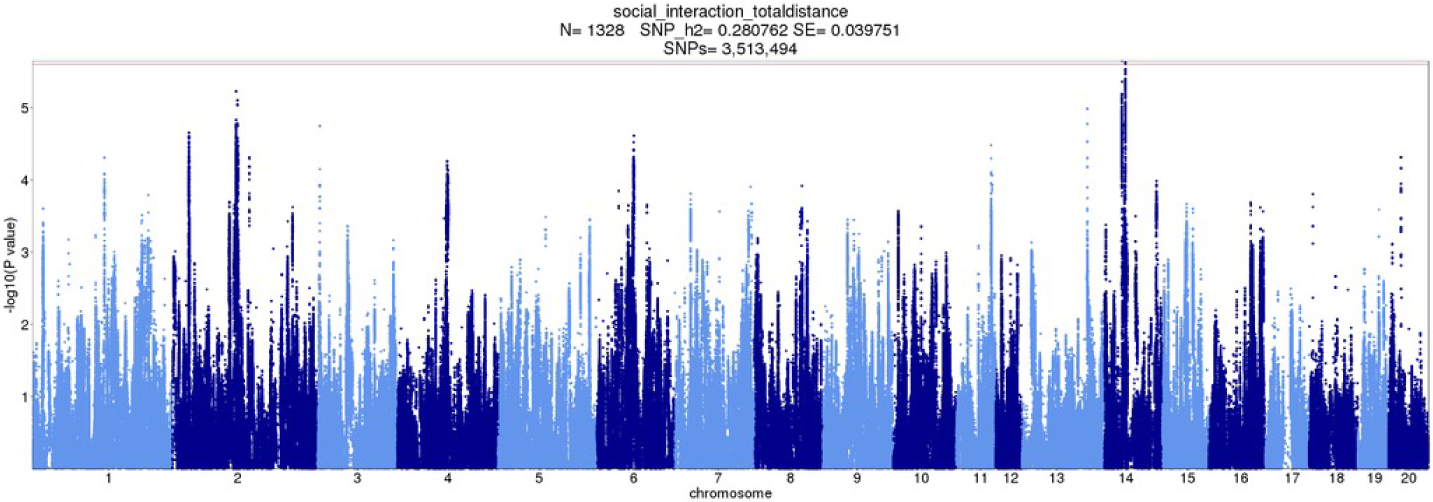
SIT GWAS: Total travel distance

**Figure S24:**
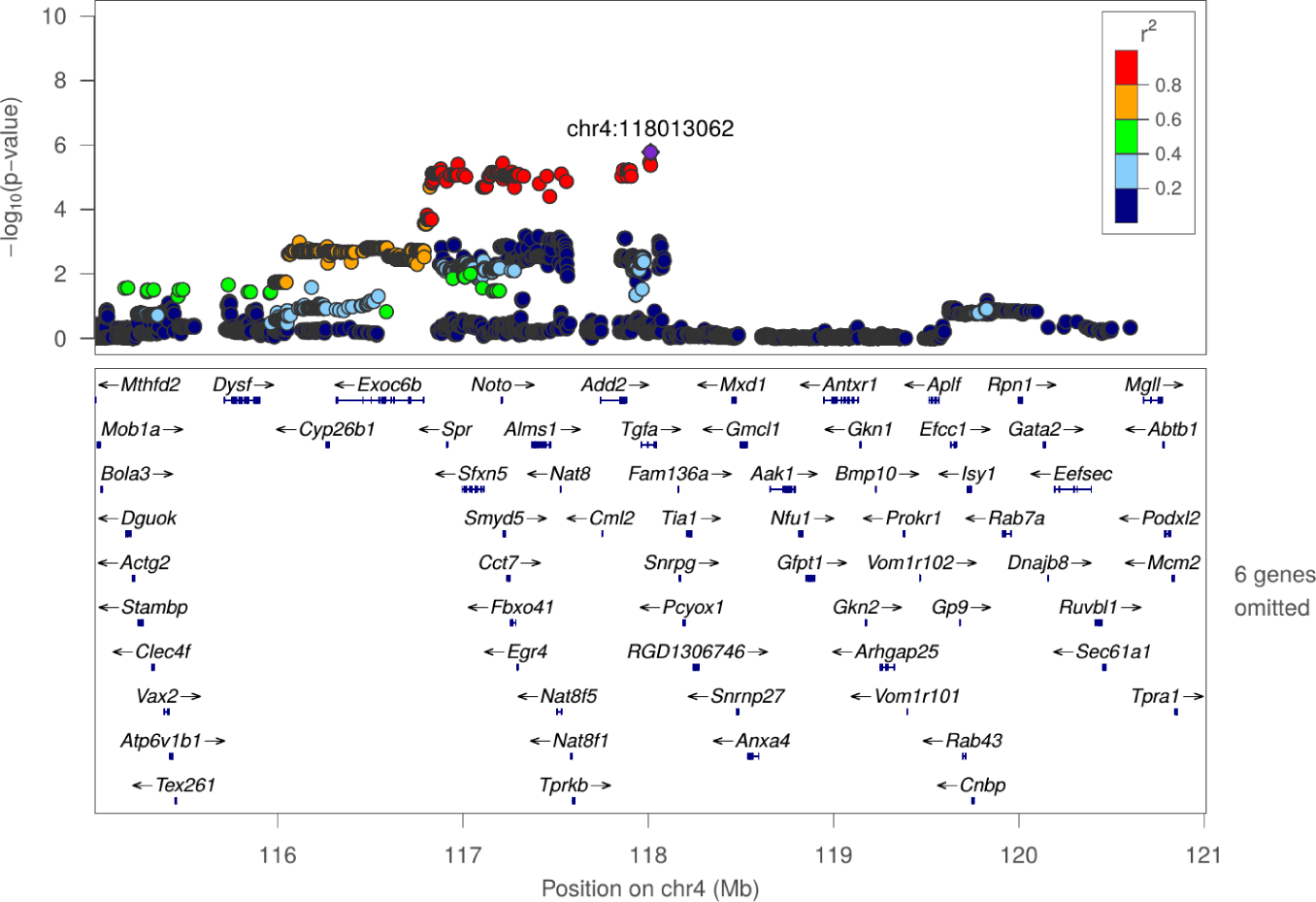
Regional association plot for OFT: Frequency of entering center zone at chr4:118013062

**Figure S25:**
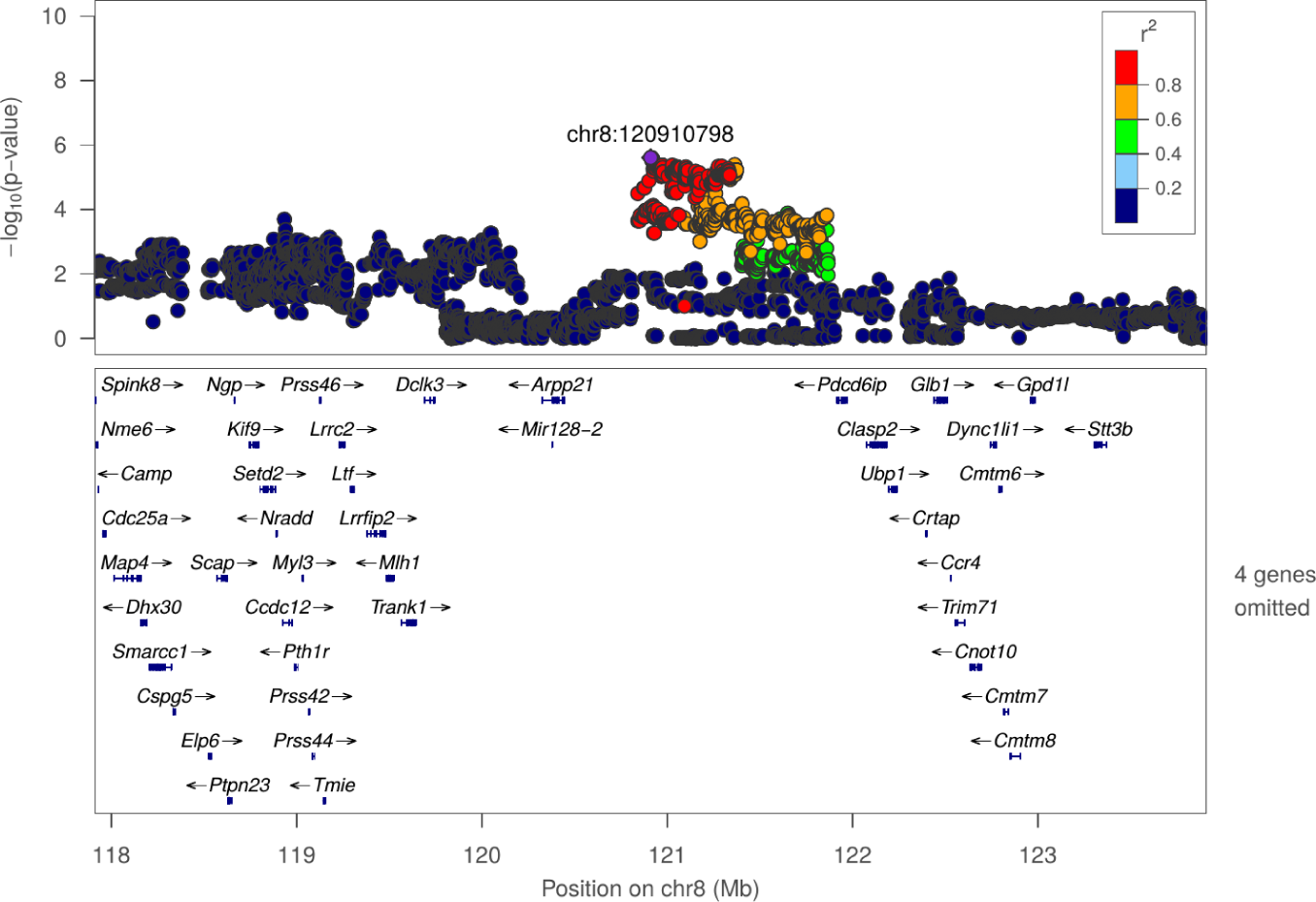
Regional association plot for OFT: Latency of entering center zone at chr8:120910798

**Figure S26:**
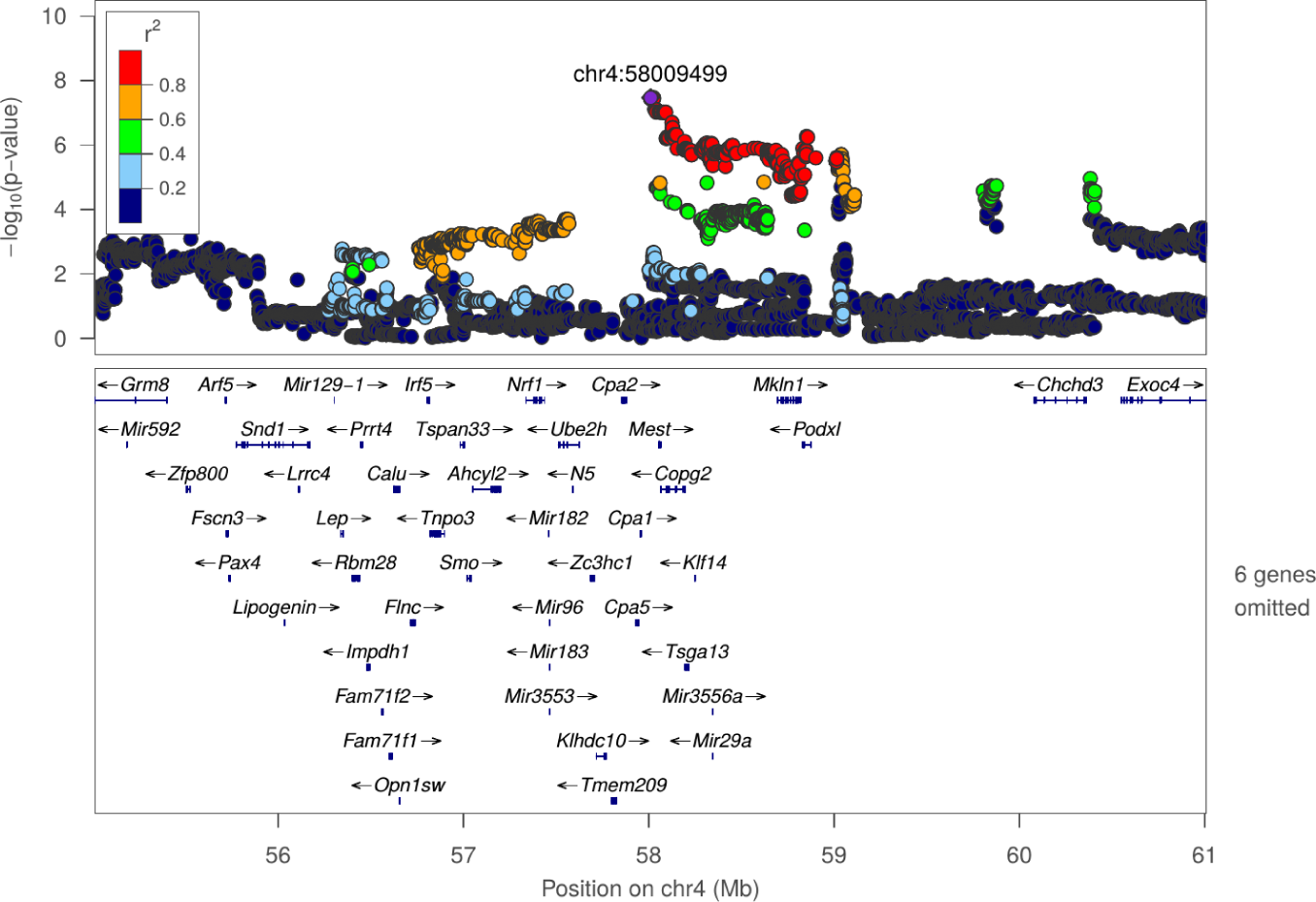
Regional association plot for OFT: Mean distance to center zone at chr4:58009499

**Figure S27:**
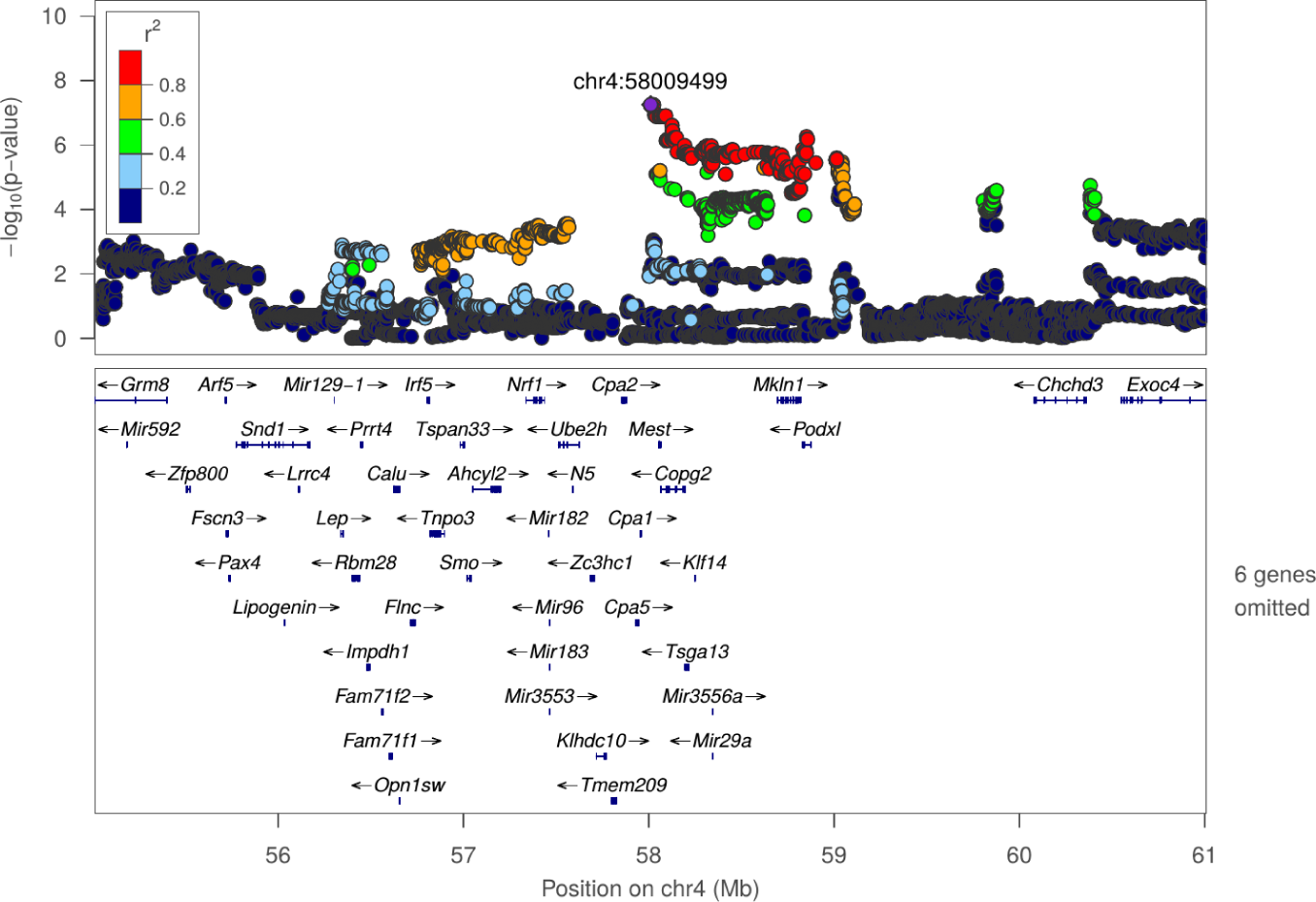
Regional association plot for OFT: Total distance to center zone at chr4:58009499

**Figure S28:**
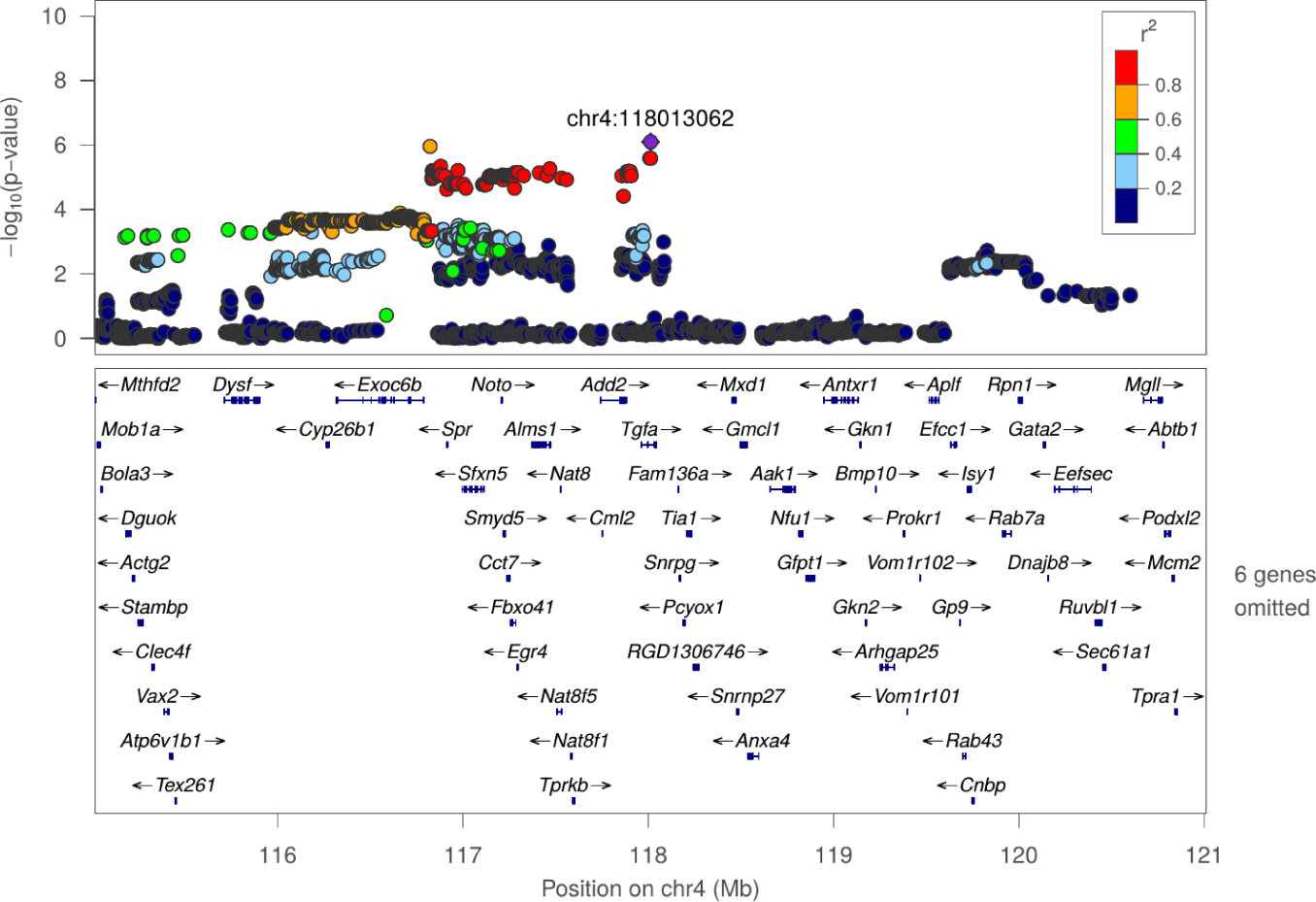
Regional association plot for OFT: Total distance to center zone at chr4:118013062

**Figure S29:**
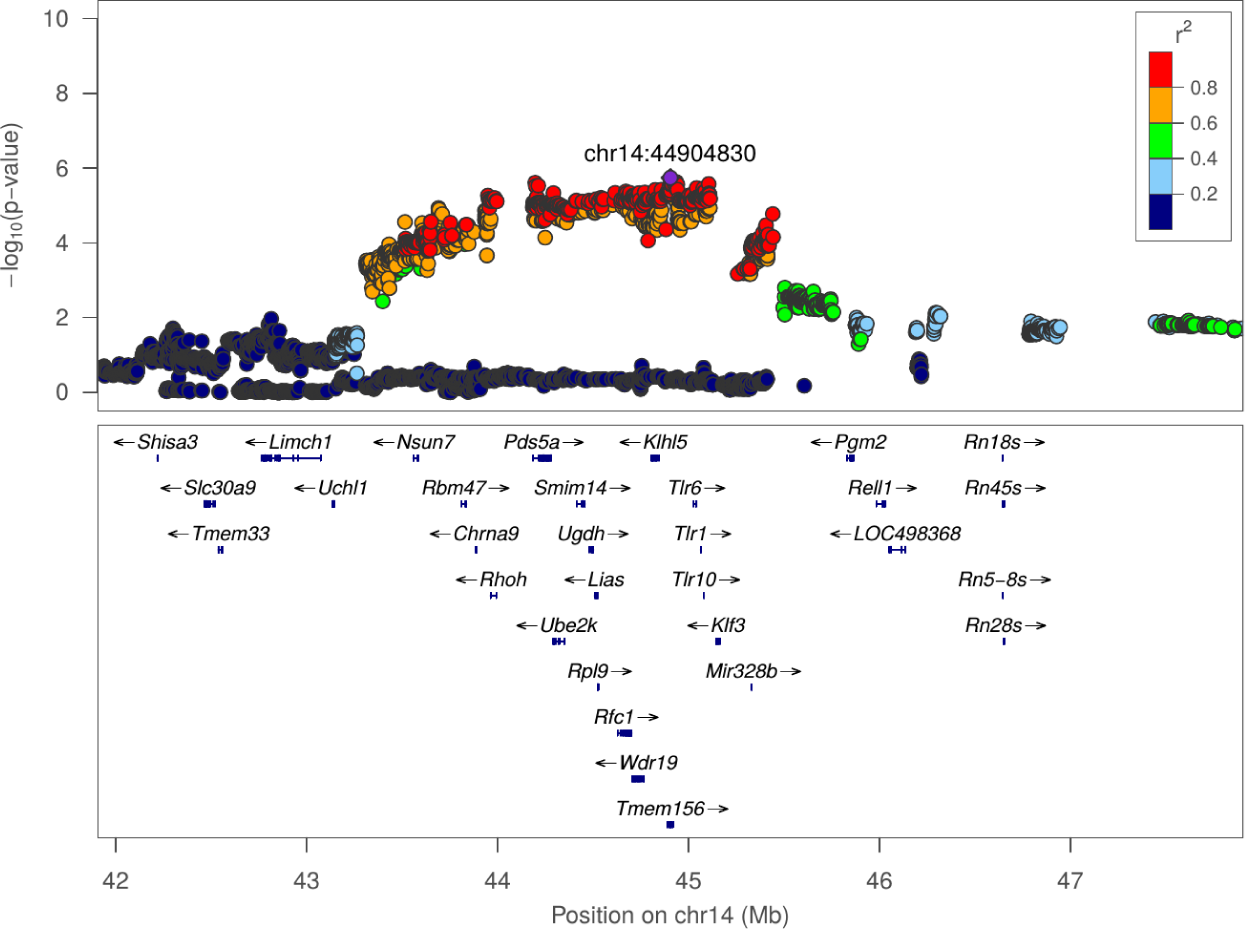
Regional association plot for OFT: Total distance to center zone at chr14:44904830

**Figure S30:**
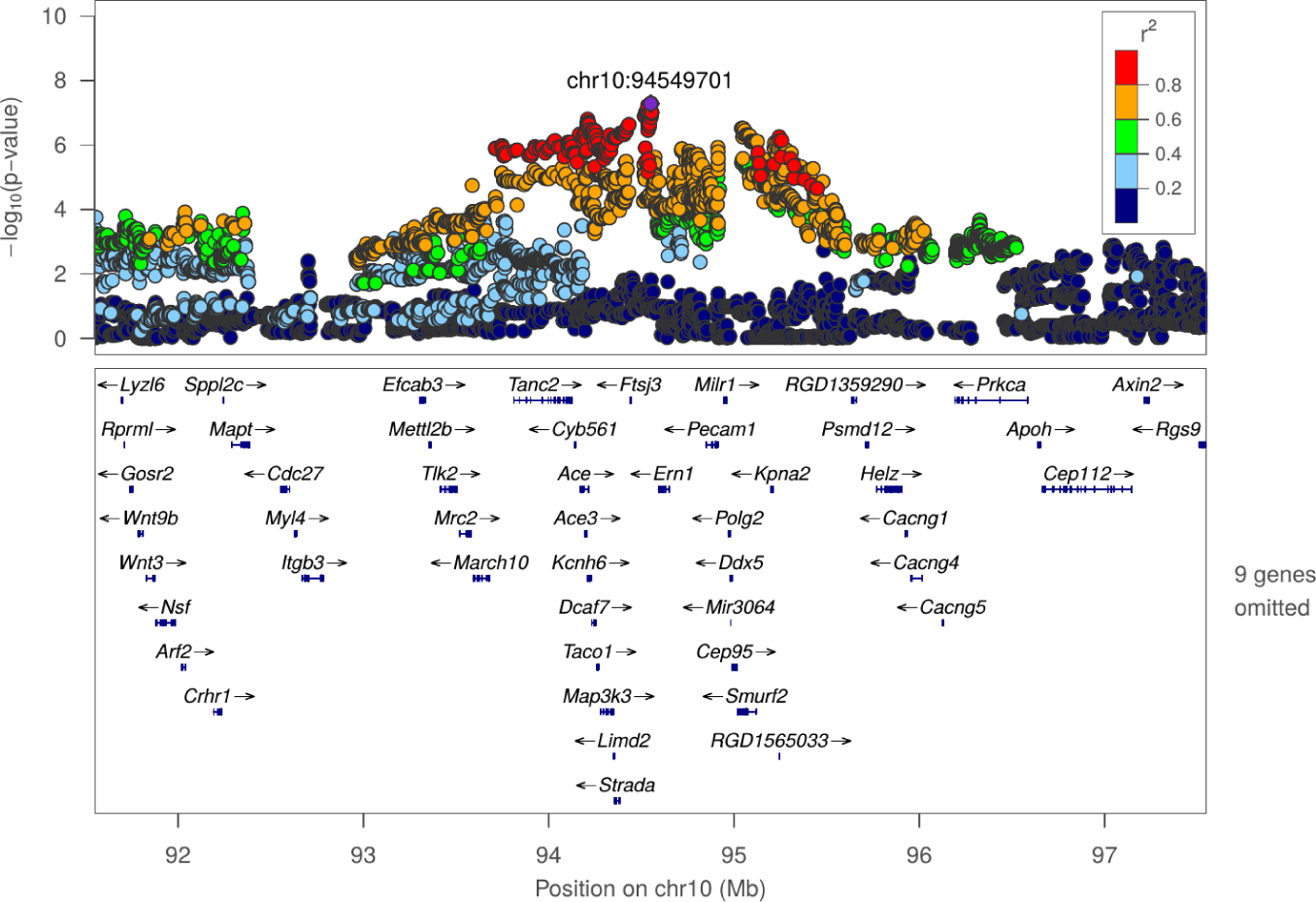
Regional association plot for OFT: Total travel distance at chr10:94549701

**Figure S31:**
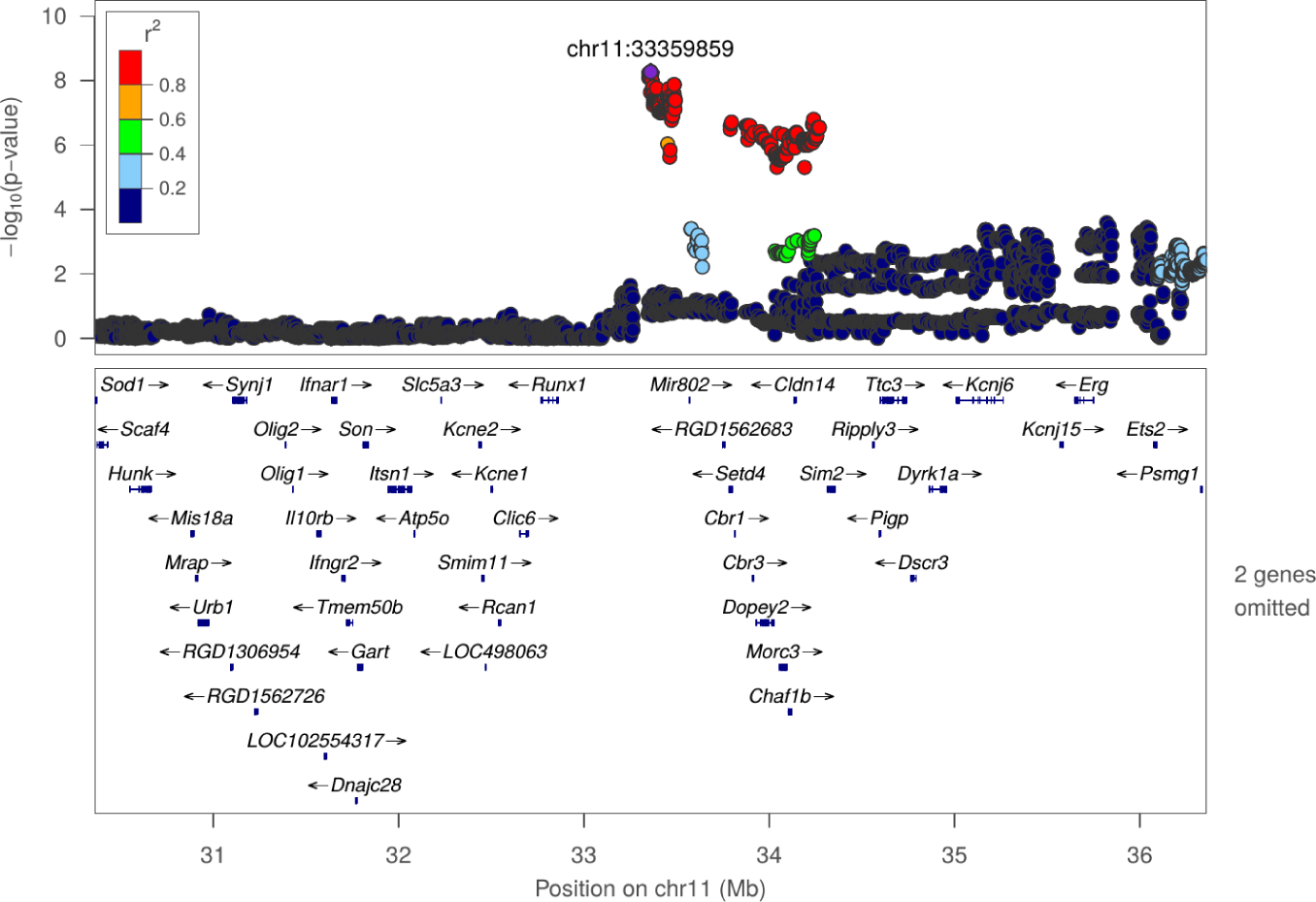
Regional association plot for OFT: Total travel distance at chr11:33359859

**Figure S32:**
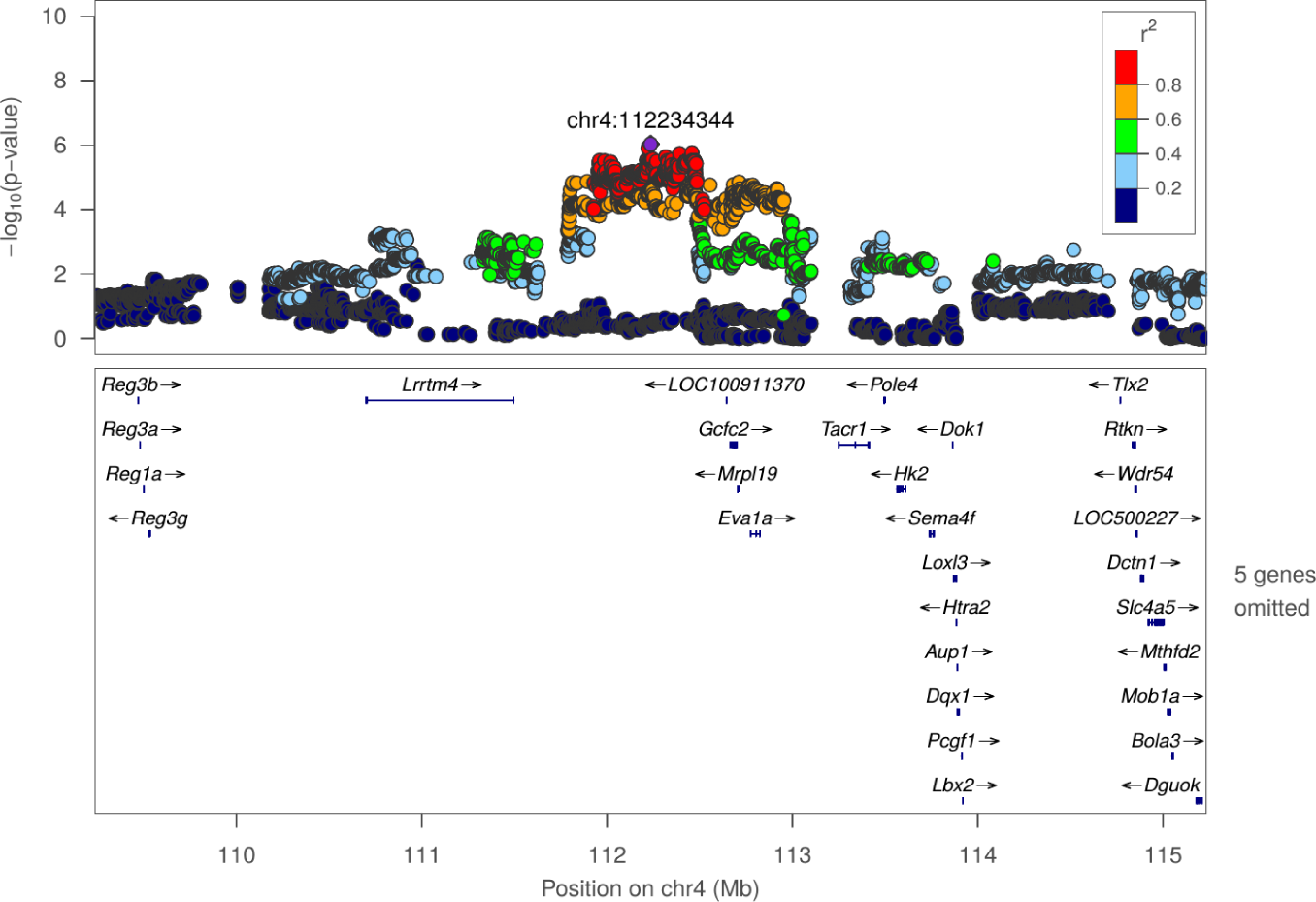
Regional association plot for NOIT: Duration in center zone at chr4:112234344

**Figure S33:**
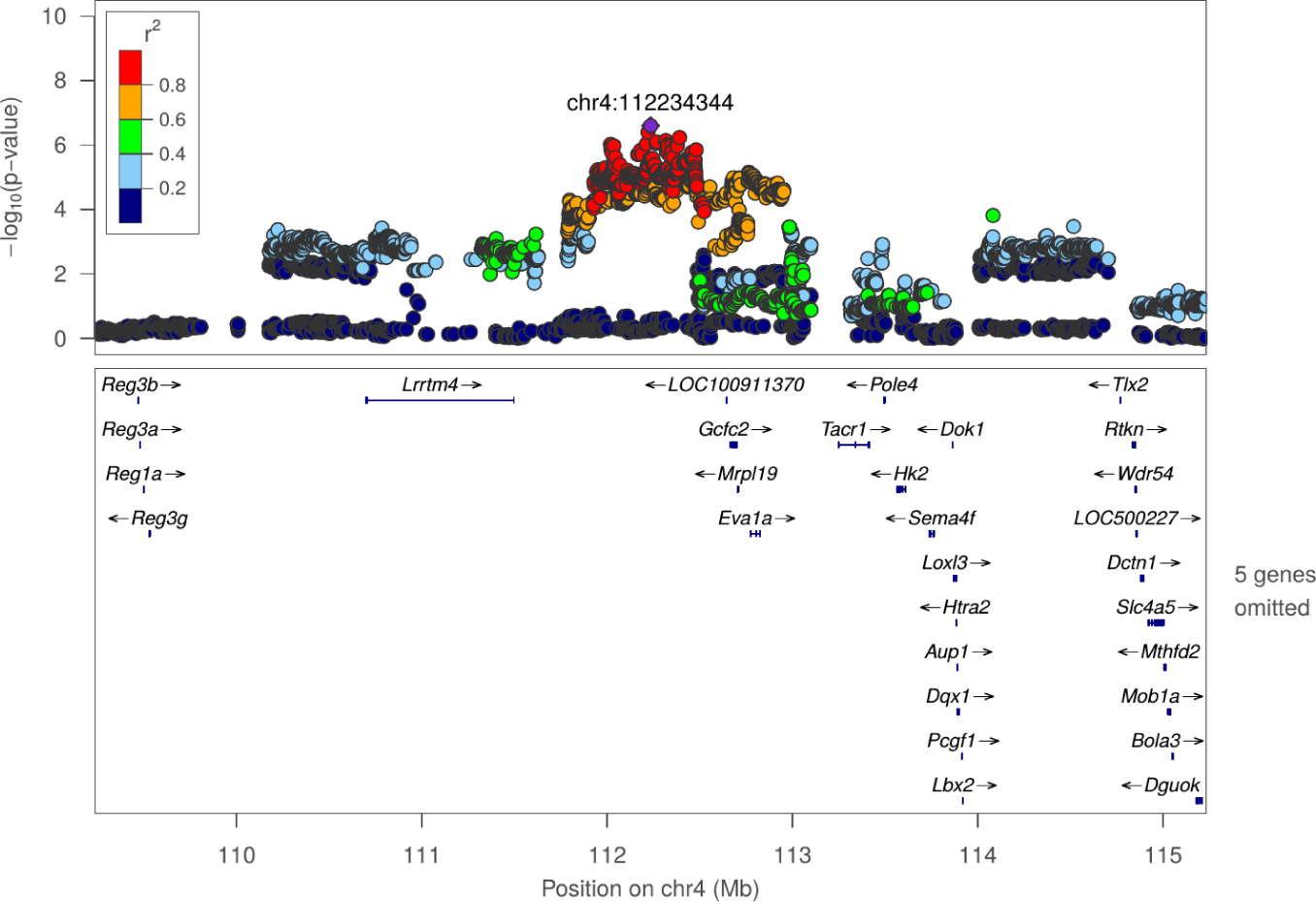
Regional association plot for NOIT: Mean distance to center zone at chr4:112234344

**Figure S34:**
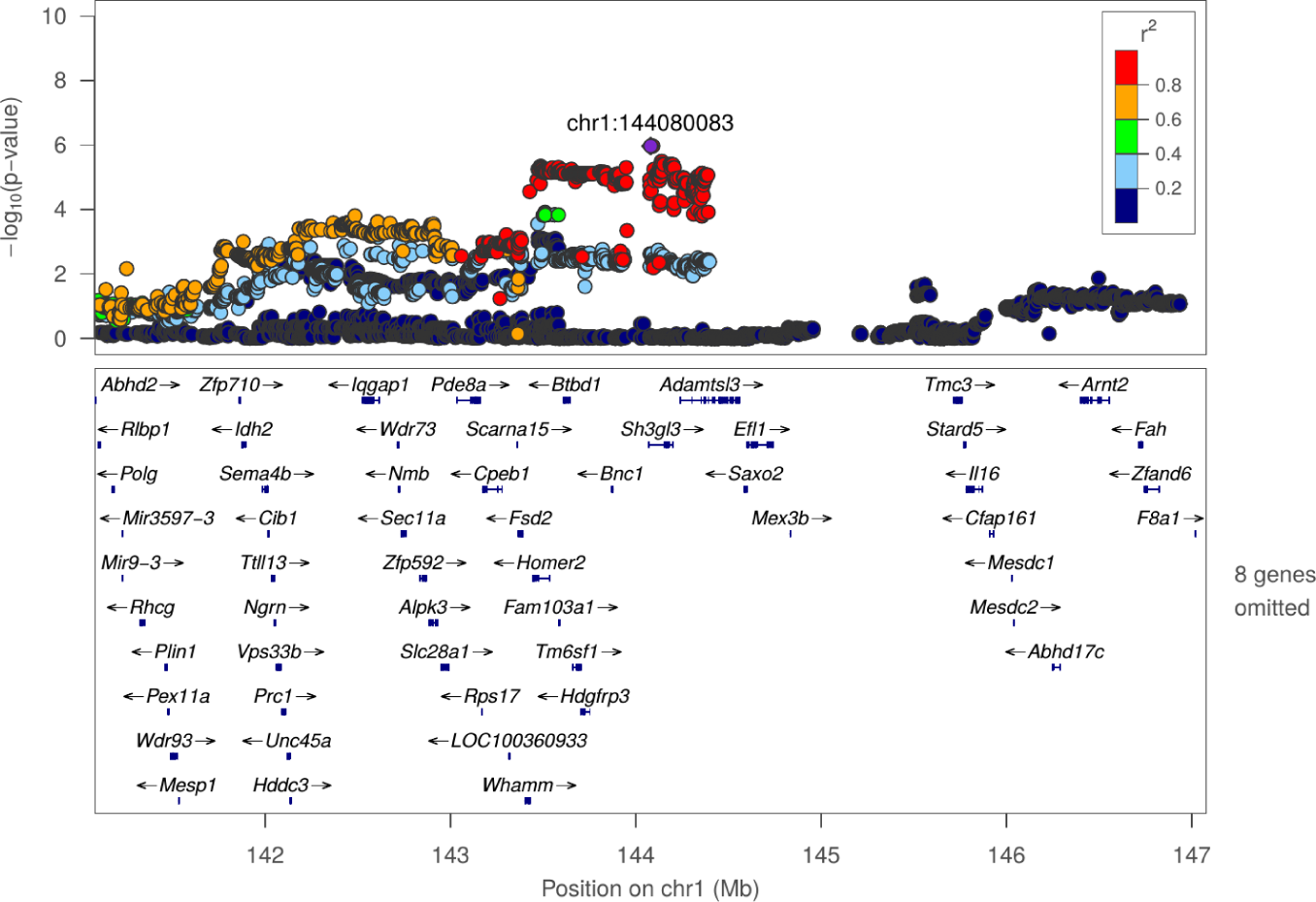
Regional association plot for NOIT: Total distance to center zone at chr1:144080083

**Figure S35:**
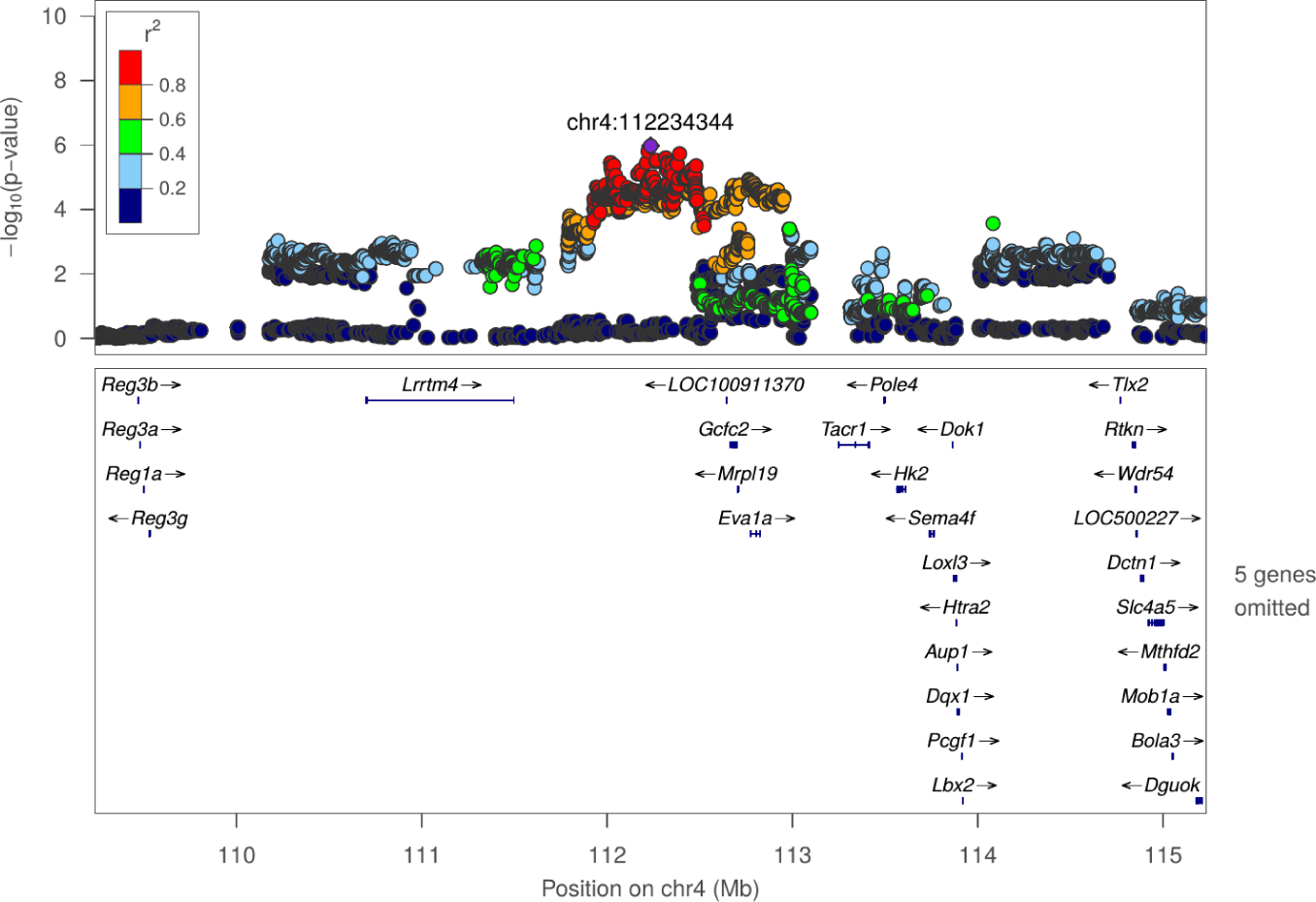
Regional association plot for NOIT: Total distance to center zone at chr4:112234344

**Figure S36:**
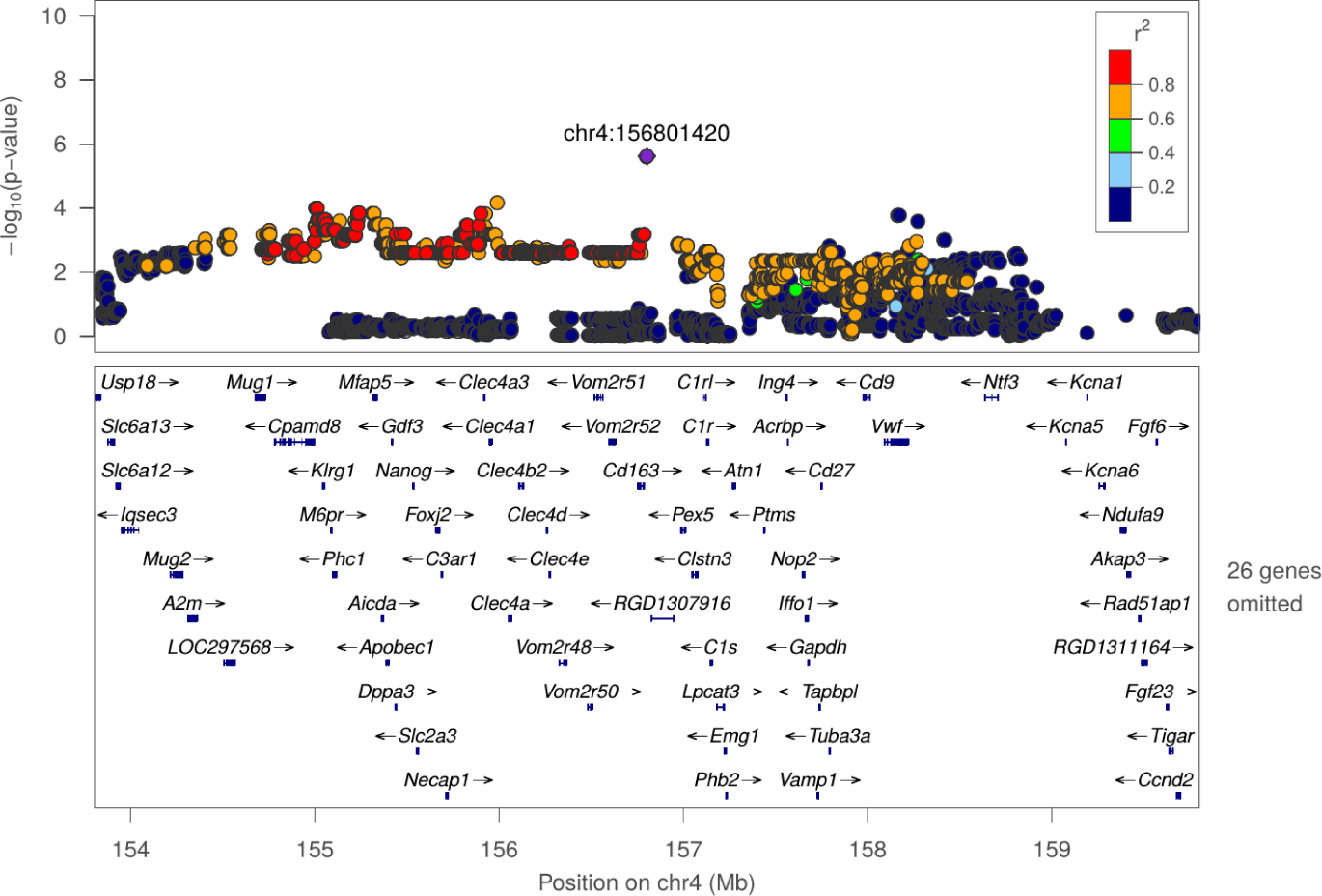
Regional association plot for NOIT: Total distance to center zone at chr4:156801420

**Figure S37:**
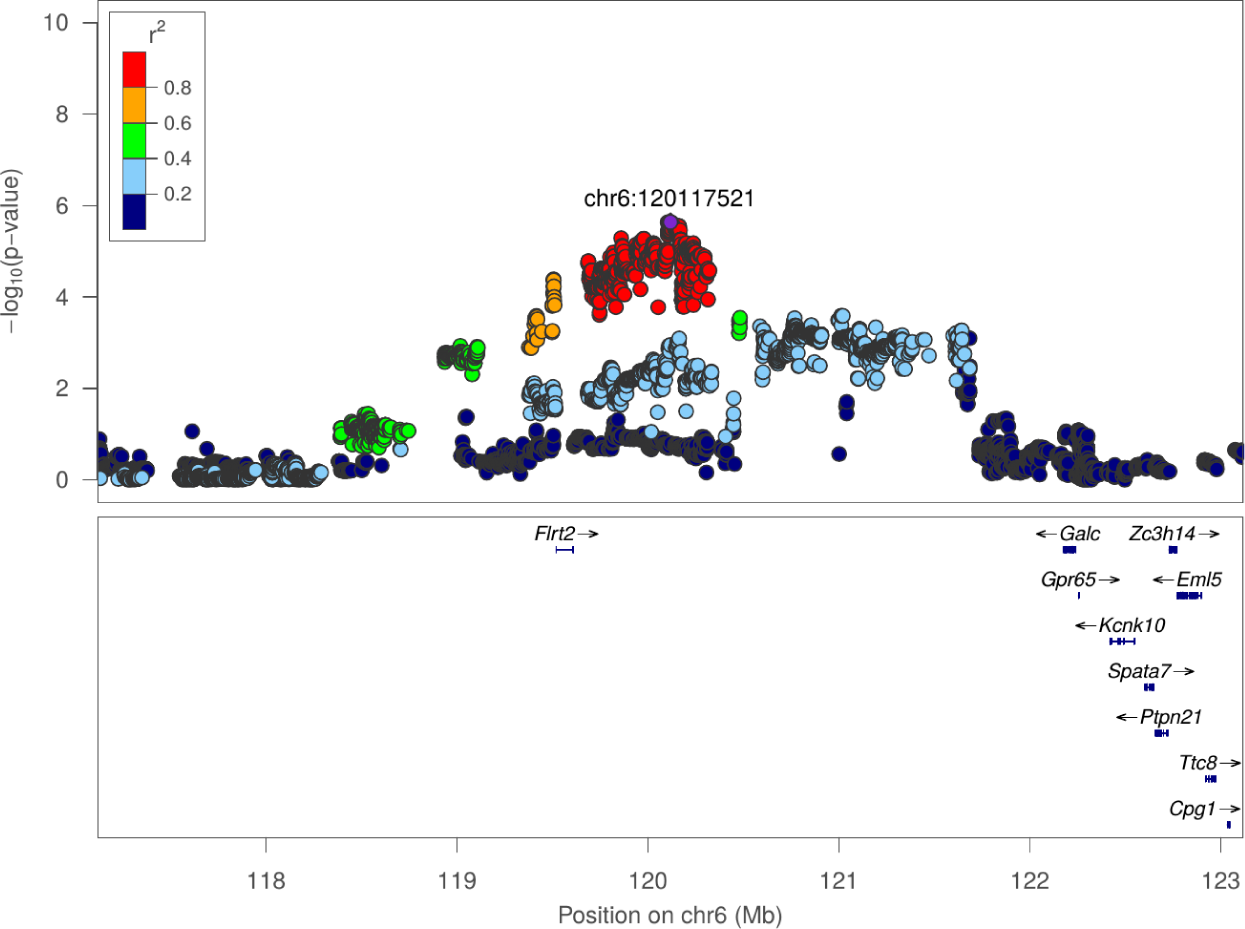
Regional association plot for NOIT: Total travel distance at chr6:120117521

**Figure S38:**
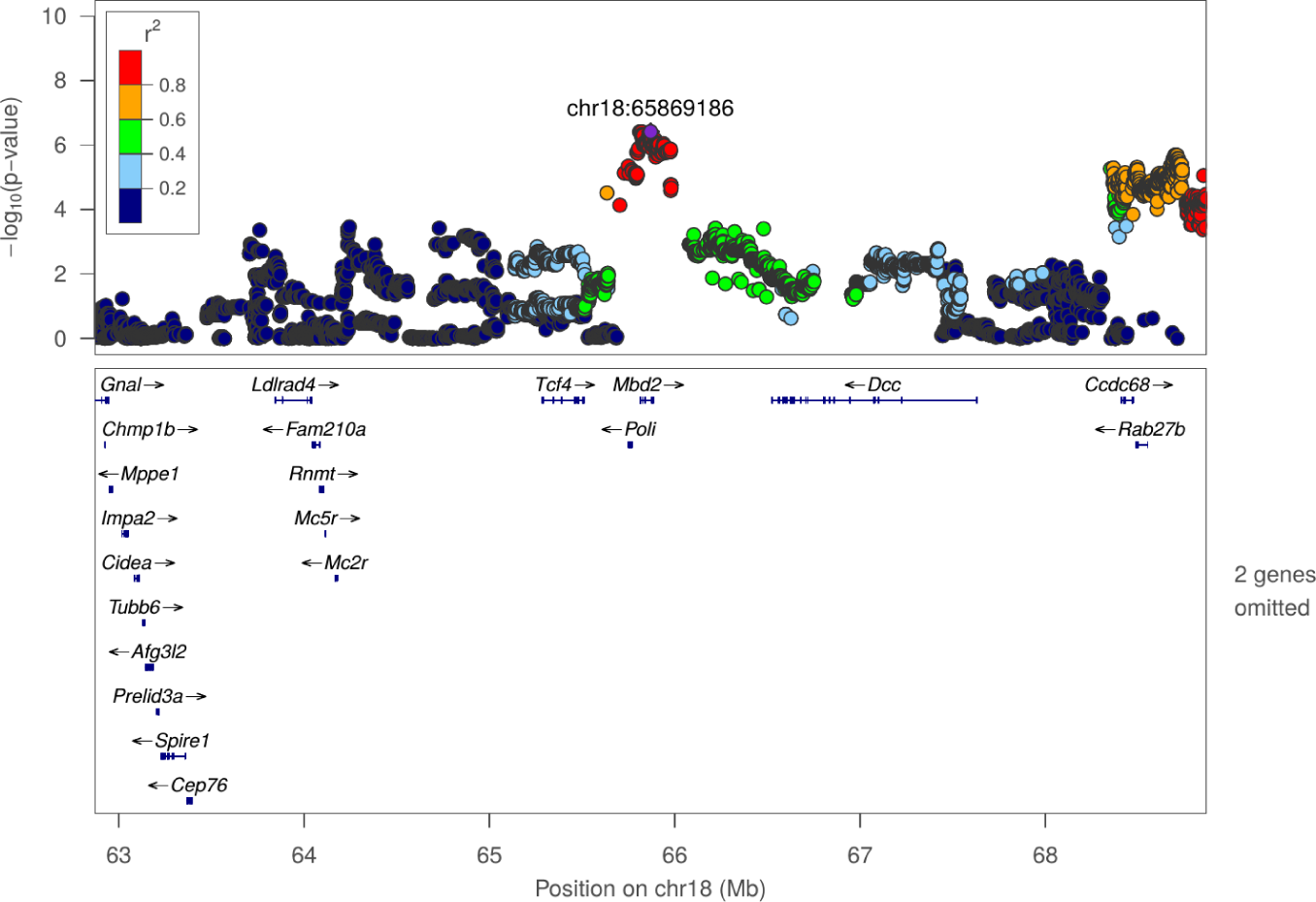
Regional association plot for SIT: Duration in object zone at chr18:65869186

**Figure S39:**
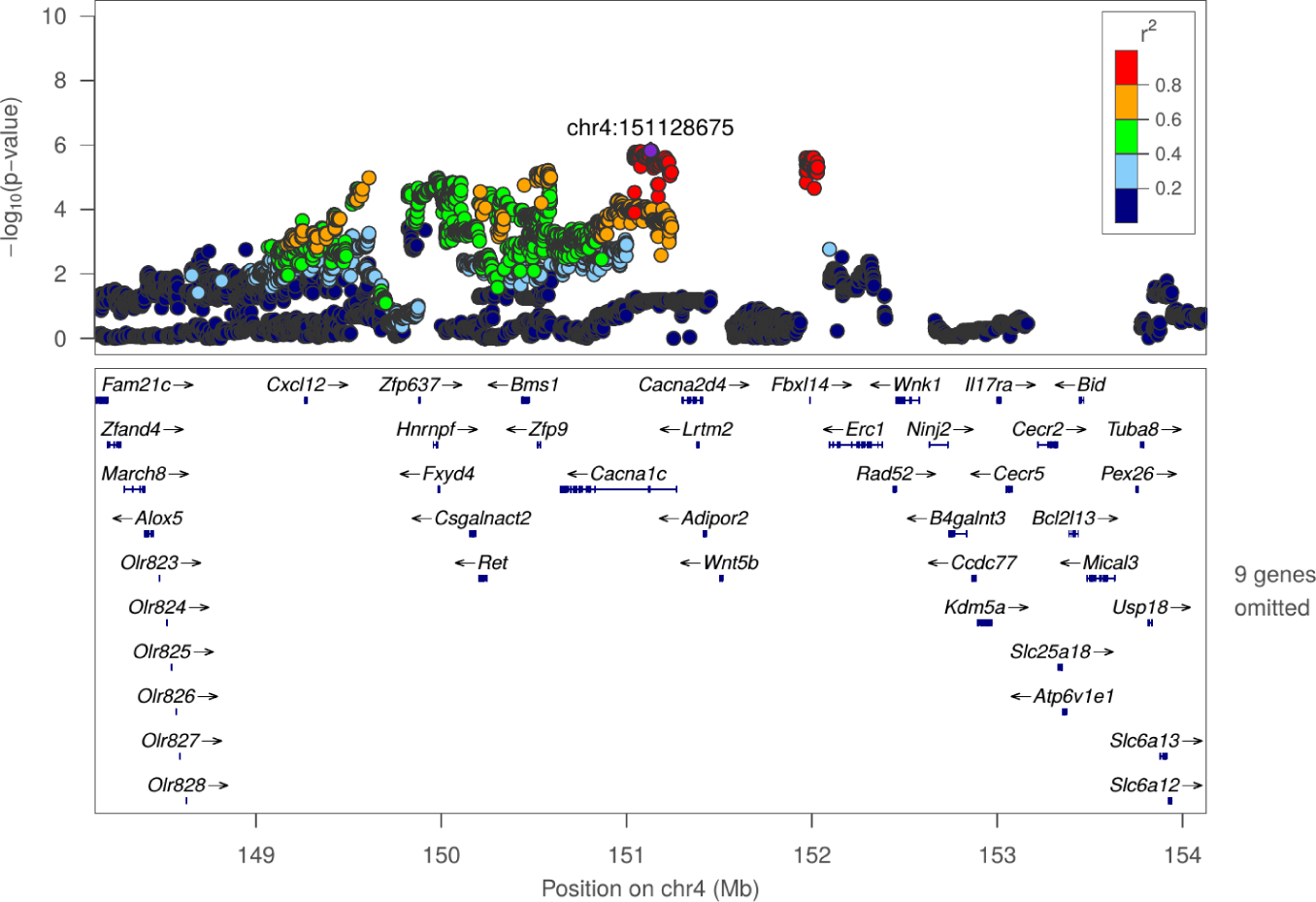
Regional association plot for SIT: Duration in social zone at chr4:151128675

**Figure S40:**
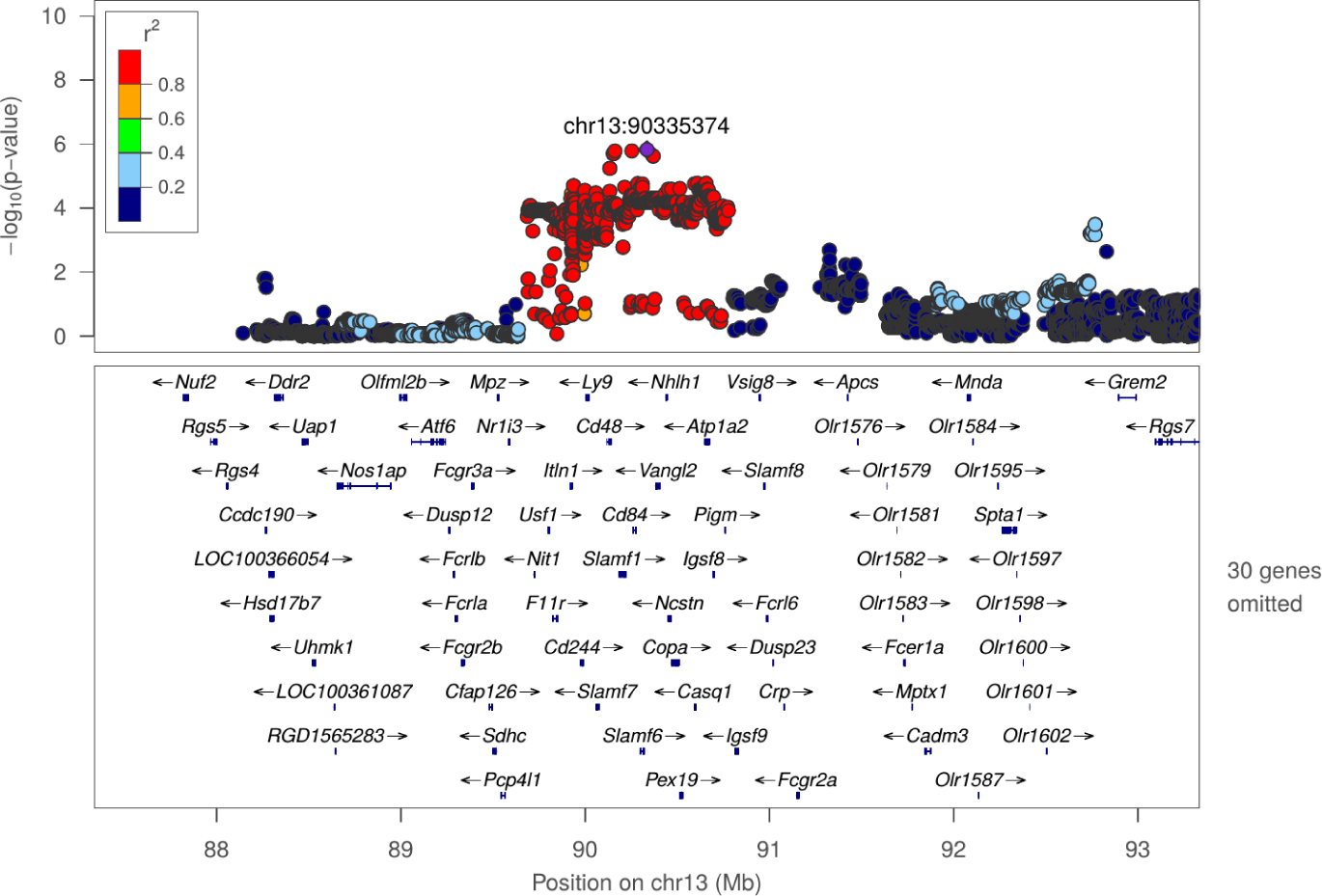
Regional association plot for SIT: Frequency of entering object zone at chr13:90335374

**Figure S41:**
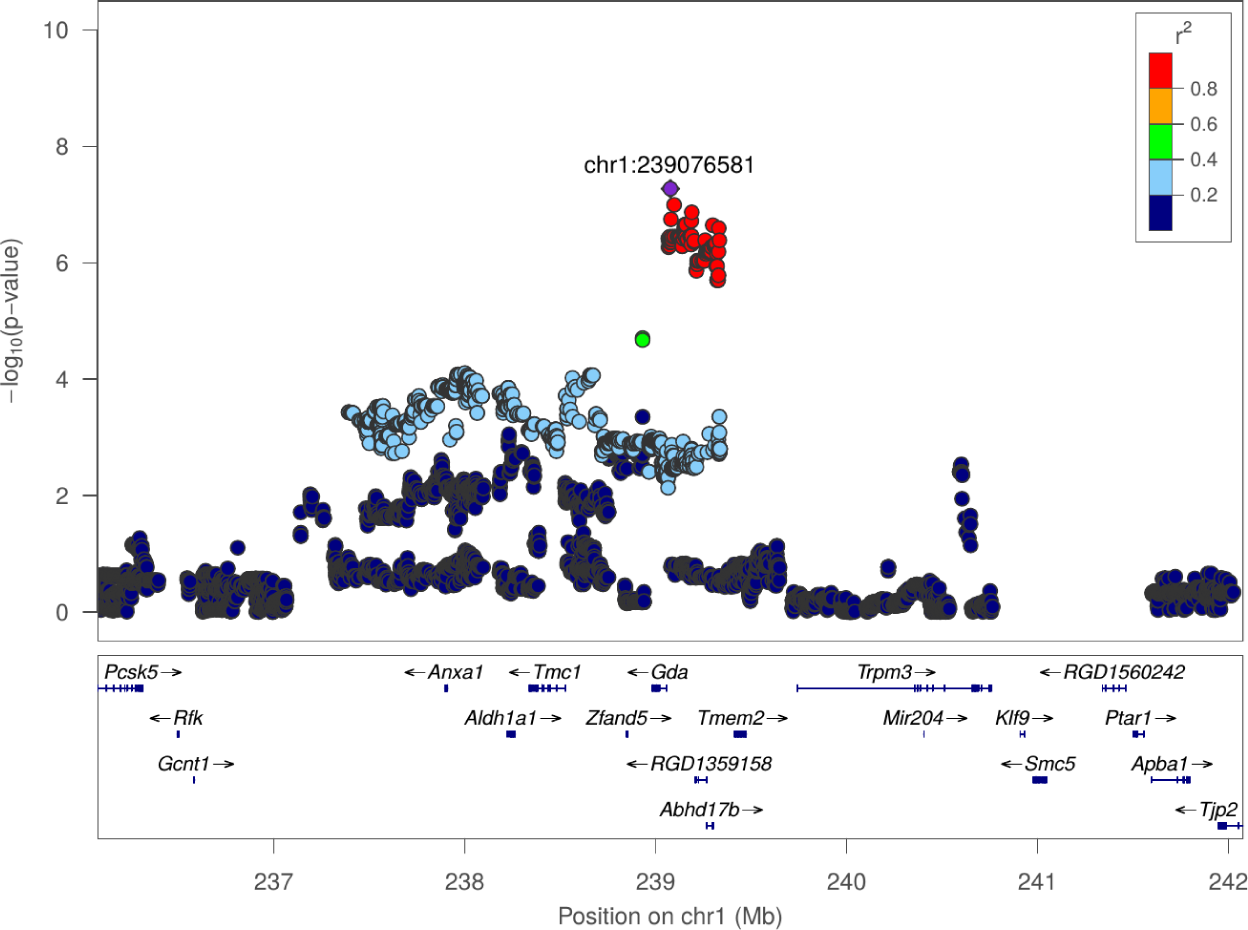
Regional association plot for SIT: Frequency of entering social zone at chr1:239076581

**Figure S42:**
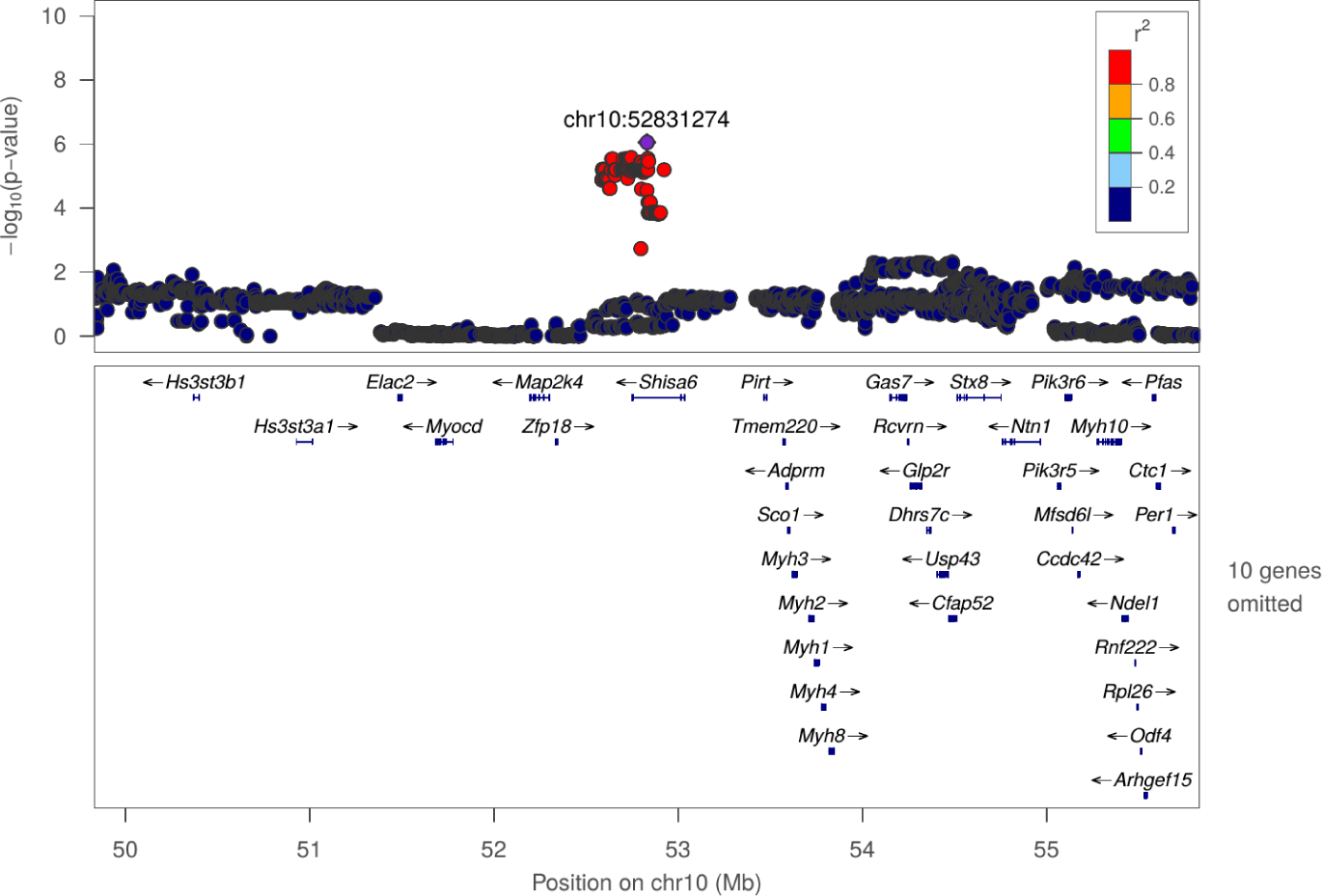
Regional association plot for SIT: Latency of entering social zone at chr10:52831274

**Figure S43:**
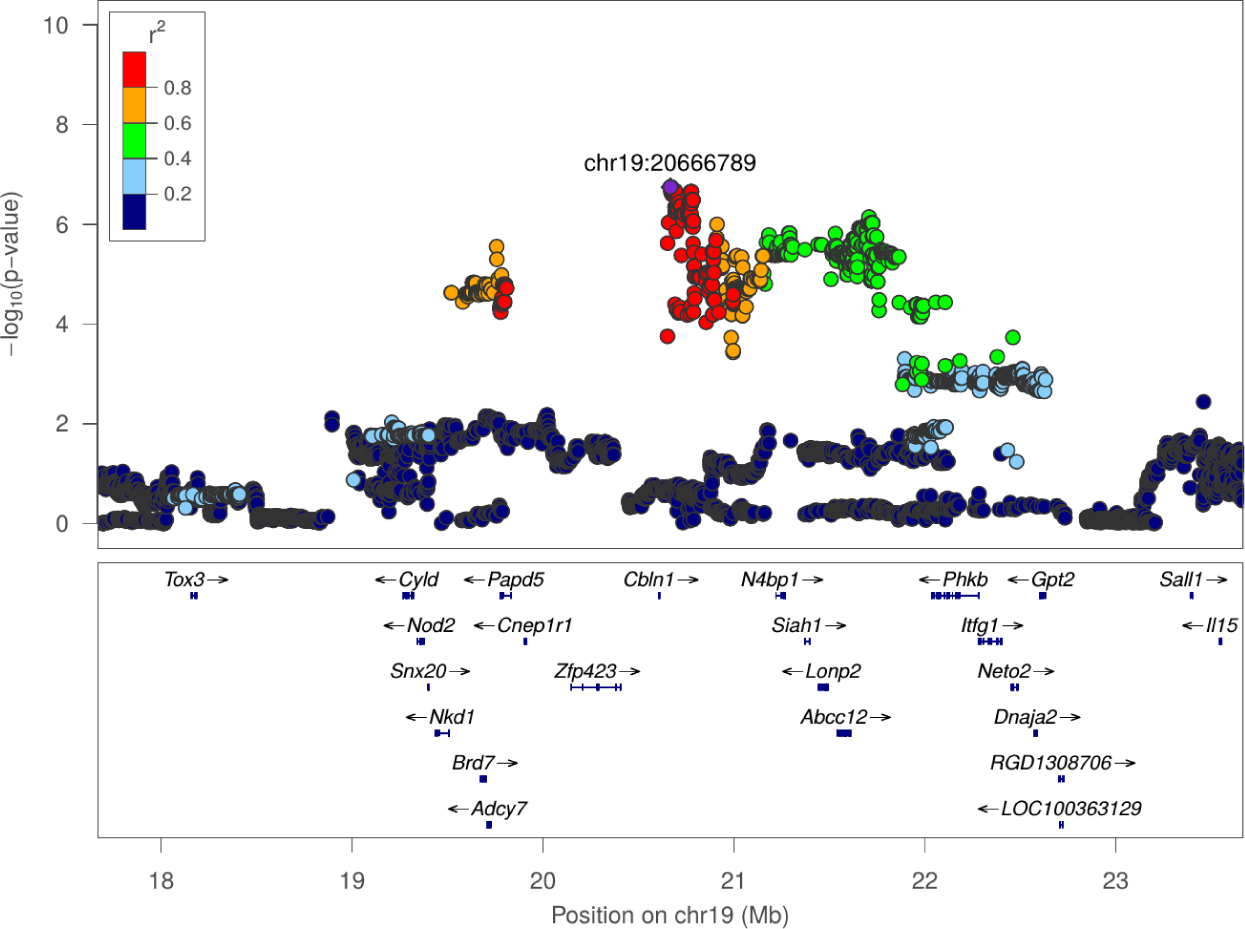
Regional association plot for SIT: Mean distance to object zone at chr19:20666789

**Figure S44:**
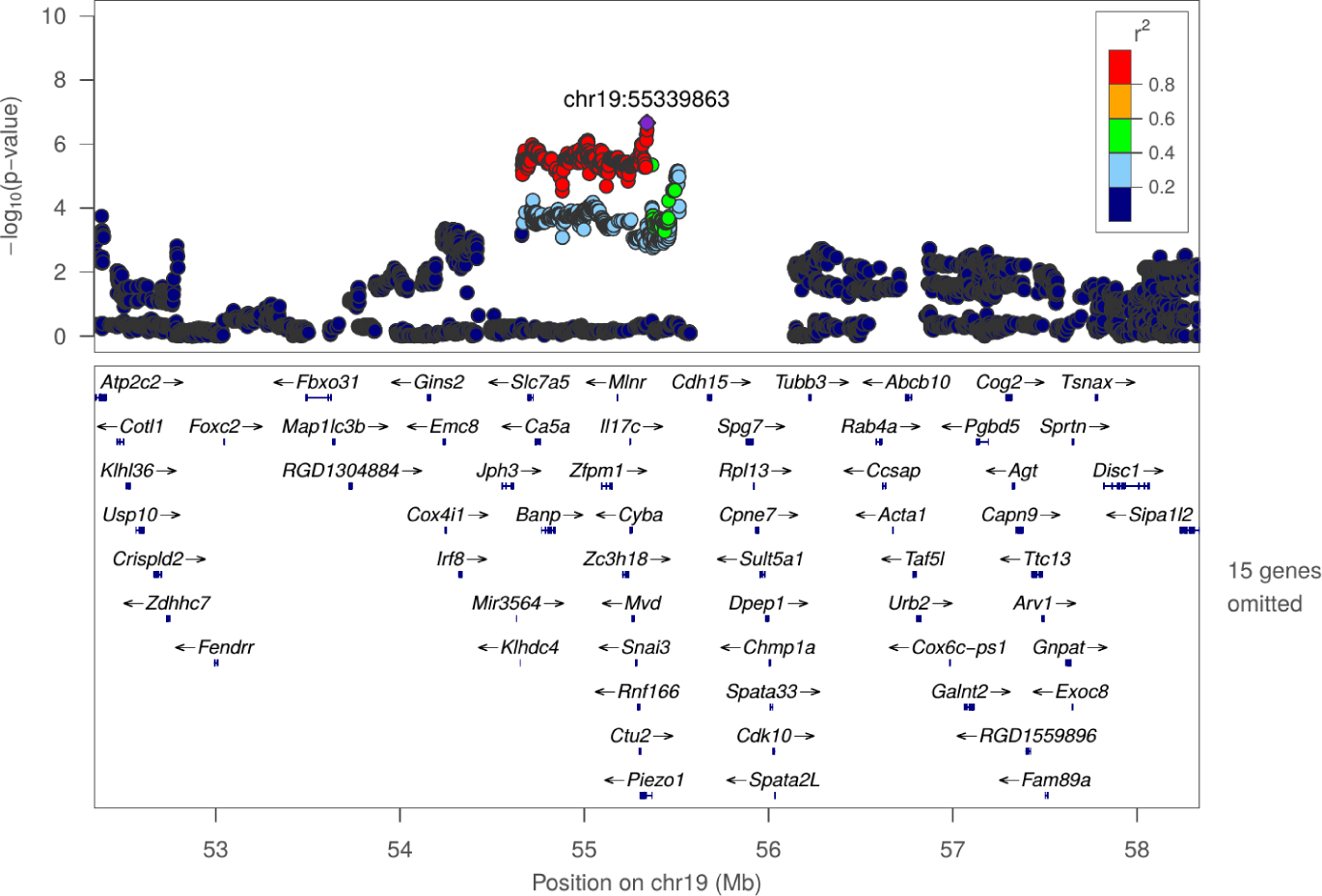
Regional association plot for SIT: Mean distance to social zone at chr19:55339863

**Figure S45:**
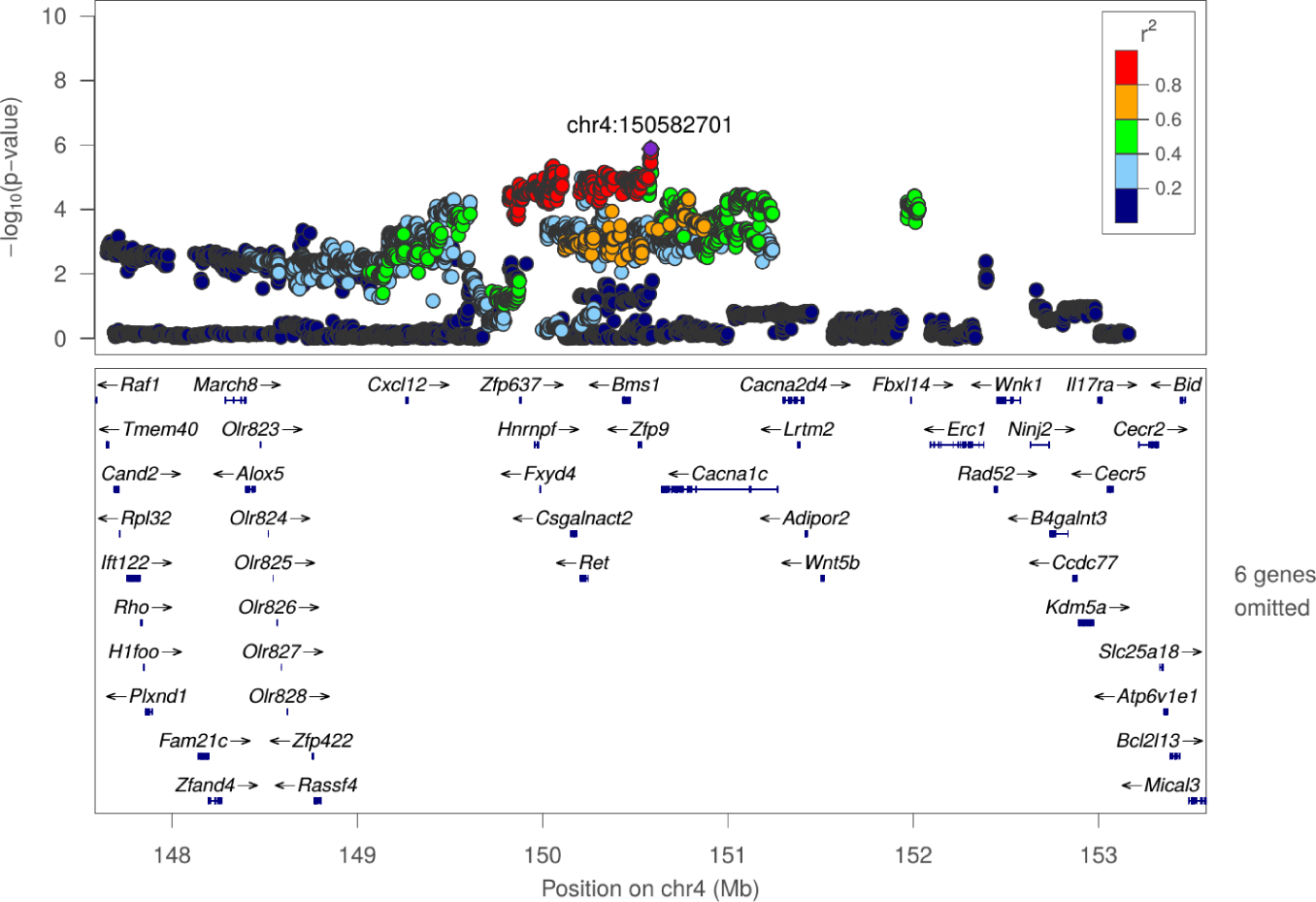
Regional association plot for SIT: Mean distance to social zone at chr4:150582701

**Figure S46:**
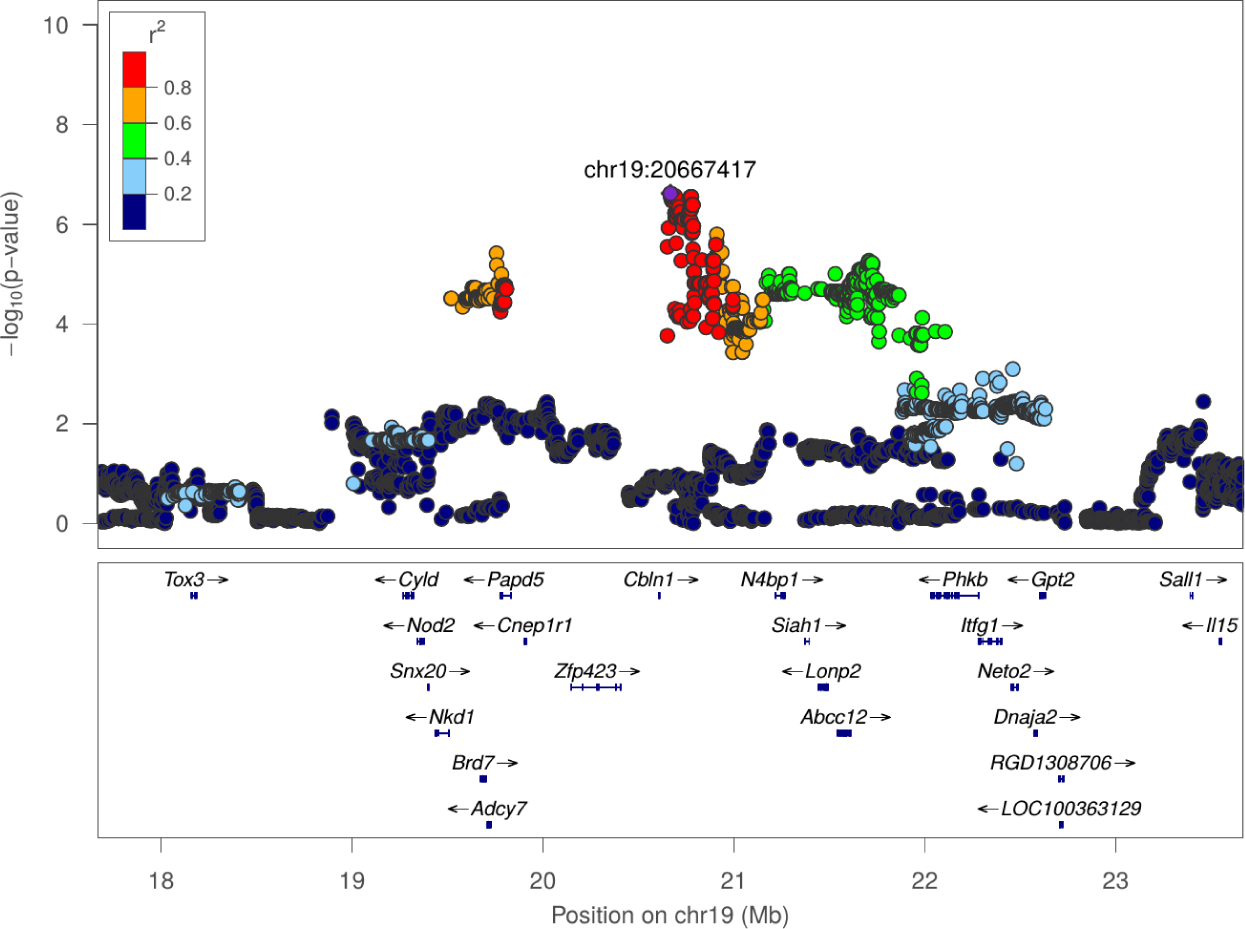
Regional association plot for SIT: Total distance to object zone at chr19:20667417

**Figure S47:**
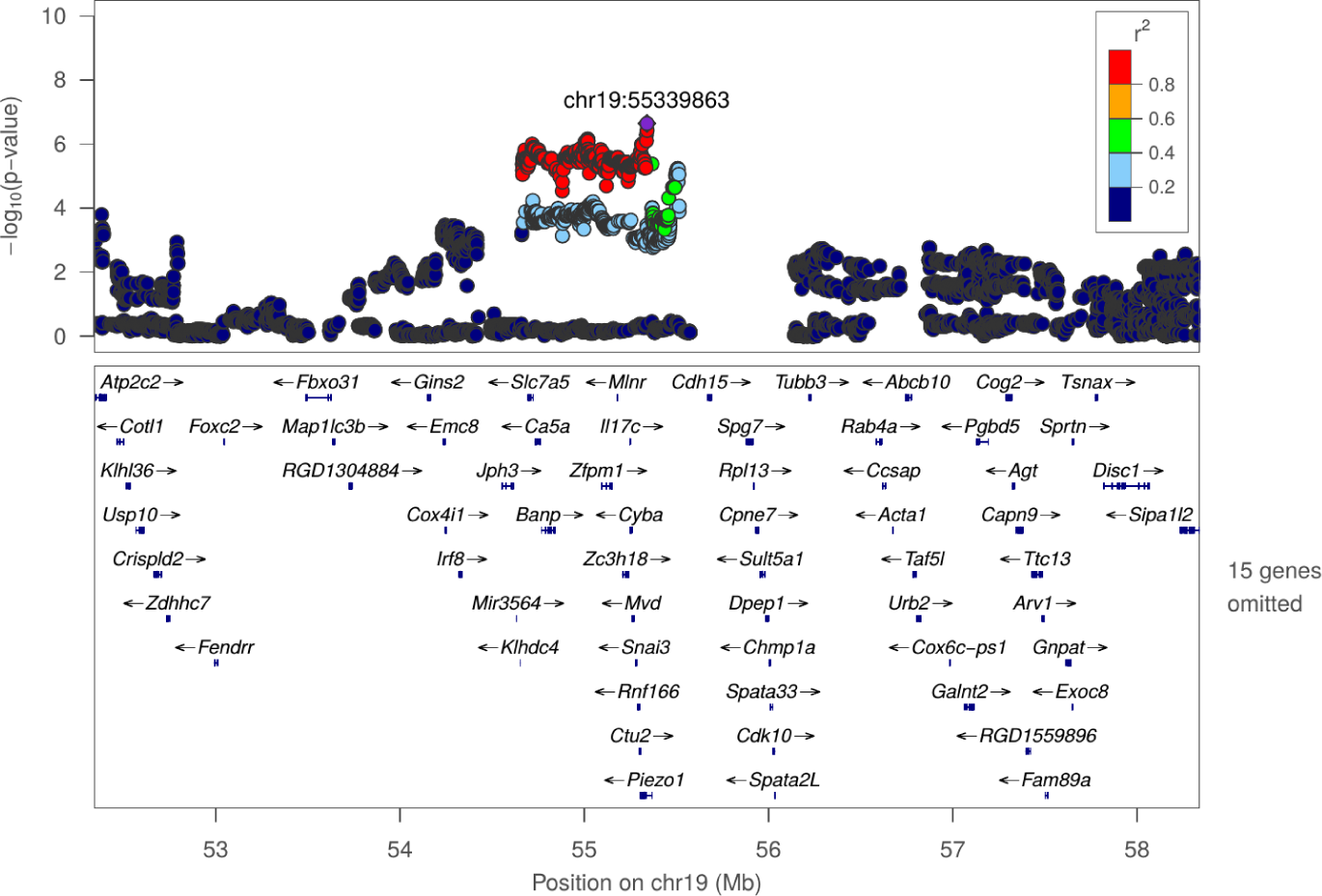
Regional association plot for SIT: Total distance to social zone at chr19:55339863

**Figure S48:**
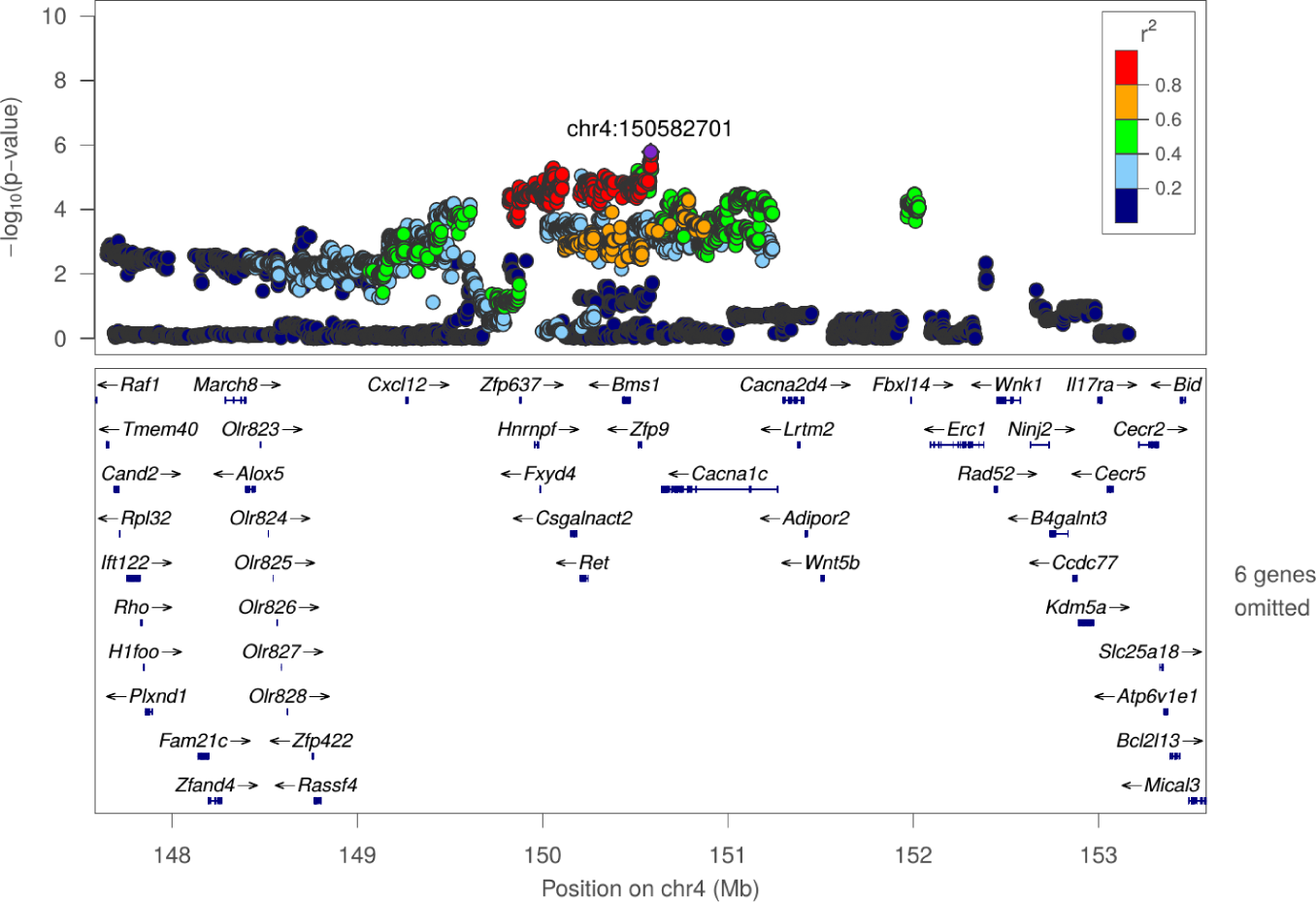
Regional association plot for SIT: Total distance to social zone at chr4:150582701

**Figure S49:**
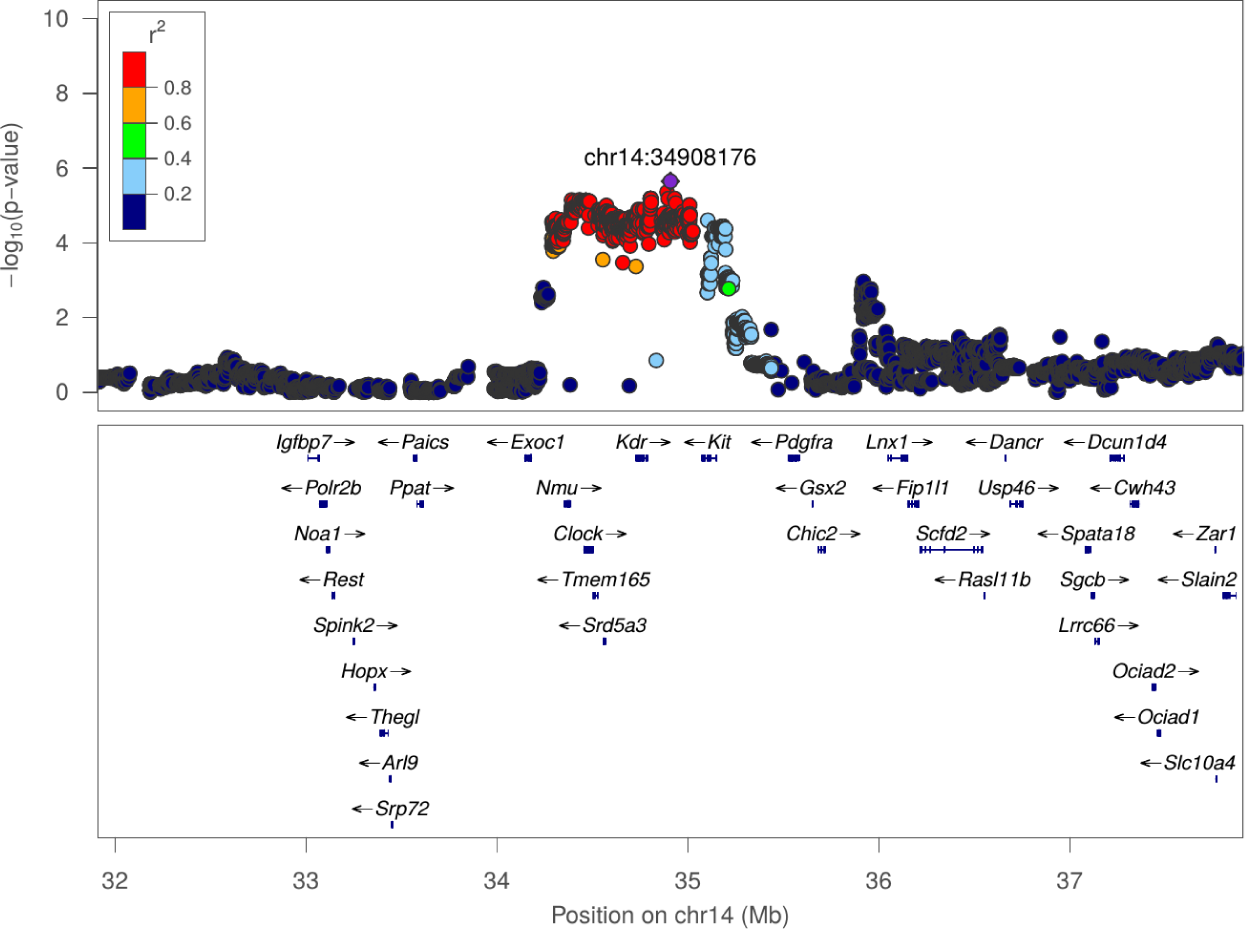
Regional association plot for SIT: Total travel distance at chr14:34908176

**Figure S50:**
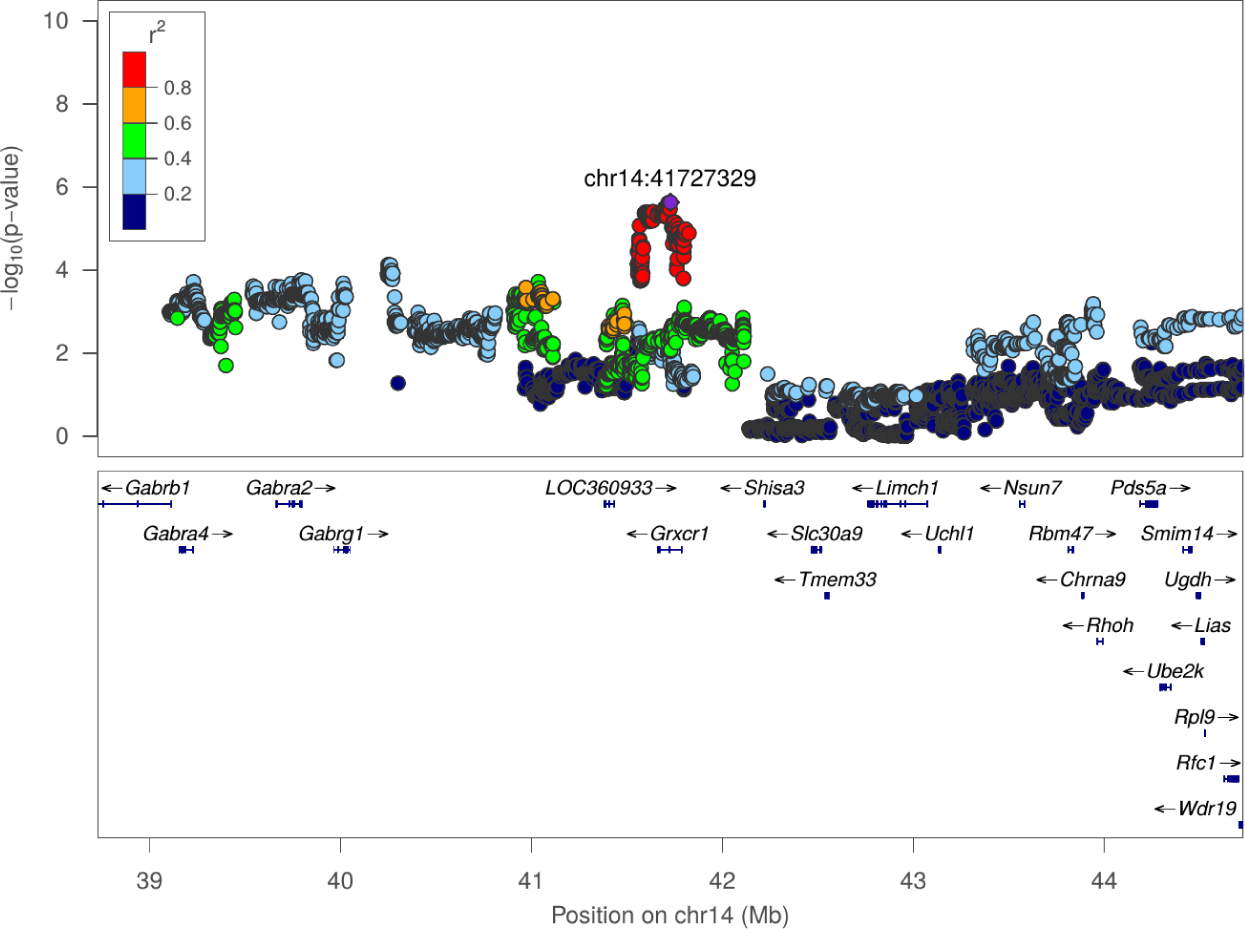
Regional association plot for SIT: Total travel distance at chr14:41727329

